# A unified hierarchical Bayesian approach to transcriptome-wide association study

**DOI:** 10.1101/2024.09.12.612639

**Authors:** Arnab Kumar Khan, Subashani Singh, Tanushree Haldar, Arunabha Majumdar

## Abstract

The transcriptome-wide association study (TWAS) has identified novel gene-trait associations, providing essential biological insights. TWAS combines reference panel transcriptome and genome-wide association study (GWAS) data. Traditional TWAS methods construct a prediction model for gene expression based on the transcriptome data, which is then employed to impute gene expression in the GWAS data. The complex trait in GWAS is regressed on the predicted expression to identify gene-trait associations. Such a two-step approach ignores the uncertainty of the imputed expression and can lead to reduced inference accuracy. We develop a unified Bayesian approach for TWAS that avoids the need for a two-step approach by modeling the two datasets simultaneously. We consider the horseshoe prior to model the relationship between gene expression and local SNPs, and the spike and slab prior to test for an association between the genetic component of expression and the trait to build an integrated Bayesian framework. We extend the method to conducting a multi-ancestry TWAS, focusing on discovering genes that affect the trait in either or both ancestries. Using extensive simulations, we demonstrate that the new approach performs better than existing methods in terms of correctly classifying non-null genes and the accuracy of effect size estimation. To demonstrate our approach, we perform a single and multi-ancestry TWAS for intraocular pressure (IOP), integrating the Geuvadis transcriptome and UK Biobank GWAS data. We find that the identified set of IOP-associated genes is enriched in relevant pathways.

## 1 Introduction

High-throughput genotyping motivated the simple yet powerful design of genome-wide association studies (GWAS), which have revolutionized the mapping of novel genetic variants that modulate complex traits [1]. However, the genetic variants identified by GWAS often reside in the non-coding region of the human genome, making it challenging to establish their functional relations with the trait. We must study intermediate molecular phenotypes and genetic variants together to better understand the biological mechanism through which these variants affect the trait. Gene expression is a crucial molecular phenotype that impacts a complex trait. Regulatory genetic variants, such as expression quantitative trait loci (eQTLs), govern the gene expression level [2] alongside other factors. Frequent overlap between GWAS signals and eQTLs underscores the importance of combining GWAS and gene expression studies to reveal the mechanistic pathway from GWAS-identified susceptibility SNPs to the trait [2]. Transcriptome-wide association study (TWAS) is a powerful approach for detecting novel gene-trait associations that address this objective.

TWAS combines a reference transcriptome and GWAS data to infer the association between a gene and a trait [3]. A reference transcriptome data contains the tissue- or cell-type-specific gene expression levels and genotypes of the gene’s cis/local SNPs, i.e., nearby SNPs, for a set of individuals. The GWAS data consists of trait values and the genotypes for genome-wide SNPs, including the gene’s local SNPs. However, gene expression measurements are not available in the GWAS data. The sample size of the transcriptome data is typically substantially smaller than the GWAS data. We train a regression model using transcriptome data, considering gene expression as the response variable and the genotypes of local SNPs as the predictors. The estimated local SNPs’ effect sizes, therefore, provide a prediction model for gene expression, which we use to predict the genetically regulated component of expression in the GWAS data. Finally, we test for an association between the outcome trait and the predicted genetically regulated component of expression (GReX) in the GWAS data. TWAS has a few advantages over GWAS. First, TWAS is a gene-level test with less multiple-testing burden. Second, TWAS bridges the functional gap in GWAS by incorporating gene expression as a potential causal mediator between SNPs and the outcome. However, the better performance of TWAS compared to GWAS in discovering novel genetic loci associated with an outcome trait crucially depends on the local heritability of gene expression, i.e., the proportion of variance of expression explained by the local SNPs. TWAS performs better than GWAS when the local heritability is higher [4].

Most TWAS methods [5, 6] employ the two-step strategy. For example, penalized regression, such as Lasso [7], is applied to the transcriptome data to obtain a prediction model for expression. A linear or logistic regression is applied to regress the outcome on the predicted GReX in the GWAS data. Such an approach disregards the uncertainty of the imputed GReX in the second step, which arises from the variance of the estimated effect size vector of local SNPs in the transcriptome data. The lack of tuning of the noise in the predicted expression can reduce the overall accuracy of TWAS, e.g., inflation of type I or II error rates [8] or less precise estimation of the gene’s effect size. A popular TWAS method, PrediXcan [3], employs the Elastic net penalized regression [9] to model the relationship between gene expression and local SNPs. A Bayesian method, BGW-TWAS [10], employed the Bayesian variable selection regression based on the spike and slab prior to model the eQTL effects. However, it also follows a two-step strategy without accounting for the uncertainty of the predicted expression. Another method, TIGAR [11], which employs Bayesian regression based on the Dirichlet process in the first step, suffers from the same limitation. These approaches finally sorted to obtain the p-values of gene-trait association, even though BGW-TWAS and TIGAR employed Bayesian regressions in the first stage of TWAS. Instead of a two-step strategy, an integrated statistical framework that jointly models the two datasets simultaneously can overcome this limitation.

We develop a unified Bayesian approach to conduct TWAS, jointly modeling the transcriptome and GWAS data. Bayesian regularization methods consider various shrinkage priors. Global local shrinkage priors are an important class of such priors that shrink the noisy small effect sizes but leave the larger effect sizes unreduced. To leverage the advantage of such efficient regularization approaches, we employ the horseshoe prior [12], a prominent choice of global-local shrinkage prior, when building the relationship between the gene expression and local SNPs in the transcriptome data. We utilize the spike and slab prior [13] while testing for an association between the outcome and GReX in the GWAS data under the joint framework. The integrated Bayesian approach surpasses the impromptu genre of the two-step approaches. A recent TWAS method, PMR-EGGER [14], accounts for the uncertainty of the predicted expression. However, it relies on the polygenic assumption that all local SNPs around the gene have a non-null effect on the expression. However, we expect the eQTL effects underlying a gene’s expression to be sparse [15]. In contrast, our approach avails the benefits of well-founded priors to model the sparsity in the TWAS data and performs a comprehensive Bayesian inference on the gene-trait association.

The horseshoe prior (HSP) and the double-exponential prior (DEP) corresponding to Lasso [7] both shrink the small effects. However, the former refrains from shrinking the sizeable effects [16]. The DEP reduces the large effect sizes by a non-vanishing amount. In the context of TWAS, we do not require explicit variable selection as done by Lasso or a two-group spike and slab prior when modeling the effects of local SNPs on gene expression. Because eQTL discovery is not the goal here. Hence, we prioritize a continuous shrinkage by the HSP to robustly model various types of sparsity present in the transcriptome data. We implement the classical spike and slab prior (SSP) with a point mass at zero to test for a gene-trait association explicitly [13].

We extend our model to include multiple ancestries, thereby enhancing genetic diversity in the TWAS analysis. TWAS requires the transcriptome and GWAS data to be ancestry-matched. The gene expression prediction model derived for one ancestry is not portable to another ancestry due to differences in eQTL effects, LD structure of the local SNPs for a gene, and other factors. Transcriptome and GWAS data from diverse ancestries, such as Europeans and Africans, are currently available. We aim to jointly analyze datasets from diverse ancestries to leverage ancestry-shared and ancestry-specific genetic bases of a complex trait, thereby improving the genetic diversity and effectiveness of TWAS analyses.

A multi-ancestry TWAS fine-mapping approach, MA-FOCUS [17], analyzes cross-ancestry TWAS data to prioritize the causal genes in a genetic region associated with the trait. However, it adopted the traditional two-step TWAS procedure and did not calibrate the uncertainty of the imputed expressions. It assumes homogeneity in the effect sizes for the causal genes across ancestries, which may not be true for many genes. Another approach, METRO [18], proposed a unified approach to multi-ancestry TWAS while considering a joint likelihood-based framework to model the transcriptome and GWAS data together. The method considered independent normal distributions for the regression coefficients in the linear model of the gene expression, treating local SNPs’ genotypes as covariates in the transcriptome data, which is analogous to a penalty function considered in ridge regression. Ridge regression applies the same amount of shrinkage to all the local SNPs’ effect sizes. On the contrary, the HSP underlying our approach differentially shrinks the effect sizes, making it a more effective approach to regularization. Although METRO considers different expression data for each ancestry, it utilizes a single GWAS dataset for all ancestries. Our approach takes into account different GWASs, each with a distinct ancestry. Likewise, in our single-ancestry setup, we combine the HSP and the SSP to form our multi-ancestry TWAS model, focusing on discovering the genes associated with the trait in either ancestry. We refer to our method as **glossTWAS** (**g**lobal-**lo**cal **s**hrinkage and **s**pike and **s**lab prior-based **TWAS**).

We conduct extensive simulation studies to assess the performance of glossTWAS in single-ancestry and multi-ancestry TWAS settings. We compare its performance with prominent existing approaches in terms of correctly classifying null and non-null genes, as well as the accuracy of effect size estimation. We apply glossTWAS to identify genes associated with the intraocular pressure (IOP). We integrate the UK Biobank GWAS data with transcriptome data from the GEUVADIS study. We perform single-ancestry TWAS for Europeans and multi-ancestry TWAS for European and African ancestries.

## 2 Results

### 2.1 Overview of methods

GlossTWAS is a unified Bayesian approach that models the uncertainty of the predicted expression simultaneously while testing for the gene-trait association. The approach provides estimates of the gene-trait effect sizes. We develop our model based on the horseshoe prior (HSP) and spike and slab prior (SSP) in a joint framework. We consider the HSP to model the relationship between gene expression and the local SNPs surrounding the gene. We implement the SSP to test for the gene-trait association. We employ the Metropolis with Gibbs sampling algorithm to perform the Markov Chain Monte Carlo (MCMC) and generate posterior samples of the model parameters. We calculate the Bayes factor and posterior probabilities for testing the association between a gene and the trait. Next, we extend the Bayesian model to perform TWAS for two ancestries. In a hierarchical Bayesian setup, we consider two HSPs for the local SNPs’ effects on the gene expression in the two ancestries. They share the same global and local shrinkage parameters. We can similarly extend the hierarchical approach for more than two ancestries. We describe the details of the models in the Materials and Methods section, and the MCMC algorithms in the supplementary materials. In multi-ancestry TWAS, the null hypothesis implies that the gene has no effect on the outcome trait in both ancestries. The alternative hypothesis implies a non-null effect of the gene on the outcome in either ancestry, or both ancestries in the same direction. However, if we find a gene to be associated, it can be due to substantial effects in both ancestries, moderate effects in both, or prominent in one but negligible in the other. We provide a diagram in the methods section (Figure 12) that outlines the main hierarchical Bayesian structure of the multi-ancestry glossTWAS.

### 2.2 Simulation results

We compare the performance of our Bayesian approach, glossTWAS, with several prominent TWAS methods using extensive simulations. We regard the accuracy of effect size estimation and the correct classification of a non-null gene as the essential metrics to evaluate a method comprehensively. The details of the simulation procedure are described in the Materials and Methods section.

#### 2.2.1 Single-ancestry TWAS

We first discuss a summary of results obtained by glossTWAS, while estimating the effect of GReX on the trait (*β*) (Table 1) for a single-ancestry TWAS. The estimates based on the posterior mean and median of *β* are similar for varying choices of the effect size. For example, when *β* = 0.2, the posterior mean and median are 0.2 and 0.19 respectively, with posterior standard deviation (SD) 0.07 (Table 1). In this case, the 95% central posterior interval (CPI) is (0.09, 0.33), which contains the true effect size. GlossTWAS performs robustly both for positive and negative effect sizes. For *β* = −0.2, the posterior mean and median are both −0.19 with the 95% CPI is (−0.32, −0.08). The width of the CPI and the magnitude of the posterior SD increase marginally as the absolute value of *β* increases (Table 1). The Bayes factor (BF) and posterior probability of null association (PPNA) obtained by glossTWAS measure the strength of gene-trait association. The empirical average of log_10_(BF) values obtained across repetitions in a simulation scenario increases steadily with the absolute value of *β*. As expected, the corresponding PPNA values decrease (Table 1). In the absence of any association with *β* = 0, the mean log_10_(BF) value is negative, −0.7, and we obtain the mean values as 1.6, 5, 11.9, 23.7 for *β* = 0.1, 0.2, 0.3, 0.4, respectively. Thus, the log_10_(BF) values increase with the strength of association. The mean PPNA value for *β* = 0 is 0.97 and it gradually decreases to 0.31, 0.04, 8 *×* 10^−4^, 1.5 *×* 10^−8^ for *β* = 0.1, 0.2, 0.3, 0.4, respectively (Table 1).

**Table 1.**
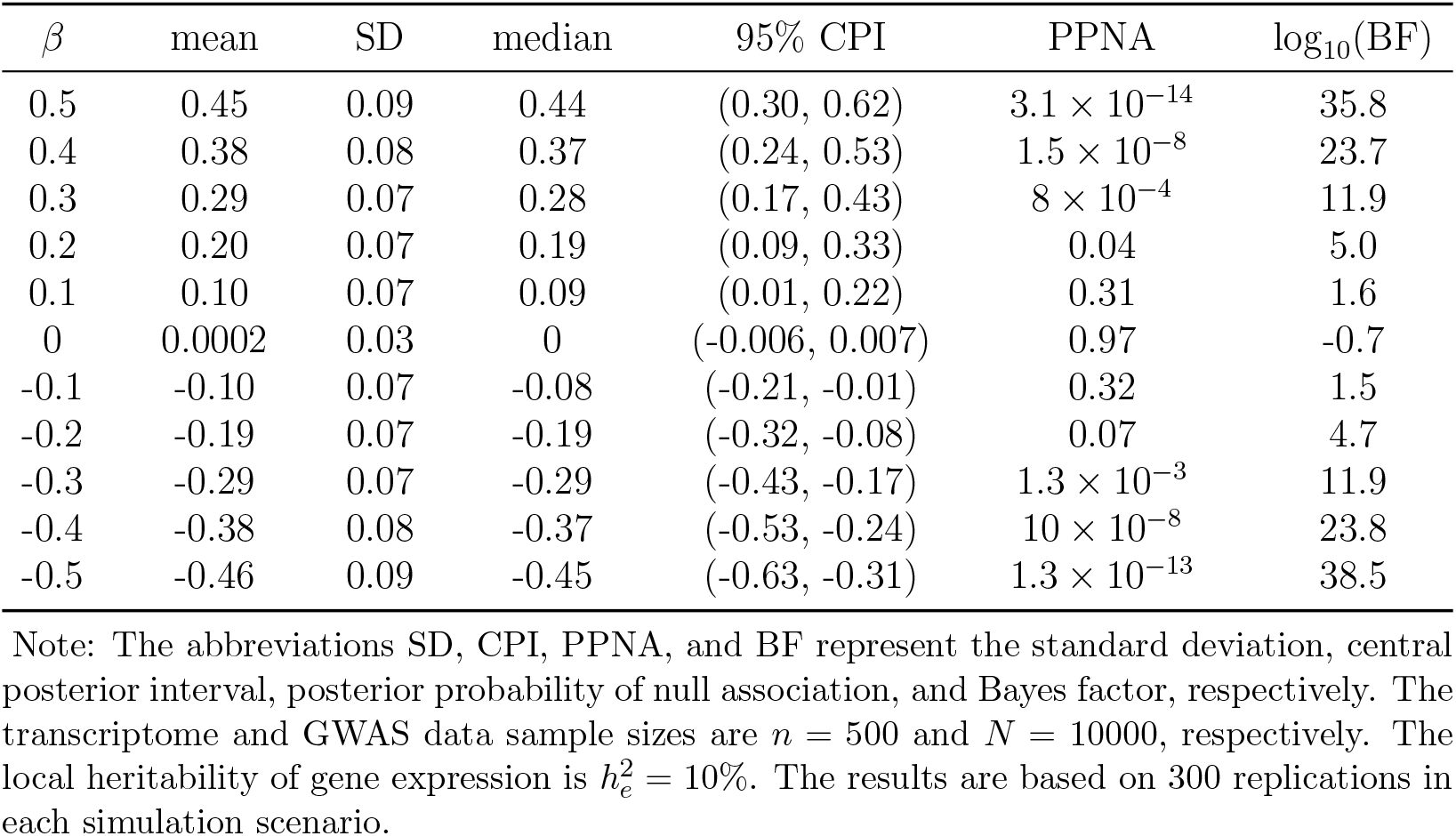
Summary of GReX effect sizes on the outcome trait estimated by glossTWAS and Bayes factors and posterior probability of null associations for single-ancestry TWAS.

We compare glossTWAS with PMR-Egger [14], TIGAR [11], and PrediXcan [3] for a single-ancestry TWAS. We first compare the methods in terms of their accuracy in correctly classifying the null and non-null genes. The partial receiver operating characteristic (ROC) curve for glossTWAS consistently dominates the partial ROC curves for PMR-Egger, TIGAR, and PrediXcan consistently for the majority of simulation settings (Figure 1). This implies that, for a given level of false positive rate (FPR), glossTWAS provides a higher true positive rate (TPR). Consequently, glossTWAS yields a higher area under the curve (AUC) than the other approaches for various choices of the simulation parameters: transcriptome data sample size (*n*), GWAS data sample size (*N* ), gene expression heritability 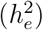, and the GWAS complex trait heritability 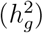. The increase in AUC for glossTWAS varies in the range of 1% − 6% compared to the other methods. For example, when *n* = 500, *N* = 10000 and 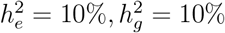, the AUC for glossTWAS is 0.76 while that of PMR-Egger, TIGAR, and PrediXcan are 0.74, 0.73, and 0.73 respectively (Figure 1A). When we increase the transcriptome data sample size from 500 to 1000 while keeping the other parameters fixed, glossTWAS yields an AUC of 0.75, compared to AUCs of 0.72, 0.69, and 0.73 for PMR-Egger, TIGAR, and PrediXcan, respectively (Figure 1E). We observe that for a given choice of 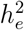 and 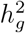, increasing the GWAS data sample size (*N* ) leads to better accuracy for all approaches than increasing the transcriptome data sample size (*n*) (simulation setting A vs. I in Figure 1). In these scenarios as well, glossTWAS achieves higher AUCs than the other approaches: 0.83 vs 0.79, 0.8, 0.81. We evaluated the methods with respect to the classification of null and non-null genes based on the partial ROC curves, primarily because the Bayesian measures of association are not directly comparable to the frequentist measures.

**Figure 1.**
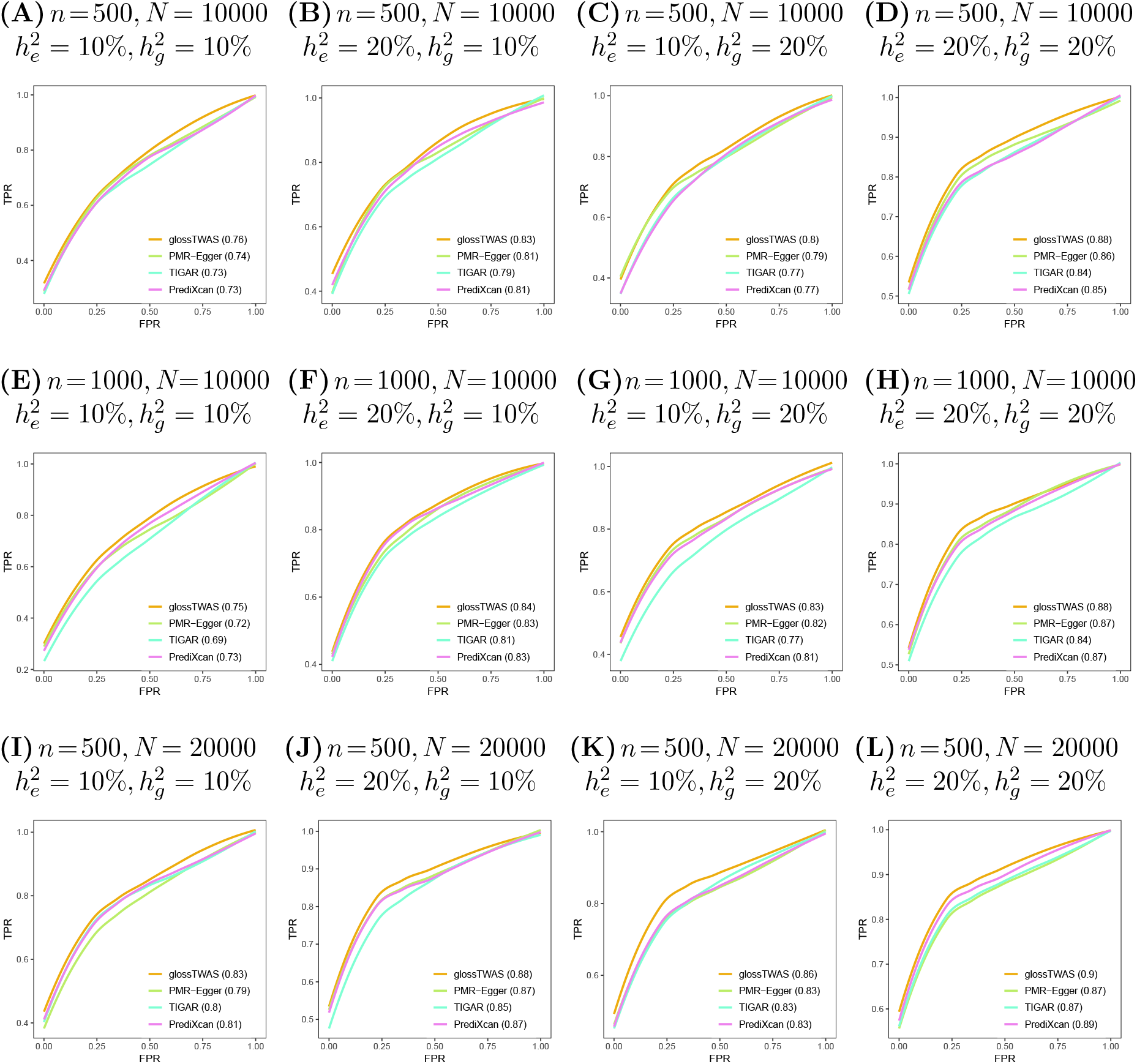
Partial receiver operating characteristic (ROC) curves for glossTWAS, PMR-Egger, TIGAR, and PrediXcan to contrast their accuracy in correctly classifying the null and non-null genes. The choices of transcriptome data sample size (*n*), GWAS data sample size (*N* ), local heritability of gene expression 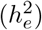, and trait heritability 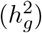 are varied. Each method’s true positive rate (TPR) and false positive rate (FPR) are calculated based on 50 replications in a given simulation scenario. The area under the curve (AUC) is provided in the first bracket corresponding to each method’s name in the plots.

Next, we assess whether the different approaches adequately control the false discovery rate (FDR) in the multiple hypothesis testing setup. Since glossTWAS is a Bayesian approach, we use the Bayesian false discovery rate (BFDR). For the frequentist approaches, we implement the Benjamini-Hochberg (BH) procedure [19] to control the FDR at a given threshold. We note that the BFDR is the Bayesian analog of FDR. Hence, they are not directly comparable. The realized FDR (rFDR) is defined as the number of false positive genes (null genes found significantly associated) divided by the total number of significant genes. We evaluate whether rFDR is controlled while implementing glossTWAS for a given threshold of BFDR.

We observe that while controlling the BFDR at a given threshold, glossTWAS controls the rFDR below or at the same threshold. We consistently observe the rFDR control for varying choices of *n, N*, 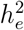, and 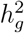 (Figure 2). At BFDR = 0.01, glossTWAS yields a mean rFDR of 0 for all simulation scenarios except for a few cases, such as *n* = 1000, *N* = 10000, 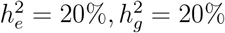, for which the estimate is 0.01 (Table S1). A possible explanation for the zero estimate is that the BFDR threshold of 0.01 is too stringent to make a false discovery when a limited number of hypotheses (200) are tested. When BFDR is controlled at 0.05 and *n* = 500, *N* = 10000, the mean rFDR for glossTWAS are 0.01, 0.02, 0.01, 0.01 for the choices of 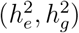 as (10%, 10%), (10%, 20%), (20%, 10%), (20%, 20%), respectively (Table S1). For BFDR = 0.1, glossTWAS yielded the mean rFDR as 0.01, 0.04, 0.05, 0.04 for the same choices of the heritability (Table S1). For this setting, the rFDRs are 0.1, 0.13, 0.12, 0.14 when BFDR = 0.2. Thus, the realized FDR for glossTWAS is overall controlled at or below the BFDR threshold in various simulation settings.

**Figure 2.**
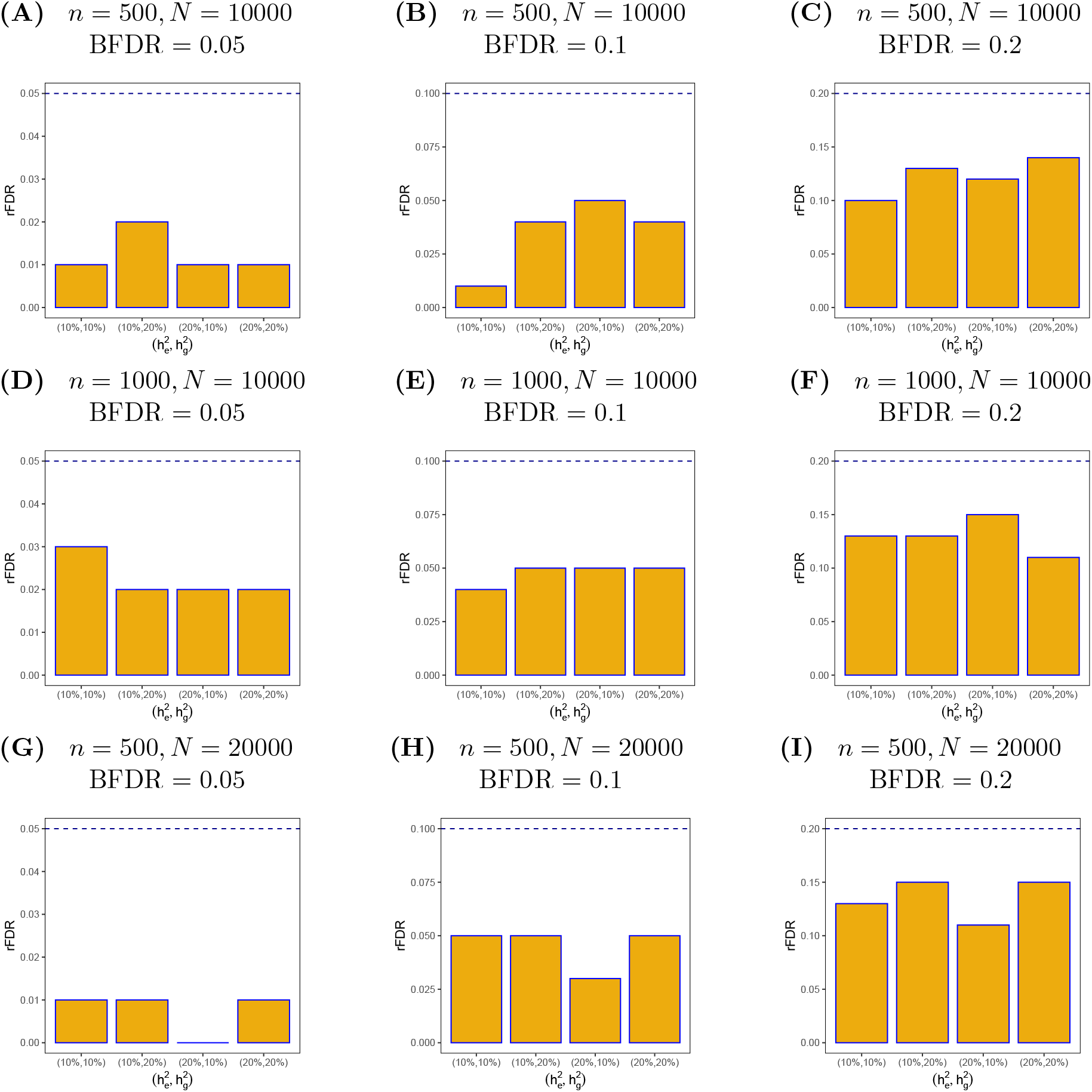
Bar plots of realized FDR (rFDR) for different levels of Bayesian FDR (BFDR) for glossTWAS in the single-ancestry TWAS setting. The choice of heritabilities are varied. In the x-axis, 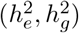 denotes the gene expression heritability and outcome trait heritability. The choices of transcriptome data sample size (*n*) and GWAS data sample size (*N* ) are varied. In each plot, the actual level of BFDR is denoted by the dashed horizontal blue line. The rFDR is calculated based on 50 replications in a given simulation scenario.

While controlling the rFDR, glossTWAS offers a good true positive rate (TPR). For BFDR = 0.05, *n* = 500, and *N* = 10000, the mean TPR varies in the range of 0.14 to 0.49 (Table S1). In general, for a given level of BFDR, the TPR increases with the increase in either expression or trait heritability or both (Figure 3). For example, when 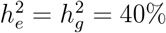, the TPR reached to 0.76 for the above setting (Table S1). The TPR will tend to one with higher choices of heritability. Otherwise, for a given choice of heritabilities, the TPR will continue to increase as the sample sizes increase.

**Figure 3.**
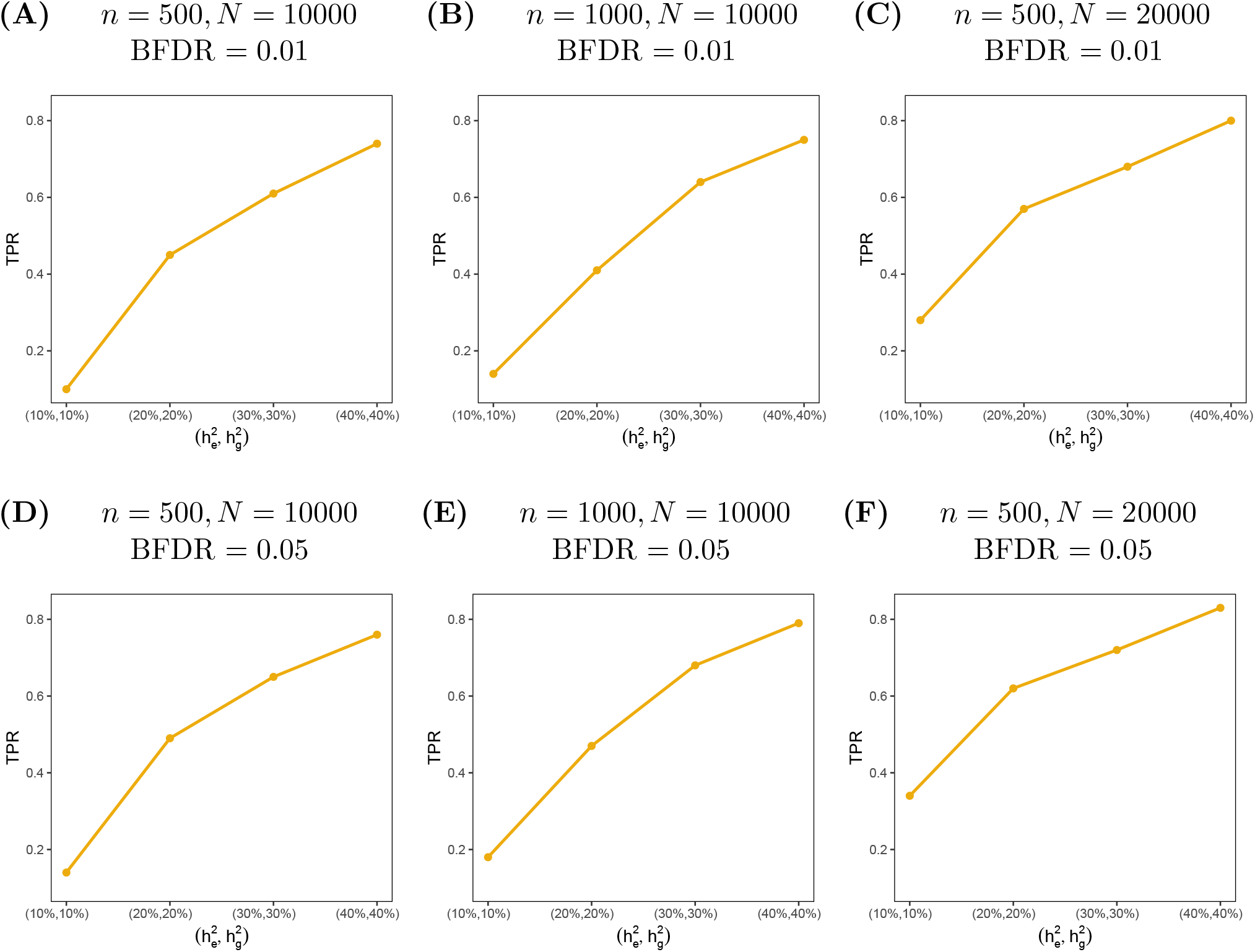
Line plots of true positive rate (TPR) by glossTWAS against different choices of heritabilities in the single-ancestry TWAS setting. In the x-axis, 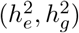 denotes the gene expression heritability and outcome trait heritability. The choices of transcriptome data sample size (*n*) and GWAS data sample size (*N* ) are varied. The TPR is calculated based on 50 replications in a given simulation scenario.

For the classical approaches, while PMR-Egger mostly controls the rFDR at different levels, PrediXcan and TIGAR frequently breach the desired threshold. For a couple of settings, the rFDR obtained by PMR-Egger exceeded the target FDR threshold. When *n* = 500, *N* = 10000, the rFDR for PMR-Egger for the FDR threshold = 0.01, 0.05 are 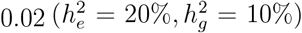 and 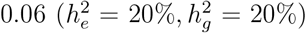, respectively (Table S2). In many instances, TIGAR and PrediXcan do not control the FDR at desired levels, regardless of varying sample sizes and heritability choices. For example, when *n* = 500, *N* = 10000, and FDR level = 0.05, both TIGAR and PrediXcan overestimate the rFDR as 0.08 for 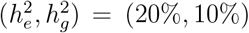 (Table S2). PrediXcan generally shows higher TPR than PMR-Egger, albeit at the cost of higher FDR. For example, when *n* = 1000 and *N* = 10000, PMR-Egger yields a TPR in the range of 0.1 − 0.44 at FDR level = 0.01, while PrediXcan produced a TPR of 0.14−0.46. However, the rFDR ranges in (0−0.01) for PMR-Egger and (0−0.03) for PrediXcan (Table S2). In these settings, glossTWAS offers TPR in (0.14−0.41) while controlling BFDR at 0.01 (Table S1). We note that the TPR estimates obtained by glossTWAS and PMR-Egger are not comparable, as they were obtained while controlling for BFDR and FDR, respectively. However, the partial ROC curves demonstrate that glossTWAS offers a higher TPR for the same false positive rate obtained by both methods.

Next, we discuss the relative performance of these methods while estimating the GReX effect sizes. For a small non-null effect, *β* = 0.1, glossTWAS accurately estimates it with a mean 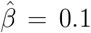 and SD = 0.07 across repetitions. The other methods overestimate it: for PMR-Egger, mean 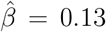 with SD = 0.08; for TIGAR, mean 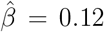 with SD = 0.04; for PrediXcan, mean 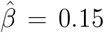 with SD = 0.05 (Table S3). As we increase the absolute values of *β* from 0.2 to 0.5, all the methods underestimate the true *β*. Compared to the other approaches, TIGAR substantially underestimates the larger effect sizes (Table S3). GlossTWAS produces similar mean effect sizes to those of PMR-Egger and PrediXcan in these cases. For example, when *β* = −0.4, glossTWAS, PMR-Egger, and PrediXcan produce similar estimates with mean 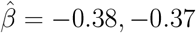, and − 0.37, respectively, whereas TIGAR significantly underestimates it with mean 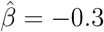 (Table S3).

We now evaluate the relative bias and root mean squared error (RMSE) of the estimates. The relative bias is defined as 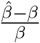, where 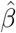 is an estimate of *β*. GlossTWAS exhibits lesser relative bias for varying choices of effect sizes compared to the competing approaches (Figure 4A). For *β* = 0.1, 0.2, 0.3, the mean relative bias for glossTWAS is −0.8%, 0.5%, 3%, respectively. PMR-Egger, TIGAR, and PrediXcan yield the mean relative bias of (33.4%, 2.2%, −4.1%), (18.8%, −15.2%, −24.7%), and (52.5%, 6.2%, −4.1%), respectively, for these choices of *β* (Table S4). Therefore, the magnitude of the mean relative bias for glossTWAS is substantially lower than that of the other approaches. The SDs of the relative bias estimates for glossTWAS are lower than those of the other methods. It gradually decreases as the absolute values of *β* increase. For *β* = 0.4, 0.5, glossTWAS produces an SD of relative biases as 17%, 14%. PMR-Egger, TIGAR, and PrediXcan produce (18%, 15%), (25%, 24%), and (34%, 30%) SDs respectively (Table S4). Thus, glossTWAS performs better than the others in terms of the relative bias of the effect size estimates.

**Figure 4.**
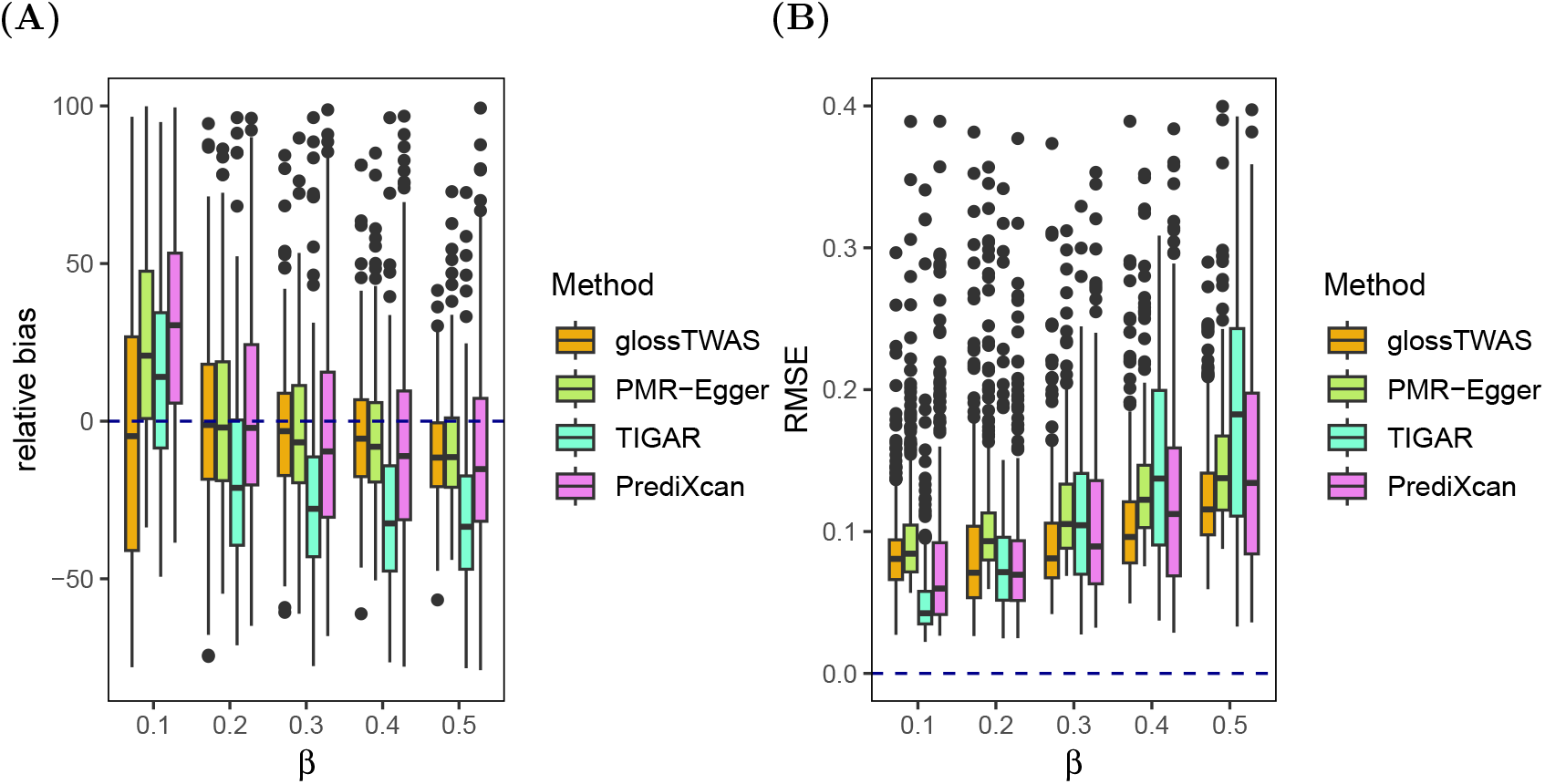
Comparison of accuracy in TWAS effect size estimation by glossTWAS, PMR-EGGER, TIGAR, and PrediXcan for single-ancestry TWAS. Two measures are considered: (**A**) relative bias and (**B**) root mean squared error (RMSE). The transcriptome and GWAS data sample sizes are *n* = 500 and *N* = 10000, respectively. The local heritability of gene expression is fixed at 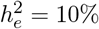.

In terms of the RMSE, glossTWAS consistently performs better than the other approaches, yielding either comparable or smaller estimates of the RMSE across the different choices of *β* (Table S3). The RMSEs increase with the true *β* for all approaches. GlossTWAS yields lower RMSEs than others for larger effects (Figure 4B). For the null effect of GReX, i.e., when *β* = 0, glossTWAS produces substantially lower bias 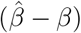 and RMSE than the other approaches (Figure S2). Overall, glossTWAS achieves better precision in estimating the GReX effect sizes. We note that glossTWAS offers an overall better accuracy in the effect size estimation than PMR-Egger, which is a unified frequentist approach that models the transcriptome and GWAS data jointly.

### 2.3 Multi-ancestry TWAS

We first describe a summary of outputs obtained by glossTWAS when the proportion of eQTLs overlapping between the two ancestries (*f*_0_) is 50%, and the TWAS effect sizes in the two ancestries are equal, i.e., *β*_1_ = *β*_2_. Results are similar when the effect sizes are not equal. GlossTWAS provides comparable estimates of effect sizes based on the posterior mean and median (Table 2). When *β*_1_ = *β*_2_ = 0.2, the posterior means for both ancestries are 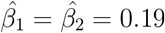 with the same posterior SDs of 0.06. In this case, the posterior medians are 0.18 and 0.19 for two ancestries (Table 2). For negative effects *β*_1_ = *β*_2_ = −0.2, glossTWAS yields posterior means as 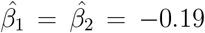, with posterior SDs of 0.06 for each. In this case, the posterior medians are both equal to −0.18. The 95% CPIs for *β*_1_ and *β*_2_ are (0.01, 0.18) and (0.01, 0.19), respectively. The CPI has a larger width for larger effect sizes relative to smaller effects. When *β*_1_ = *β*_2_ = 0.4, the 95% CPIs for *β*_1_ and *β*_2_ are (0.25, 0.5) and (0.25, 0.51). GlossTWAS accurately estimates the null effect of GReX, *β*_1_ = *β*_2_ = 0 (Table 2). While testing for an association when there is no effect, glossTWAS efficiently produces a mean value of log_10_(BF) = −0.6 and PPNA = 0.96 (Table 2) across simulation iterations. For the non-null choices of *β*_1_ and *β*_2_, mean log_10_(BF) increases and PPNA decreases as the absolute values of *β*_1_ and *β*_2_ increase (Table 2).

**Table 2.** Summary of GReX effect sizes estimated by multi-ancestry glossTWAS. Summary of Bayes factor and PPNA are also provided. Here, half of the eQTLs are shared between the two ancestries, i.e., *f*_0_ = 50%.

| $\beta_1$ | $\beta_2$ | EAS ancestry | | | | EUR ancestry | | | | PPNA | $\log_{10}(\text{BF})$ |
| --- | --- | --- | --- | --- | --- | --- | --- | --- | --- | --- | --- |
|  |  | mean | SD | median | 95% CPI | mean | SD | median | 95% CPI |  |  |
| 0.5 | 0.5 | 0.45 | 0.08 | 0.44 | (0.31, 0.60) | 0.45 | 0.08 | 0.44 | (0.31, 0.60) | $1.6 \times 10^{-40}$ | 75.8 |
| 0.4 | 0.4 | 0.37 | 0.07 | 0.36 | (0.25, 0.50) | 0.37 | 0.07 | 0.36 | (0.25, 0.51) | $1.6 \times 10^{-21}$ | 48.9 |
| 0.3 | 0.3 | 0.29 | 0.06 | 0.28 | (0.18, 0.42) | 0.29 | 0.06 | 0.28 | (0.18, 0.41) | $2.3 \times 10^{-10}$ | 27.4 |
| 0.2 | 0.2 | 0.19 | 0.06 | 0.18 | (0.09, 0.30) | 0.19 | 0.06 | 0.19 | (0.09, 0.30) | $6.8 \times 10^{-3}$ | 9.8 |
| 0.1 | 0.1 | 0.08 | 0.06 | 0.07 | (0.01, 0.18) | 0.09 | 0.06 | 0.08 | (0.01, 0.19) | 0.23 | 2.1 |
| 0 | 0 | 0.0002 | 0.02 | 0 | (-0.004, 0.005) | -0.0002 | 0.02 | 0 | (-0.004, 0.004) | 0.96 | -0.6 |
| -0.1 | -0.1 | -0.09 | 0.06 | -0.08 | (-0.19, -0.01) | -0.09 | 0.06 | -0.08 | (-0.19, -0.01) | 0.21 | 2.1 |
| -0.2 | -0.2 | -0.19 | 0.06 | -0.18 | (-0.30, -0.09) | -0.19 | 0.06 | -0.18 | (-0.30, -0.08) | 0.01 | 9.3 |
| -0.3 | -0.3 | -0.29 | 0.06 | -0.28 | (-0.41, -0.18) | -0.29 | 0.07 | -0.28 | (-0.42, -0.18) | $3.3 \times 10^{-10}$ | 26.5 |
| -0.4 | -0.4 | -0.38 | 0.07 | -0.37 | (-0.51, -0.25) | -0.37 | 0.07 | -0.37 | (-0.51, -0.25) | $3.4 \times 10^{-22}$ | 51.8 |
| -0.5 | -0.5 | -0.45 | 0.08 | -0.44 | (-0.59, -0.31) | -0.45 | 0.08 | -0.44 | (-0.60, -0.31) | $4.9 \times 10^{-36}$ | 76 |

Note: $\beta_1$ and $\beta_2$ denote the true GReX effect sizes for the EAS and EUR ancestries, respectively. The abbreviations SD, CPI, PPNA, and BF represent the standard deviation, central posterior interval, posterior probability of null association, and Bayes factor, respectively. The transcriptome data sample sizes are $n_1 = n_2 = 500$ , and GWAS data sample sizes are $N_1 = N_2 = 10000$ for the two ancestries. The local heritability of gene expression is fixed at $h_e^2 = 10\%$ in each ancestry. The summaries are obtained based on 300 effective replications in a given simulation scenario.

Importantly, glossTWAS performs efficiently when the GReX effect sizes are unequal between the two ancestries. For *β*_1_ = 0.1, *β*_2_ = 0.5, the posterior means of effect sizes are 0.1 and 0.46 corresponding to the EAS and EUR ancestries, respectively. The posterior SDs are 0.05 and 0.09, respectively. The 95% CPI for *β*_1_ = 0.1 is (0.02, 0.19), and (0.31, 0.63) for *β*_2_ = 0.5 (Table 3), which contain the true effect sizes. Similarly, for negative effect sizes, when *β*_1_ = −0.1 and *β*_2_ = −0.5, the posterior means are −0.1 and −0.46 for the two ancestries with posterior SDs 0.05 and 0.09. In this setting, log_10_(BF) and PPNA are 36.6 and 3.8 *×* 10^−15^, respectively, implying a strong association (Table 3). When eQTLs do not overlap between the ancestries, i.e., *f*_0_ = 0, glossTWAS provides a similar posterior summary both for equal (Table S5) and unequal (Table S6) choices of *β*_1_ and *β*_2_.

**Table 3.**
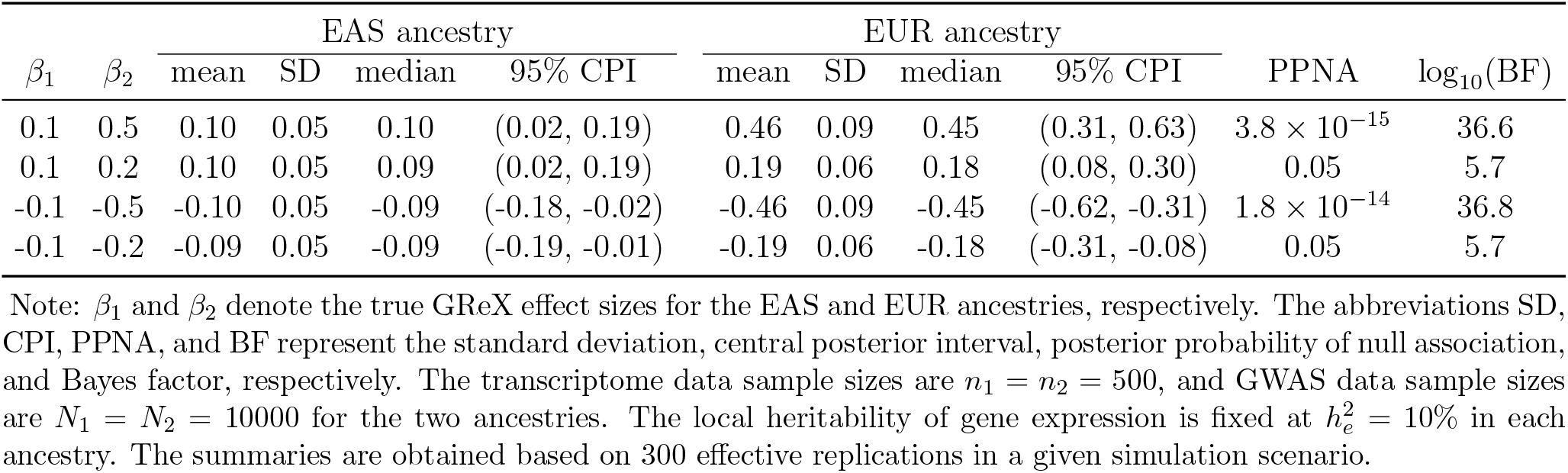
Summary of GReX effect sizes estimated by multi-ancestry glossTWAS when the effect sizes differ across the two ancestries. Summary of Bayes factor and PPNA are also provided. Here, half of the eQTLs are shared between the two ancestries, i.e., *f*_0_ = 50%.

| $\beta_1$ | $\beta_2$ | EAS ancestry | | | | EUR ancestry | | | | PPNA | $\log_{10}(\text{BF})$ |
| --- | --- | --- | --- | --- | --- | --- | --- | --- | --- | --- | --- |
|  |  | mean | SD | median | 95% CPI | mean | SD | median | 95% CPI |  |  |
| 0.1 | 0.5 | 0.10 | 0.05 | 0.10 | (0.02, 0.19) | 0.46 | 0.09 | 0.45 | (0.31, 0.63) | $3.8 \times 10^{-15}$ | 36.6 |
| 0.1 | 0.2 | 0.10 | 0.05 | 0.09 | (0.02, 0.19) | 0.19 | 0.06 | 0.18 | (0.08, 0.30) | 0.05 | 5.7 |
| -0.1 | -0.5 | -0.10 | 0.05 | -0.09 | (-0.18, -0.02) | -0.46 | 0.09 | -0.45 | (-0.62, -0.31) | $1.8 \times 10^{-14}$ | 36.8 |
| -0.1 | -0.2 | -0.09 | 0.05 | -0.09 | (-0.19, -0.01) | -0.19 | 0.06 | -0.18 | (-0.31, -0.08) | 0.05 | 5.7 |

Note: $\beta_1$ and $\beta_2$ denote the true GReX effect sizes for the EAS and EUR ancestries, respectively. The abbreviations SD, CPI, PPNA, and BF represent the standard deviation, central posterior interval, posterior probability of null association, and Bayes factor, respectively. The transcriptome data sample sizes are $n_1 = n_2 = 500$ , and GWAS data sample sizes are $N_1 = N_2 = 10000$ for the two ancestries. The local heritability of gene expression is fixed at $h_e^2 = 10\%$ in each ancestry. The summaries are obtained based on 300 effective replications in a given simulation scenario.

We compare the performance of glossTWAS with METRO in terms of the correct classification of null and non-null genes. In various simulation scenarios, where the proportion of eQTLs shared between ancestries (*f*_0_), expression heritability 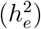, and trait heritability 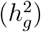 vary, the partial ROC curves for glossTWAS mostly dominate those for METRO (Figure 5). In most scenarios, glossTWAS yields an increase in AUC of 1% − 5%. For example, when *f*_0_ = 0, 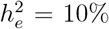, and 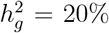, the mean AUC for glossTWAS and METRO are 0.87 and 0.82, respectively (Figure 5B). For the same choice of heritabilities with *f*_0_ = 50%, glossTWAS and METRO yielded mean AUCs of 0.87 and 0.84, respectively (Figure 5F).

**Figure 5.**
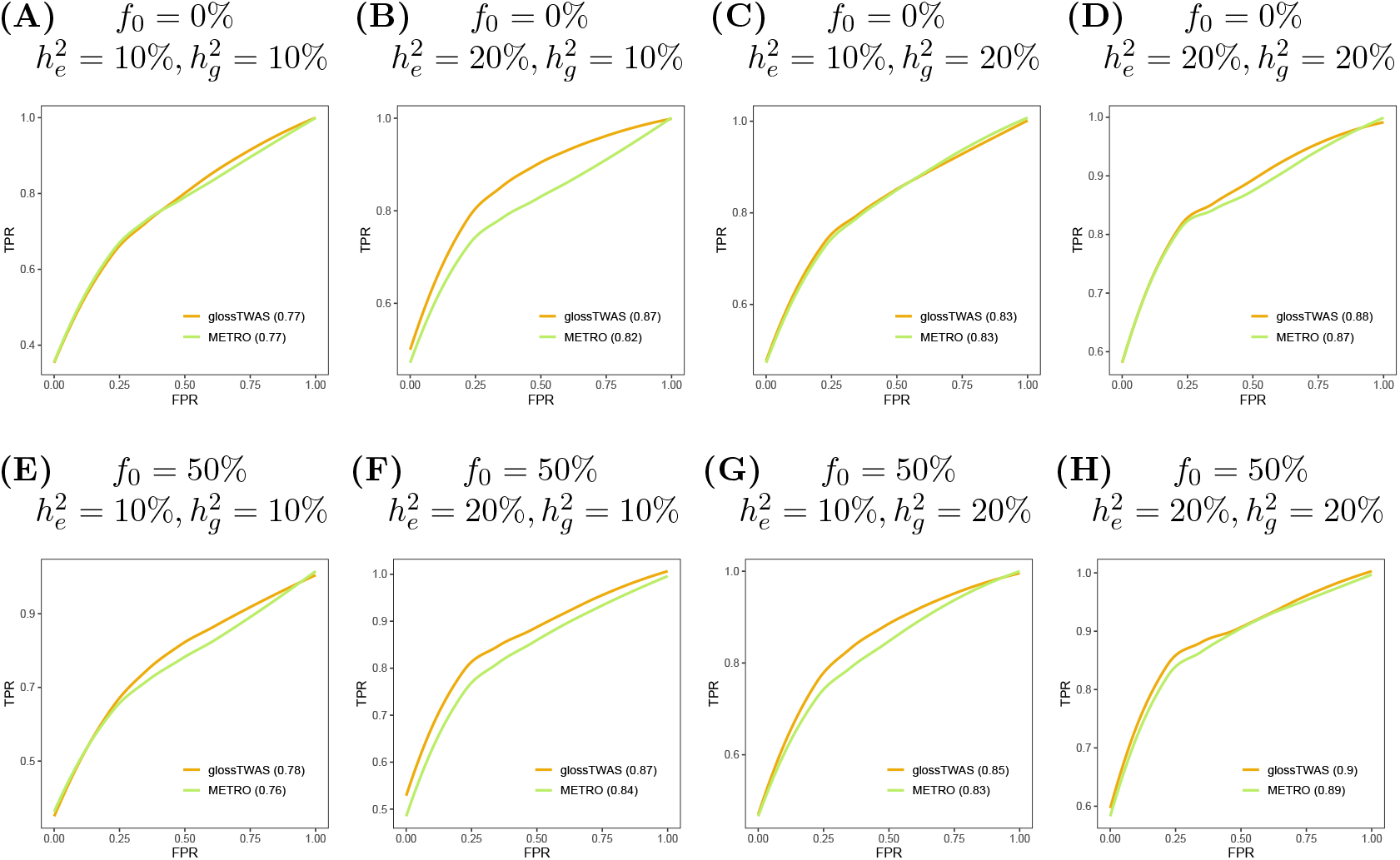
Partial receiver operating characteristic (ROC) curves for multi-ancestry glossTWAS and METRO to contrast their accuracy in correctly classifying null and non-null genes. The choices of the proportion of eQTLs shared between the two ancestries (*f*_0_), local heritability of gene expression 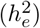, and trait heritability 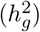 are varied. The transcriptome data sample sizes are *n*_1_ = *n*_2_ = 500, and GWAS data sample sizes are *N*_1_ = *N*_2_ = 10000 for the two ancestries. Each method’s true positive rate (TPR) and false positive rate (FPR) are calculated based on 50 replications in a given simulation scenario. The area under the curve (AUC) is provided corresponding to each method’s name in the plots.

In two out of eight settings, the methods produced the same mean AUC (Figure 5A, 5C) where the ROC curves overlap. In general, the AUC increases for both as we increase the expression heritability and/or trait heritability while keeping the other parameters fixed. When *f*_0_ = 50%, 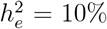, and 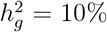, the mean AUCs are 0.78 and 0.76 for glossTWAS and METRO (Figure 5E). As we increase 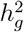 from 10% to 20%, keeping *f*_0_ and 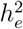 fixed, the AUCs increase to 0.85 and 0.83 for glossTWAS and METRO, respectively (Figure 5G). Overall, glossTWAS classifies the null and non-null genes more accurately than METRO.

We next assess how well multi-ancestry glossTWAS controls the error rate with respect to the Bayesian false discovery rate (BFDR). For various choices of *f*_0_, 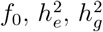, glossTWAS controls the realized false discovery rate (rFDR) below or at different levels of BFDR (Figure 6). For the scenario with *f*_0_ = 0 and BFDR= 0.01, glossTWAS yields rFDR of 0, 0.01, 0, and 0.01 when 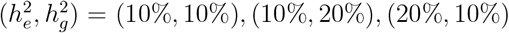, and (20%, 20%), respectively (Table S7). For the same settings with BFDR = 0.05, glossTWAS yields rFDR of 0.01, 0.03, 0.02, and, 0.01. When *f*_0_ = 50%, the rFDRs are 0.01, 0.02, 0.02, 0.01 (Table S7). For BFDR = 0.1, the rFDR lies in the ranges of 0.03−0.08 and 0.03−0.06 for *f*_0_ = 0% and *f*_0_ = 50%, respectively, under the same choices of heritabilities (Figure 6B,6E). Thus, multi-ancestry glossTWAS controls the rFDR at different levels of BFDR. While testing for an association in either ancestry, glossTWAS produces a good true positive rate (TPR) for different thresholds of BFDR (Figure 7). Given *f*_0_ = 0 and BFDR = 0.05, the TPR lies in the range of 0.24 − 0.58 for the varying choices of heritabilities (Table S7). In general, TPR increases with increasing heritability for fixed BFDR. For example, when BFDR = 0.01, the mean TPR for glossTWAS is 0.21 when 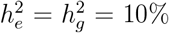 and 0.8 when 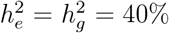 (Table S7). The TPR will reach one for higher values of these heritabilities. For fixed heritabilities, the TPR will continue to increase as the sample sizes increase.

**Figure 6.**
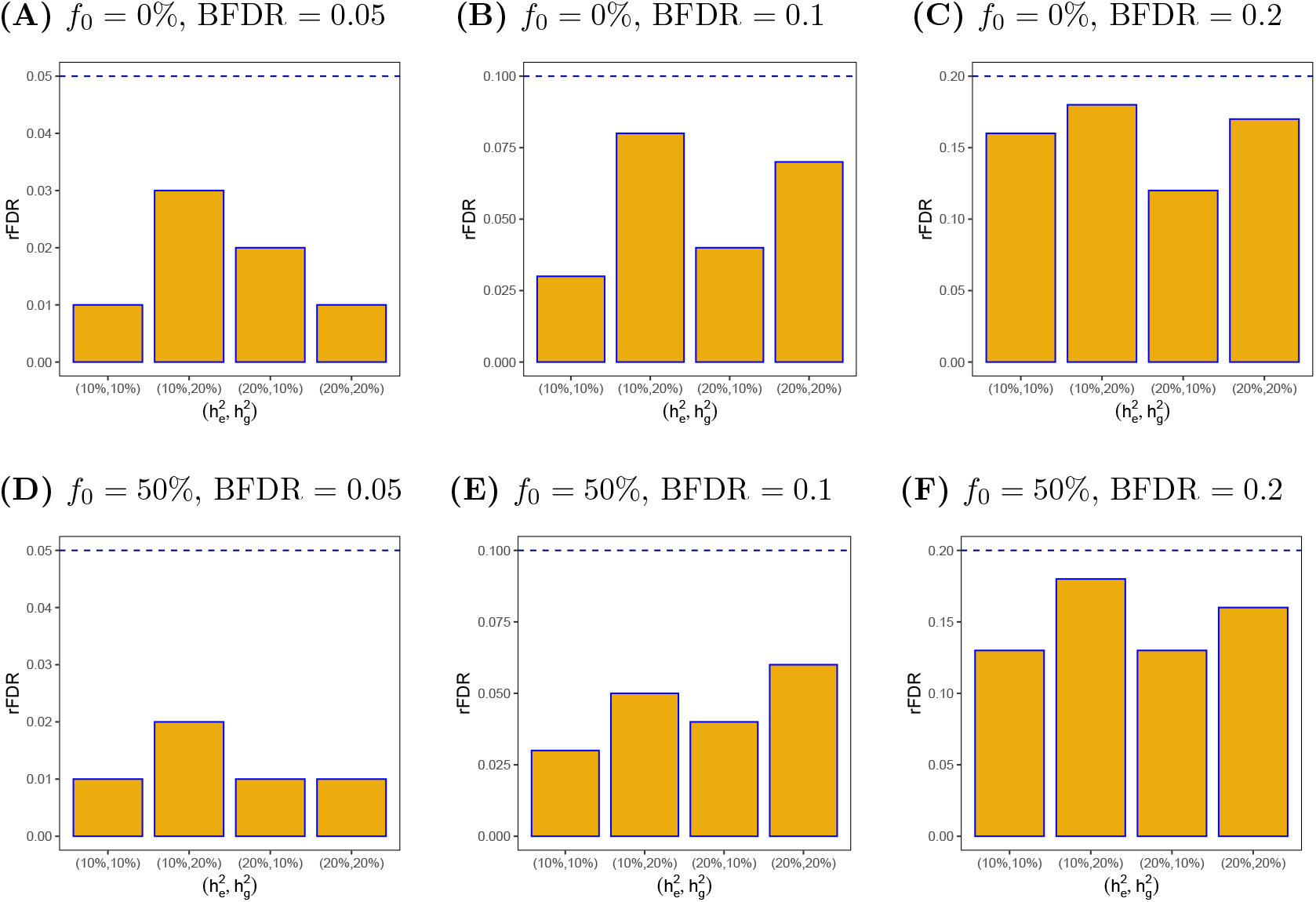
Bar plots of realized FDR (rFDR) for different levels of Bayesian FDR (BFDR) for multi-ancestry glossTWAS. The choice of heritabilities are varied. In the x-axis, 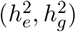 denotes the gene expression heritability and outcome trait heritability. The choices of the proportion of eQTLs shared between the two ancestries (*f*_0_) are varied. The transcriptome data sample sizes are *n*_1_ = *n*_2_ = 500, and GWAS data sample sizes are *N*_1_ = *N*_2_ = 10000 for the two ancestries. In each plot, the true level of BFDR is denoted by the dashed horizontal blue line. The rFDR is calculated based on 50 replications in a given simulation scenario.

**Figure 7.**
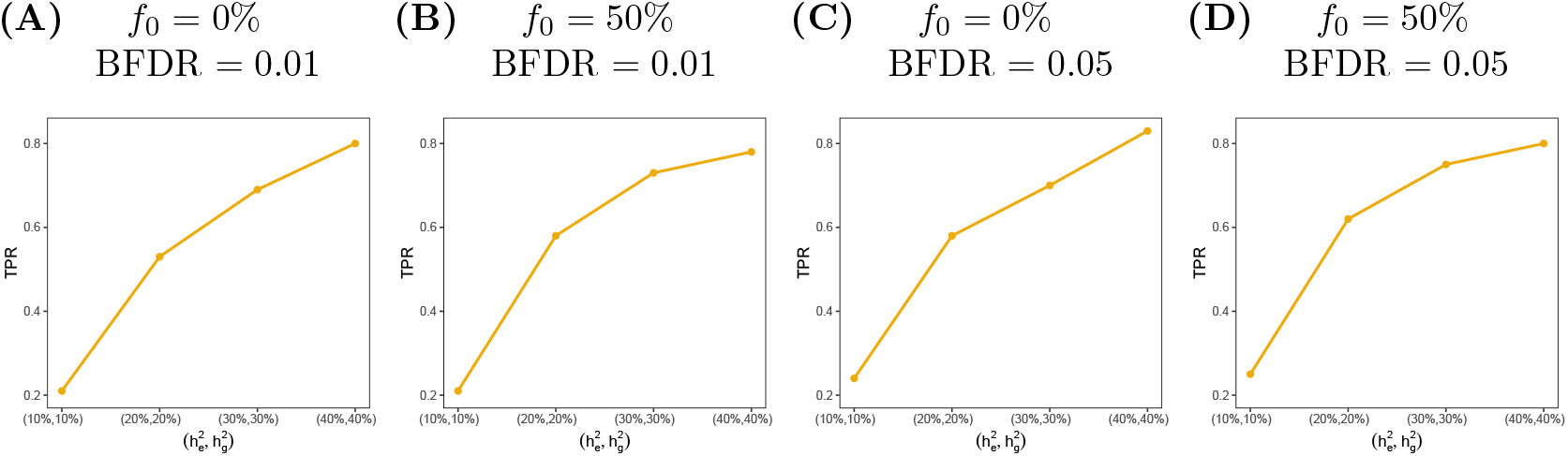
Line plots of true positive rate (TPR) for multi-ancestry glossTWAS. The choice of heritabilities are varied. In the x-axis, 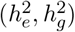 denotes the gene expression heritability and outcome trait heritability. The choices of the proportion of eQTLs shared between the two ancestries (*f*_0_) are varied. The transcriptome data sample sizes are *n*_1_ = *n*_2_ = 500, and GWAS data sample sizes are *N*_1_ = *N*_2_ = 10000 for the two ancestries. The TPR is calculated based on 50 replications in a given simulation scenario.

The frequentist approach METRO adequately controls the FDR at various thresholds except for one setting. For *f*_0_ = 0, 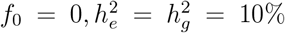, the rFDR is 0.02 for the target FDR = 0.01 (Table S8). METRO offers good TPR in various settings (Table S8). For example, when *f*_0_ = 0 and BFDR = 0.05, the TPR lies in the range of 0.26 − 0.57 for the varying choices of heritabilities. In these scenarios, glossTWAS offered a TPR of 0.24 − 0.58. However, the TPR estimates obtained by glossTWAS and METRO are not directly comparable, as they were calculated while controlling BFDR and FDR, respectively. The partial ROC curves in the above demonstrate that glossTWAS offers higher TPR for the same false positive rate obtained by both approaches.

Next, we contrast the performance of glossTWAS with METRO in terms of the accuracy of effect size estimation. For *f*_0_ = 50% and *β*_1_ = *β*_2_, glossTWAS marginally underestimates the true effect size (Table S9). On the contrary, METRO consistently overestimates the effect sizes. In general, the estimates obtained by glossTWAS are more accurate than those by METRO. For example, when *β*_1_ = *β*_2_ = 0.1, the mean estimates by glossTWAS in the two ancestries are: 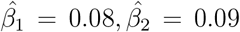 (Table S9). METRO yielded the mean 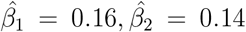. Similarly, when *β*_1_ = *β*_2_ = 0.3, glossTWAS provides the mean 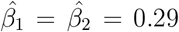 (Table S9), whereas METRO estimates 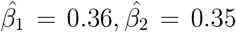. The SDs of the estimated effect sizes are also smaller for glossTWAS compared to METRO. When *β*_1_ = *β*_2_ = 0.1, the SDs for glossTWAS corresponding to EAS and EUR ancestries are both 0.05, while for METRO, the values are 0.1 and 0.11, respectively (Table S9). Concerning relative bias, glossTWAS maintains it in a lower range of magnitude than METRO (Figures 8A and 8C). For various choices of *β*_1_ and *β*_2_, glossTWAS yields the mean relative bias in a range of (−18.8%, −2.9%) for EAS ancestry, and (−14.8%, −3.6%) for EUR ancestry. For METRO, it varies in (10.7%, 55.6%) for EAS ancestry, and (9.2%, 41.7%) for EUR ancestry (Table S10). GlossTWAS produces a lower SD of relative biases than METRO (Table S10). The SD of the relative biases decreases for both methods as the magnitude of effect sizes increases. Regarding the RMSE of the estimates, glossTWAS offers a lower RMSE than METRO (Figure 8B-8D). In general, the RMSE increases with the magnitude of effect sizes for both methods. As we increase common effect sizes from 0.1 to 0.5, glossTWAS shows RMSE of 0.07 − 0.11 for either of the ancestries. In contrast, METRO exhibits higher RMSE: (0.14 − 0.19) for EAS, and (0.14 − 0.2) for EUR (Table S9).

**Figure 8.**
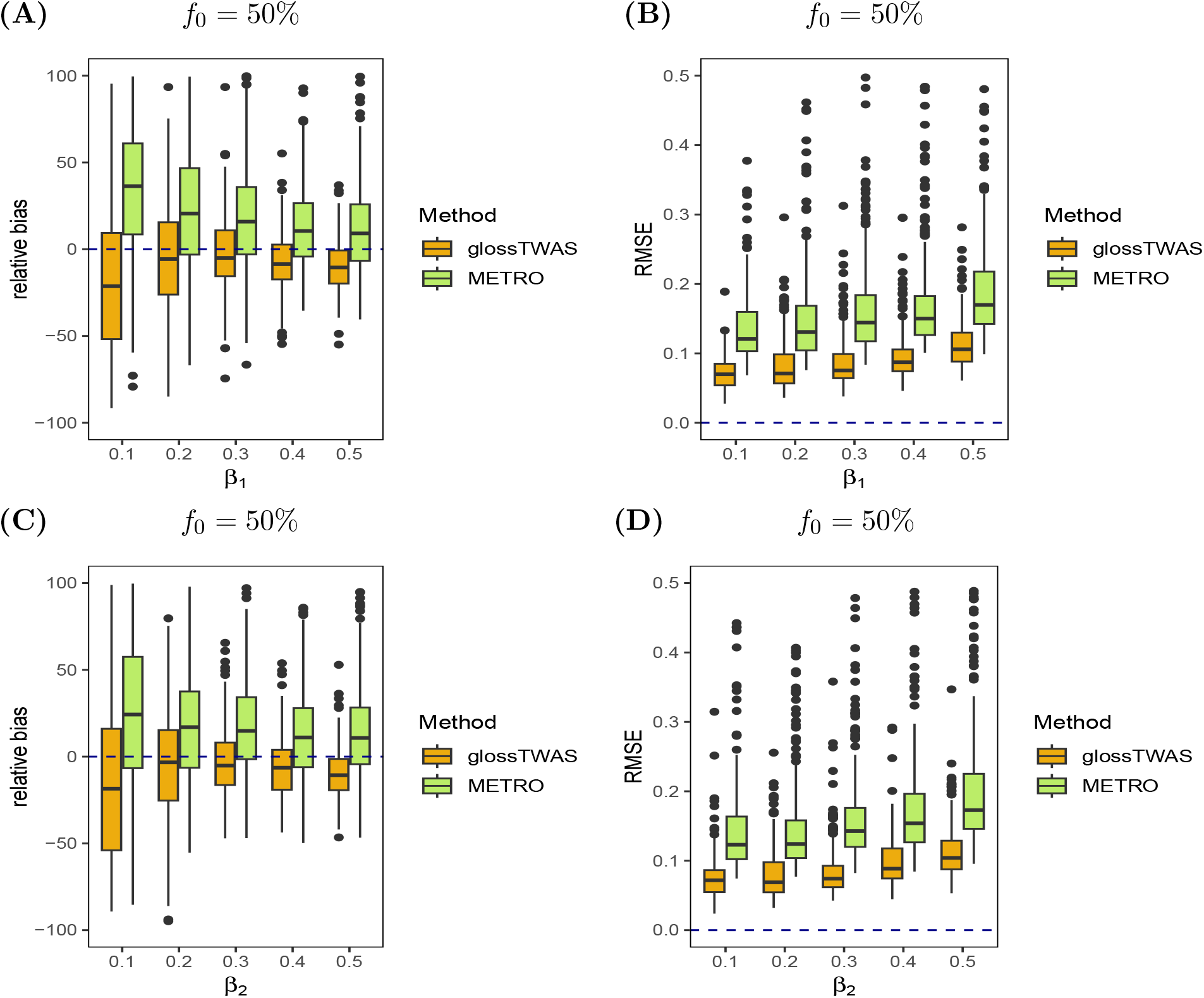
Comparison of GReX effect size estimation accuracy by multi-ancestry glossTWAS and METRO when the effect sizes corresponding to EAS ancestry (*β*_1_) and EUR ancestry (*β*_2_) are equal. Here, half of the eQTLs are shared between the two ancestries, i.e., *f*_0_ = 50%. Two measures are considered: relative bias and root mean squared error (RMSE) for the EAS ancestry (Figure **A**-**B**) and the EUR ancestry (Figure **C**-**D**). The transcriptome data sample sizes are *n*_1_ = *n*_2_ = 500, and GWAS data sample sizes are *N*_1_ = *N*_2_ = 10000 for the two ancestries. The local heritability of gene expression is fixed at 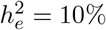 in each ancestry.

When the effect sizes are unequal, glossTWAS maintains better estimation accuracy than METRO. For *f*_0_ = 50% and *β*_1_ = 0.1, *β*_2_ = 0.2, glossTWAS yields the mean 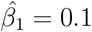 and 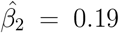, whereas METRO produces the mean 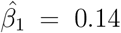 and 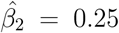. For this setting, the SDs corresponding to EAS and EUR ancestries are 0.05 and 0.06 for glossTWAS, while for METRO, they are both 0.11 (Table S11). Overall, glossTWAS exhibits lower relative bias than METRO (Figure 9A, 9C). With *f*_0_ = 50%, glossTWAS leads to −4.4% mean relative bias for estimating *β*_1_ = 0.1 and −6.7% for *β*_2_ = 0.2 (Table S12). The corresponding relative biases for METRO are 36.1% and 23.6%. The SDs of the relative bias are also lower for glossTWAS than METRO (Table S12). The RMSEs of the estimates obtained by glossTWAS are lower than those by METRO (Figure 9B,9D). For *f*_0_ = 50%, glossTWAS produces RMSEs of 0.06 and 0.12 when estimating *β*_1_ = 0.1 and *β*_2_ = 0.5 in the two ancestries, respectively. The corresponding RMSEs for METRO are 0.13 and 0.19 (Table S11). GlossTWAS performs similarly for the negative effect sizes (Figure S3, S4). Finally, glossTWAS accurately estimates the null effects, showing a tiny bias and RMSE compared to METRO (Figure S6, S5). When there is no overlap in ancestry-specific eQTLs between ancestries, i.e., *f*_0_ = 0, we observe similar comparative performance by the two methods (Table S13,S14,S15,S16, and Figure S7,S8,S9,S10).

**Figure 9.**
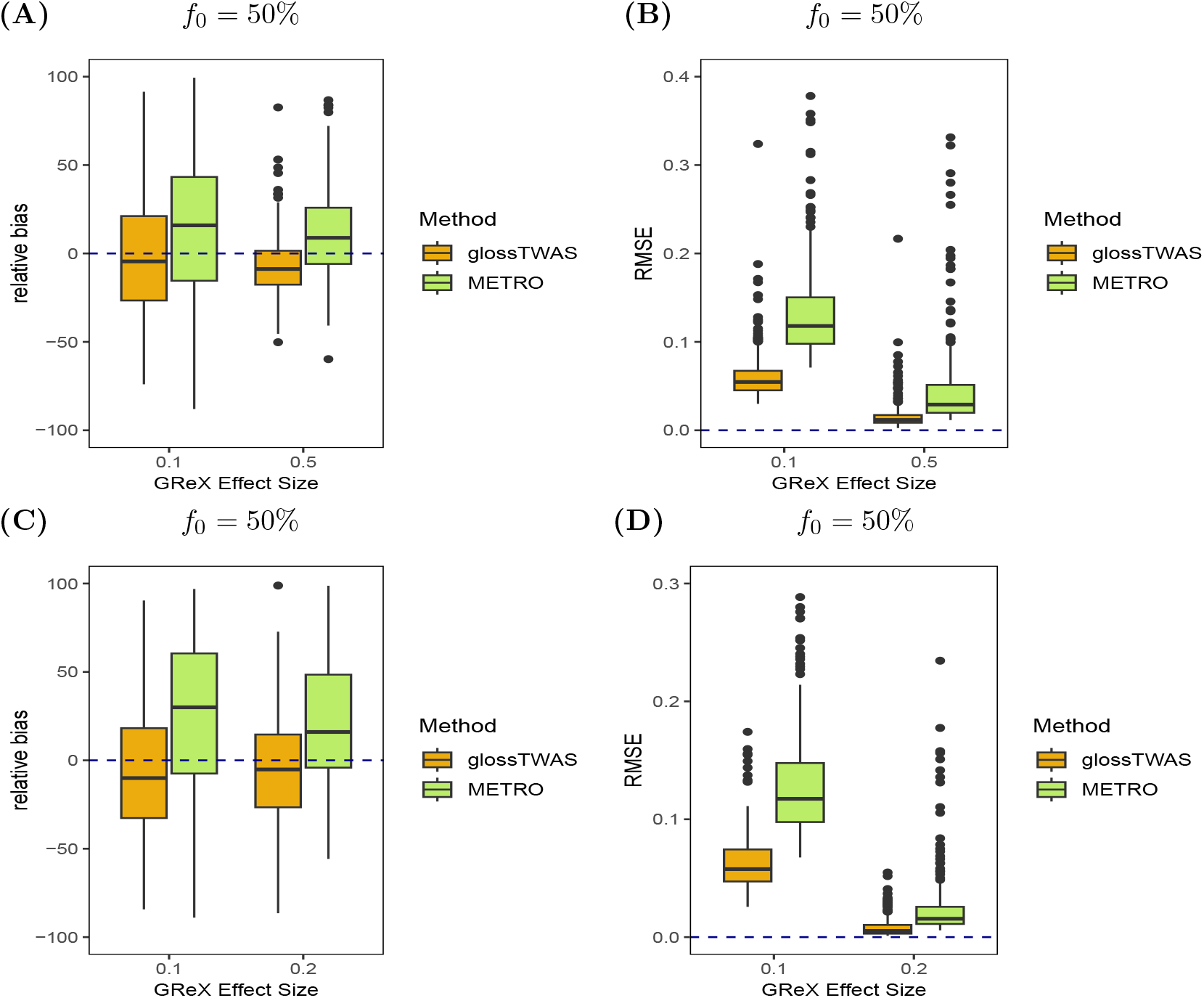
Comparison of effect size estimation accuracy by multi-ancestry glossTWAS and METRO when the GReX effect sizes corresponding to EAS ancestry (*β*_1_) and EUR ancestry (*β*_2_) are unequal. Here, half of the eQTLs are shared between the two ancestries, i.e., *f*_0_ = 50%. Two measures, relative bias and root mean squared error (RMSE), are considered for the following settings: *β*_1_ = −0.1, *β*_2_ = −0.5 (Figure **A**-**B**) and *β*_1_ = −0.1, *β*_2_ = −0.2 (Figure **C**-**D**). The transcriptome data sample sizes are *n*_1_ = *n*_2_ = 500, and GWAS data sample sizes are *N*_1_ = *N*_2_ = 10000 for the two ancestries. The local heritability of gene expression is fixed at 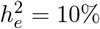 in each ancestry.

**Figure 10.**
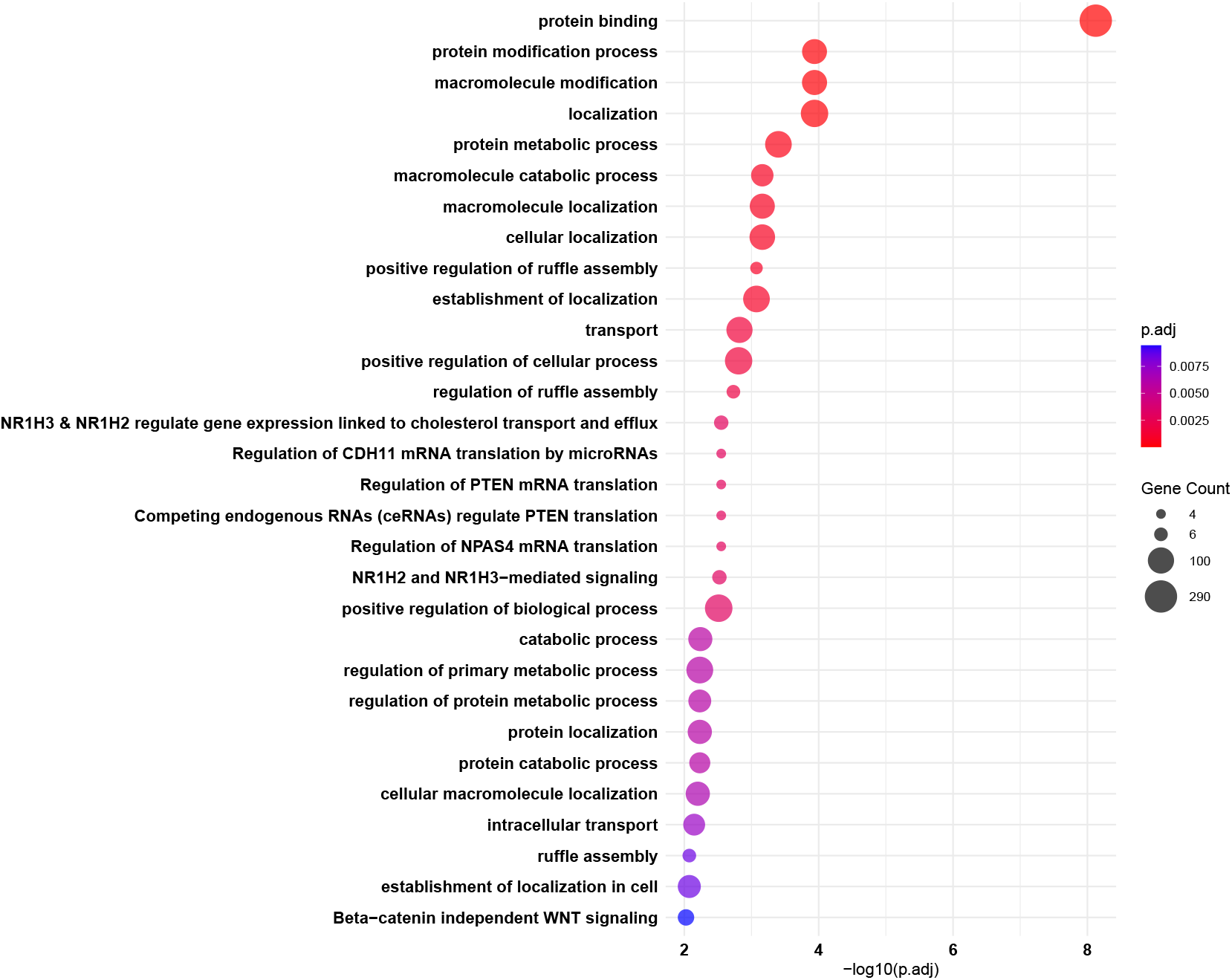
The top 30 pathways significantly over-represented (FDR *<* 0.05) in g:profiler analyses are shown for genes associated with intraocular pressure (IOP) obtained from multi-ancestry glossTWAS analyses of European and African ancestries. Enriched pathways represent shared biological processes and molecular mechanisms that may contribute to IOP regulation.

### 2.4 Real data application: TWAS for intraocular pressure

We apply our approach, glossTWAS, to conduct a single and multi-ancestry TWAS for intraocular pressure (IOP), also known as eye pressure. IOP is a measurement of the fluid pressure inside the eye. It is measured in units of millimeters of mercury (mmHg). An IOP measurement between 10 and 20 mmHg is considered normal. Too low or too high IOP can damage an individual’s vision. Elevated IOP without other symptoms is called ocular hypertension. When an individual has glaucoma, eye pressure damages the optic nerve, resulting in permanent vision loss. Without treatment, glaucoma can lead to complete blindness. Thus, it is crucial to investigate the genetic basis of IOP. Our TWAS analyses aim to identify genes that influence variation in intraocular pressure, the primary biomarker for glaucoma.

We obtained the GWAS data for IOP from the UK Biobank (UKB). We used the transcriptome data from the GEUVADIS expression study. We analyzed 12517 protein-coding genes. To adjust for multiple testing across the genes, we fix the threshold of BFDR at 1% for glossTWAS. For the frequentist approaches, we control the FDR at the same level. We acknowledge that BFDR and FDR thresholds are not directly comparable. We chose a lower 1% threshold to reduce the number of false positives in the identified set of significantly associated genes. The BFDR depends on the prior probability that a null hypothesis is true, i.e., *π*_0_ = *P* (*H*_0*i*_ is true), *i* = 1, · · ·, 12517. Similar to the simulation studies, we primarily consider *π*_0_ = 0.9. We also explored the performance for *π*_0_ = 0.95.

In single-ancestry TWAS for Europeans, glossTWAS identified 382 genes to be associated with the IOP (Table 4 and supplementary information). PMR-Egger detected 189 significant genes (Table S18). The two methods spotted a common set of 171 genes. Thus, glossTWAS detected an additional set of 211 genes associated with IOP. PrediXcan identified only 16 genes (Table S19). The small number of associations for PrediXcan is partly due to its inability to construct GReX prediction models for 5547 genes. Similar observations have also been made in previous studies [10, 11].

**Table 4.**
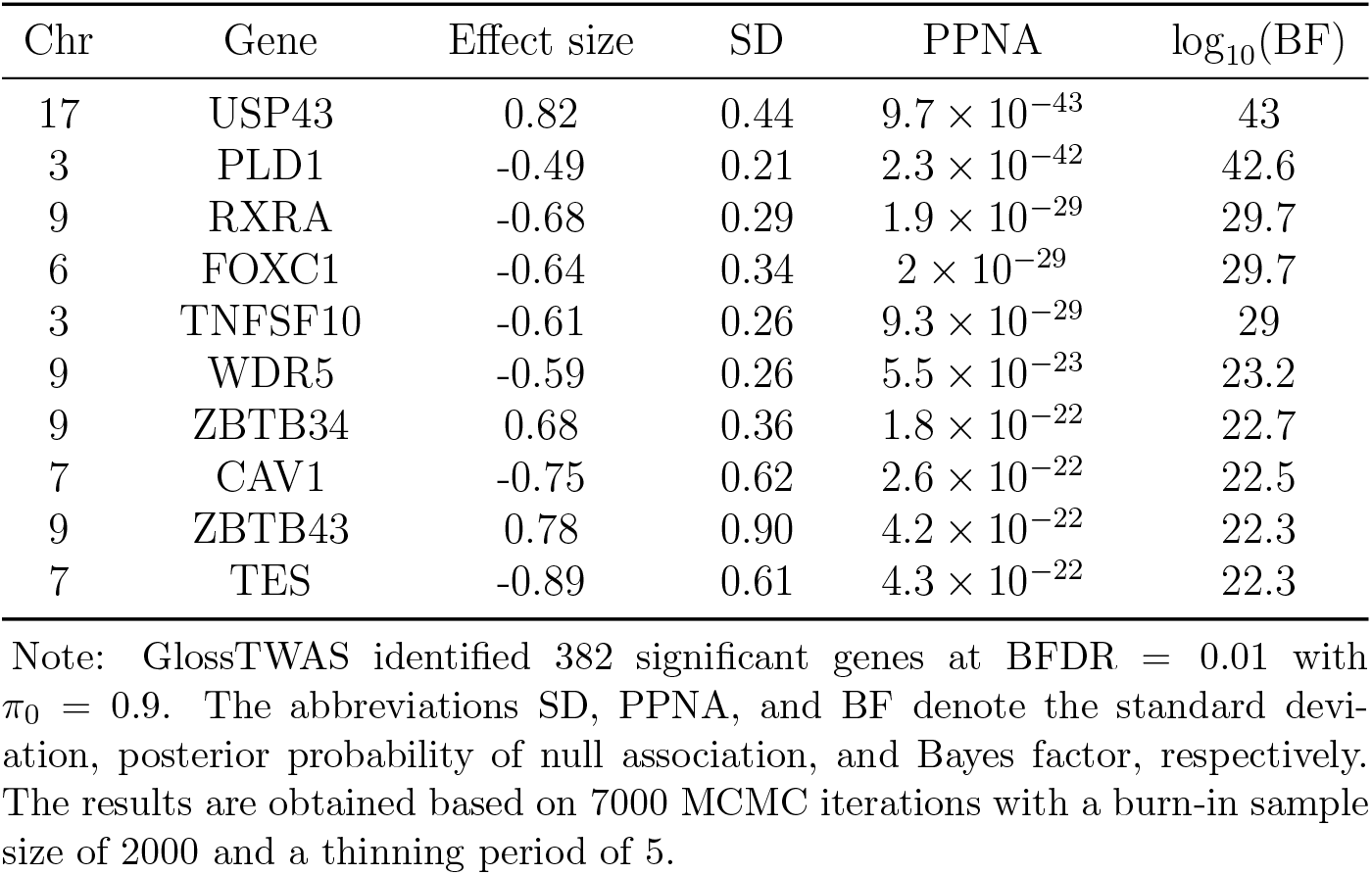
Results for top 10 most significant genes associated with intraocular pressure (IOP) identified by glossTWAS in single-ancestry TWAS for Europeans at BFDR = 0.01.

| Chr | Gene | Effect size | SD | PPNA | $\log_{10}(\text{BF})$ |
| --- | --- | --- | --- | --- | --- |
| 17 | USP43 | 0.82 | 0.44 | $9.7 \times 10^{-43}$ | 43 |
| 3 | PLD1 | -0.49 | 0.21 | $2.3 \times 10^{-42}$ | 42.6 |
| 9 | RXRA | -0.68 | 0.29 | $1.9 \times 10^{-29}$ | 29.7 |
| 6 | FOXC1 | -0.64 | 0.34 | $2 \times 10^{-29}$ | 29.7 |
| 3 | TNFSF10 | -0.61 | 0.26 | $9.3 \times 10^{-29}$ | 29 |
| 9 | WDR5 | -0.59 | 0.26 | $5.5 \times 10^{-23}$ | 23.2 |
| 9 | ZBTB34 | 0.68 | 0.36 | $1.8 \times 10^{-22}$ | 22.7 |
| 7 | CAV1 | -0.75 | 0.62 | $2.6 \times 10^{-22}$ | 22.5 |
| 9 | ZBTB43 | 0.78 | 0.90 | $4.2 \times 10^{-22}$ | 22.3 |
| 7 | TES | -0.89 | 0.61 | $4.3 \times 10^{-22}$ | 22.3 |

Note: GlossTWAS identified 382 significant genes at BFDR = 0.01 with $\pi_0 = 0.9$ . The abbreviations SD, PPNA, and BF denote the standard deviation, posterior probability of null association, and Bayes factor, respectively. The results are obtained based on 7000 MCMC iterations with a burn-in sample size of 2000 and a thinning period of 5.

Among the genes identified by glossTWAS, the USP43 gene on chromosome 17 appeared to be most significantly associated. It has the highest log_10_(BF) = 43 and lowest PPNA = 9.7 *×* 10^−43^ (Table 4). The estimated effect size for this gene is 0.82 with a posterior SD of 0.44. This gene, located in the 17p13.1 region, is reported to be associated with high myopia (difficulty with distant vision) in the NHGRI-EBI GWAS catalog. The next most significant gene is PLD1, located in 3q26.31, with an effect size estimate of −0.47 and posterior SD of 0.21 (Table 4). The same gene was found by PMR-Egger to be associated with the strongest signal of association (*P* = 5 *×* 10^−29^) (Table S18). PMR-Egger estimated the effect size as −0.74. PrediXcan detected the KBTBD4 gene, located in 11p11.2, with the strongest signal of association (*P* = 1.4 *×* 10^−15^). PrediXcan estimated the effect size for KBTBD4 as 1.53 with SD 0.19 (Table S19). GlossTWAS also flagged this gene as associated with log_10_(BF) = 14 and PPNA = 8.3 *×* 10^−14^. However, glossTWAS estimated the gene’s effect size as 0.35 with posterior SD 0.14. PMR-Egger also identified this gene (*P* = 9.3 *×* 10^−11^) with an estimated effect size of 0.29, which is similar to that provided by glossTWAS, but not PrediXcan. For 258 genes among the 382 genes detected by glossTWAS, the magnitude of effect size was estimated to be greater than 0.3. For the remaining 124 genes, it is smaller than 0.3. We found that some genes are positively associated and some are negatively associated.

We note that some genes uniquely identified by glossTWAS are reported in the literature to be associated with relevant traits. For example, the TMCO1 gene on chromosome 1, located in 1q24.1, was previously reported to be associated with IOP and open-angle glaucoma [20]. While PMR-Egger did not show any association with this gene (*P* = 0.99), glossTWAS detected a significant association with log_10_(BF) = 17.6, and PPNA = 2 *×* 10^−17^. Another gene, PTPRJ, located in 11p11.2, was reported to be associated with IOP [21]. PMR-Egger missed this gene (*P* = 0.99), but glossTWAS identified an association with log_10_(BF) = 12.4 and PPNA = 3.7 *×* 10^−12^.

In the multi-ancestry TWAS for Europeans and Africans, glossTWAS identified 342 significant genes with a non-null effect on IOP in either ancestry at 1% BFDR (gene list provided in the supplementary information). On the other hand, METRO detected 304 genes associated with IOP at 1% FDR. The two methods identified a common set of 259 genes. GlossTWAS detected an additional set of 83 genes. Multi-ancestry glossTWAS found the USP43 gene to have the strongest association signal with log_10_(BF) = 41.9 and PPNA = 1.2 *×* 10^−41^ (Table 5). The effect size for the USP43 gene was estimated as 0.59 in EUR ancestry (posterior SD = 0.23) and 0.34 (posterior SD = 0.27) in the AFR ancestry. METRO also found the USP43 gene as the most significant one with a p-value of 2.5 *×* 10^−25^ (Table S21). Both approaches detected the PLD1 gene on chromosome 3 as the second strongest signal. For glossTWAS: log_10_(BF) = 36.2 and PPNA = 5.3 *×* 10^−36^, and the gene’s effect size in the EUR ancestry is −0.47, and AFR ancestry is −0.44 (Table 5).

**Table 5.** Results for top 10 most significant genes associated with intraocular pressure (IOP) identified by multi-ancestry glossTWAS for Europeans and Africans at BFDR level 0.01.

| Chr | Gene | EUR ancestry | | AFR ancestry | | PPNA | $\log_{10}(\text{BF})$ |
| --- | --- | --- | --- | --- | --- | --- | --- |
|  |  | Effect size | SD | Effect size | SD |  |  |
| 17 | USP43 | 0.59 | 0.23 | 0.34 | 0.27 | $1.2 \times 10^{-41}$ | 41.9 |
| 3 | PLD1 | -0.47 | 0.22 | -0.44 | 0.31 | $5.3 \times 10^{-36}$ | 36.2 |
| 9 | RXRA | -0.76 | 0.39 | -0.34 | 0.32 | $2.4 \times 10^{-29}$ | 29.6 |
| 6 | FOXC1 | -1.06 | 0.51 | -0.83 | 0.65 | $1.1 \times 10^{-27}$ | 27.9 |
| 7 | CAV1 | -0.56 | 0.30 | -0.29 | 0.30 | $3 \times 10^{-24}$ | 24.5 |
| 7 | TES | 0.54 | 0.24 | 0.33 | 0.27 | $3.7 \times 10^{-24}$ | 24.4 |
| 9 | ZBTB43 | 0.60 | 0.30 | 0.18 | 0.19 | $5.8 \times 10^{-24}$ | 24.2 |
| 9 | ZBTB34 | 0.68 | 0.51 | 0.17 | 0.23 | $9.8 \times 10^{-23}$ | 23.0 |
| 7 | TFEC | -0.94 | 0.57 | -0.64 | 0.69 | $2.1 \times 10^{-22}$ | 22.6 |
| 9 | WDR5 | -0.53 | 0.26 | -0.41 | 0.37 | $1.3 \times 10^{-21}$ | 21.8 |

Note: GlossTWAS identified 342 significant genes at BFDR = 0.01 with $\pi_0 = 0.9$ . The abbreviations SD, PPNA, and BF represent the standard deviation, posterior probability of null association, and Bayes factor, respectively. The results are obtained based on 7000 MCMC iterations with a burn-in sample size of 2000 and a thinning period of 5.

We observe that the effect sizes for most of the genes identified by multi-ancestry glossTWAS are substantially non-null in both ancestries (Table 5). Moreover, the effect sizes can have different magnitudes across the ancestries. Some of the genes, uniquely identified by glossTWAS, are already reported to be associated with IOP or related traits. For example, GRIK4 on chromosome 11 and FBXL17 on chromosome 5 were found to be associated with IOP by previous studies [22, 23]. METRO did not provide any significant evidence of association for these genes.

Between the single- and multi-ancestry TWAS for IOP, glossTWAS identified an overlapping set of 315 genes. We expect a significant overlap primarily because many genes have an effect in both ancestries in the same direction. On the other hand, the multi-ancestry glossTWAS identified 27 genes that the single-ancestry approach missed. For example, only the multi-ancestry approach spotted the LOX gene on chromosome 5 and the ADGRB2 gene on chromosome 1. For LOX, glossTWAS estimated the European-specific effect size as −0.3, and the African-specific effect size as −0.16. For ADGRB2, glossTWAS obtained the effect sizes as: 0.26 and 0.2, respectively. Previous studies [21, 24] found the LOX and ADGRB2 genes to be associated with IOP. Detection of such associated genes supports the strategy of incorporating multiple ancestries in a TWAS analysis. It helps identify a gene with limited effects across ancestries. We acknowledge that the single-ancestry analysis exclusively identified some genes. Such genes may have an ancestry-specific effect. Alternatively, the substantially lower sample size of the African ancestry-specific data may contribute to this finding. We also observe that the estimated effect sizes of the genes are consistently smaller in magnitude for the African ancestry compared to the European. We attribute this pattern again to the sample size imbalance across ancestries.

We also experimented with *π*_0_ = 0.95 in glossTWAS. For the single-ancestry TWAS, the approach detected 339 genes to be associated (Table S17) compared to 382 genes for *π*_0_ = 0.9. We found fewer genes for *π*_0_ = 0.95 as we assign a smaller prior probability (5%) that a gene is associated, compared to the previous setting (10%). When *π*_0_ = 0.95, glossTWAS identified the same two genes to be most prominently associated with IOP: USP43 in 17p13.1 region as the strongest (log_10_(BF) = 43 and PPNA = 1.9 *×* 10^−42^) and PLD1 in 3q26.31 as the second strongest (log_10_(BF) = 40.4 and PPNA = 6 *×* 10^−40^) signal (Table S17) as we observed with *π*_0_ = 0.9. GlossTWAS detected 329 overlapping genes between *π*_0_ = 0.9 and *π*_0_ = 0.95. For *π*_0_ = 0.95, glossTWAS also identified the TMCO1 and PTPRJ genes. These two genes are discussed above as being uniquely detected by glossTWAS and reported in the literature to be associated with relevant traits. For the multi-ancestry TWAS, glossTWAS obtained 299 genes as noteworthy using *π*_0_ = 0.95, compared to 342 genes for *π*_0_ = 0.9 above. When *π*_0_ = 0.95, multi-ancestry glossTWAS again identified the USP43 and PLD1 genes as the two most significantly associated genes (Table S20).

In the following, we discuss the findings from various analyses that we conducted to validate the gene set for IOP biologically. Since there was substantial overlap between the gene sets identified by the single and multi-ancestry glossTWAS for the datasets analyzed, we summarized the biological validation results only for the gene set obtained from the multi-ancestry glossTWAS.

### 2.5 Functional enrichment analysis

To evaluate the functional relevance of TWAS-associated genes, we performed over-representation analysis (ORA) using g:Profiler (v0.2.3 R package) [25] in R (v4.5.2). We conducted ORA for Gene Ontology Biological Process and Molecular Function categories [26] as well as KEGG [27] and Reactome [28] pathways, using FDR *<* 0.05. To identify the association of TWAS-associated genes with human phenotypes or clinical features, we utilized Human Phenotype (Monarch) enrichment [29] via the STRING web platform [30].

Overrepresentation analysis using g:Profiler R package revealed significant enrichment (FDR *<* 0.05) of pathways involved in regulating metabolism (GO:0051246, GO:0080090, GO:0019538), catabolic processes (GO:0009056), and mRNA translation via miRNAs/ceRNAs targeting PTEN, CDH11, NPAS4 (R-HSA-8948700, R-HSA-8943723, R-HSA-9759811, R-HSA-9768778). We observed additional enrichment in NR1H2/NR1H3 (LXR)–mediated cholesterol transport, efflux (RHSA9024446), and ruffle assembly regulation (GO:1900027). These pathways are consistent with established mechanisms governing intraocular pressure (IOP) homeostasis in the trabecular meshwork (TM), the primary tissue regulating aqueous humor outflow from the anterior chamber of the eye. Structural and compositional changes in the TM extracellular matrix are known to impair aqueous humor drainage, increase outflow resistance, and elevate IOP, a key risk factor for glaucoma [31]. In particular, enrichment of LXR-mediated cholesterol transport pathways underscores the importance of lipid homeostasis in TM function. Cholesterol levels in TM cells regulate membrane composition, actin cytoskeletal dynamics, and cell–extracellular matrix interactions, thereby influencing cellular stiffness and pressure-dependent outflow resistance [32]. Pathways related to WNT signaling, including *β*-catenin–independent WNT signaling (R-HSA-3858494), were also enriched, consistent with the role of WNT pathways in maintaining cadherin-mediated cell–cell junctions within the TM. Disruption of WNT signaling weakens intercellular connectivity, alters cytoskeletal organization, and increases aqueous outflow resistance, leading to elevated IOP [33]. General biological processes, such as localization (GO:0051179) and positive regulation of cellular processes (GO:0048522), were additionally enriched, reflecting coordinated adaptations in intracellular trafficking and proteostasis.

Phenotype enrichment analysis using Monarch Initiative annotations [29] through the STRING web platform [30] revealed significant over-representation of Intraocular pressure measurement and eye measurement as the top hits. Subsequent analysis of the subset of genes associated with these phenotypes identified additional cornea-specific phenotypes including Corneal resistance factor, central corneal thickness, corneal topography, and corneal hysteresis. Collectively, these results highlight its potential relevance to intraocular pressure regulation and corneal biomechanical traits.

### 2.6 Drug analysis

We cross-referenced the TWAS-associated genes with drug-gene interactions to explore therapeutic potential. For this analysis, we obtained drug interaction data from Drugbank [34], parsing the XML file (v5.1.13), and we mapped to TWAS-associated genes to identify potential pharmacological targets.

The integration of TWAS-associated genes with drug-gene interaction data identified around 40 genes that are targets of known pharmacological agents. Notably, PLD1, a gene involved in lipid metabolism, was identified as a target of drugs Choline, LAX-101, and Choline salicylate. Previous studies have reported that PLD1 prevents the inflammatory response of RPE cells and decreases oxidative stress generated in retinal pigment epithelium cells [35]. We provide the complete results in the supplementary information.

## 3 Discussion

We have proposed a novel Bayesian approach to conduct TWAS that circumvents a crucial methodological limitation inherent to standard two-step TWAS. Traditional TWAS methods overlook the uncertainty in predicted expression in the second stage of analysis, which may lead to sub-optimal inference on the gene-trait association. Our proposed method, glossTWAS, overcomes this limitation by jointly modeling the transcriptome and GWAS data in a unified Bayesian framework. Instead of implementing two separate models for transcriptomic and GWAS data, we build a joint Bayesian foundation where a gene’s local SNPs’ effect sizes connect the two seemingly distinct models. We perform a comprehensive Bayesian inference based on MCMC. The simultaneous update of model parameters in the MCMC steps, including the local SNPs’ effect sizes and the corresponding variances, accommodates the uncertainty of the predicted expression in the TWAS analysis. To model the sparse effects of local SNPs on gene expression in transcriptome data, we adapt a Bayesian regularization approach based on a popular global-local shrinkage prior, the horseshoe prior (HSP). The lesser shrinkage of the larger eQTL effects by the HSP is more desirable than the Laplace prior, corresponding to Lasso, which penalizes potent eQTL effects substantially. This property of HSP improves GReX prediction and enhances the overall performance of glossTWAS.

We extend the single-ancestry approach to develop a unified hierarchical Bayesian model for conducting multi-ancestry TWAS combining ancestry-specific expression and GWAS data. Existing methods either use a single GWAS dataset for all ancestries [18] or separate ancestry-specific models for expression and GWAS data, assuming homogeneity in a gene’s effect across all ancestries [17]. Local SNPs can differentially affect gene expression across ancestries due to heterogeneity in eQTL effects, as well as differences in allele frequency and LD pattern of the SNPs. At the same time, we expect some similarity due to the sharing of eQTL effects between the ancestries. We accommodate both possibilities by considering two HSPs for the ancestry-specific eQTL effects of the local SNPs, but with the same local and global shrinkage parameters across ancestries. This setup follows the structure of a hierarchical Bayesian model. A gene can have similar and heterogeneous effect sizes across ancestries, assuming that the effects are in the same direction. Genetic effects on a complex trait across ancestries are strongly correlated. Several GWAS confirmed the similarity between the effect sizes [36]. For example, the FTO gene has been found to be strongly associated with increased BMI [37], whereas the TCF7L2 gene is associated with increased risk of type-2 diabetes across various populations [38]. Primarily, we have developed glossTWAS for two ancestries. Along similar lines, it is feasible to extend the approach for more than two ancestries.

Using extensive simulations, we demonstrate that glossTWAS performs efficiently while classifying null and non-null genes. In most simulation scenarios, glossTWAS yields a higher AUC than the competing approaches based on partial ROC curves. This implies that for a given false positive rate obtained by the methods, glossTWAS offers higher TPR than others. While controlling the Bayesian FDR at a given level, glossTWAS controls the realized FDR below the same threshold. We observe a similar pattern for both single and multi-ancestry TWAS. For the latter, glossTWAS performed robustly well for various extents of eQTL sharing between ancestries. While estimating effect sizes, glossTWAS demonstrates good accuracy, surpassing that of the competing approaches.

In multi-ancestry TWAS, METRO does not explicitly model separate GWAS data for each ancestry, unlike glossTWAS. Despite this major difference, we compared glossTWAS with this approach mainly to validate our findings. In addition, METRO models the transcriptome and GWAS data simultaneously. Ancestry-specific distinct GWAS studies are commonly conducted across global populations. Meta-analyses are widely performed to combine these genetic studies. Therefore, our multi-ancestry TWAS setup is relevant and useful in practice. Recent methods [17] have also considered such ancestry-specific GWAS data.

We implemented glossTWAS for single and multi-ancestry TWAS of intraocular pressure. The latter combined TWAS data for Europeans and Africans. We identified genes that affect the variation of IOP, the primary biomarker for glaucoma. We demonstrate that the gene sets are enriched in biological pathways relevant to eye pressure, highlighting the potential of glossTWAS. Future studies can investigate these genes in more detail to gain a better understanding of the biology of eye pressure.

We discuss a few limitations of our approach and future directions of work. First, we considered that local SNPs around the gene affect the outcome trait only by modulating the gene’s expression level. However, for some genes, local SNPs can influence the trait without affecting gene expression, known as horizontal pleiotropy. In future work, we aim to extend glossTWAS to account for horizontal pleiotropy. Second, we developed the model to analyze continuous outcomes. Binary disease traits are crucial, and we plan to extend the approach to analyze binary outcomes. Third, in the absence of individual-level data, we aim to develop the approach based on summary-statistics data, which we are currently working on. Fourth, we plan to extend the multi-ancestry model to accommodate more than two ancestries, allowing us to analyze multiple populations, such as Europeans, Africans, East Asians, and South Asians, simultaneously. We will apply glossTWAS to a wider range of outcome traits and transcriptome datasets of larger sample sizes. We will also apply this approach to other ancestries, such as South-Asian ancestry.

Overall, glossTWAS is a novel unified hierarchical Bayesian approach to performing TWAS. The method is based on Bayesian priors that have attractive properties. It is an efficient and powerful approach for a single and multi-ancestry TWAS. The method can be applied to identify genes associated with various complex traits and to gain a deeper understanding of their genetic basis.

## Supporting information

Link for Supplementary Material

## 4 Acknowledgments

We sincerely thank Rahul Kumar for helpful discussions related to this work. Arunabha Majumdar was supported by the Anusandhan National Research Foundation (ANRF) advanced research grant with an identification number ANRF/ARG/2025/005607/MS. This research used the UK Biobank resource under application 77327.

## Data and software information

The dataset and software used in this article are available (either openly or via application) on the following websites.

Geuvadis data: https://www.internationalgenome.org/data-portal/data-collection/Geuvadis.

UK Biobank: https://www.ukbiobank.ac.uk/

HAPGEN: https://mathgen.stats.ox.ac.uk/genetics_software/hapgen/hapgen2.html

PLINK: https://www.cog-genomics.org/plink/

## Declaration of interests

The authors have no competing interests.

## 5 Materials and methods

### 5.1 Single-ancestry TWAS

Consider a gene and its expression in the tissue or cell type relevant to the outcome trait. Suppose the reference transcriptome data comprises *n* individuals. Let **x**_ref_ denote the vector of gene expression levels for *n* individuals. We consider *m* local SNPs surrounding the gene’s transcription start and end sites. Let *G*_ref_ denote the *n × m* genotype matrix for the local SNPs for the individuals in the transcriptome data. We consider the following linear model regressing the expression on the local SNPs’ genotypes:

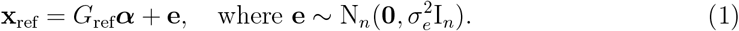

We do not include an intercept term since the vectors of gene expressions and genotypes for each local SNP are standardized. Here, ***α*** is the vector of *m* local SNPs’ effects on the gene expression level. Each component of the random error vector **e** independently follows a normal distribution with mean 0 and common variance 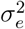.

Next, we consider separate GWAS data that comprises *N* individuals. Let **y** denote the vector of outcome trait values for the *N* individuals. Let *G* stand for the *N × m* genotype matrix for *m* SNPs in the GWAS data, where these SNPs are the same local SNPs considered in the transcriptome data. To model the relation between the trait and the genetically regulated component of expression for the GWAS individuals, we consider another linear model:

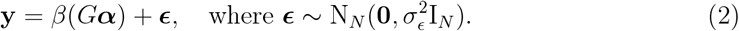

We do not include an intercept term, considering **y** to be mean-centered. The effect of local SNPs on the expression, ***α***, remains the same between the models based on the transcriptome (Equation 1) and GWAS (Equation 2) data. These two datasets are assumed to be ancestry-matched. *G****α*** in the Equation 2 represents the true genetically regulated component of expression (GReX) for the GWAS individuals, and *β* denotes the effect of true GReX on the outcome trait. 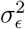 is the common error variance for each component in the residual vector ***ϵ***. Since the individuals are non-overlapping between the transcriptome and GWAS data, **e** and ***ϵ*** (Equations 1 and 2) are independently distributed. The above two linear models are connected by the common parameter vector ***α***. We use the horseshoe prior for ***α***, which has the following structure:

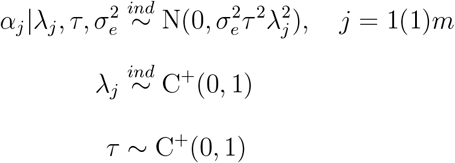

Here, C^+^(0, 1) is the standard half-Cauchy distribution with its support on the positive real numbers. Two distinct kinds of shrinkage parameters are considered to regulate the overall sparsity. For *j* = 1, …, *m, λ*_*j*_ is the local shrinkage parameter specific to the effect of the *j*^*th*^ local SNP (*α*_*j*_), and *τ* is the global shrinkage parameter. *τ* applies an overall shrinkage to the effects toward zero due to the sharp peak of the prior near the origin. *λ*_*j*_s help local SNPs with sizable effect sizes escape overall shrinkage, which is possible due to the prior’s heavy Cauchy-like flat tail [12].

To evaluate the impact of the GReX on the outcome trait, we employ the Dirac spike and slab prior for *β* (Equation 2), the effect of the GReX on the trait, as:

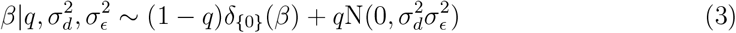

*δ*_{0}_(*β*) is the point mass function that has a discrete measure if *β* = 0 and 0 when *β*≈ 0. The spike component constitutes the null effect, whereas the slab component represents the non-null effect of the GReX on the outcome. *q* is the prior probability that *β* is non-null. We consider a Beta prior for *q* as: *q* ∼ Beta(*b*_1_, *b*_2_). Here, 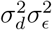 denotes the slab variance for the GReX effect size *β*. The slab variance is proportional to the trait variance 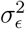, with 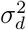 as the ratio between them. Thus, 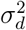 is a crucial component of the prior variability of *β*. In the prior, we consider inverse gamma (IG) distributions for the variances 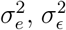, and 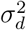. Collating the layers (Equations 1 and 2), we model the transcriptome and GWAS data together in a unified and multi-layered Bayesian framework as:

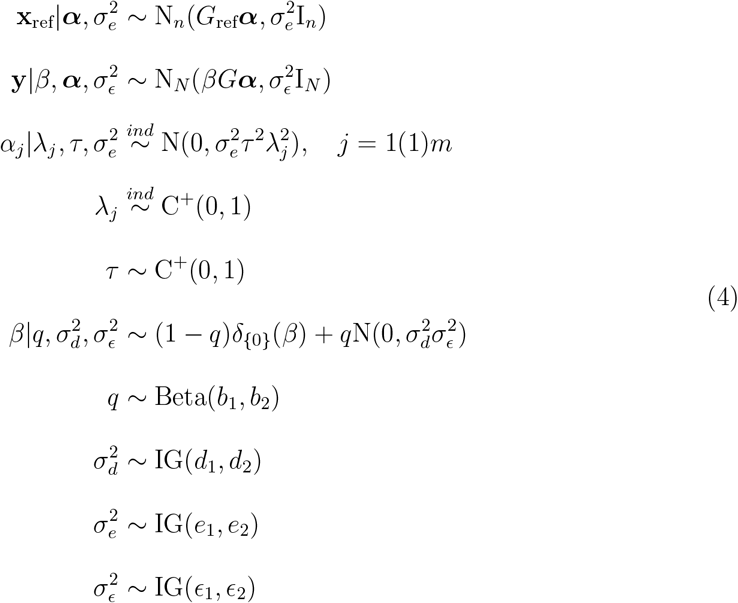

We implement the Markov chain Monte Carlo (MCMC) algorithm to generate posterior samples of the model parameters and perform a comprehensive Bayesian inference. We devise a Metropolis within Gibbs sampler to obtain the posterior samples. The detailed derivations of the full conditional distributions are presented in the supplemental materials.

### 5.2 Estimation and testing

In glossTWAS, *β* denotes the effect of GReX on the trait (Equation 2). We consider the posterior mean of *β* using the MCMC sample as the estimate of *β* and calculate the posterior standard deviation (SD) based on the sample to assess its uncertainty. We regard the null hypothesis *H*_0_ : *β* = 0 versus *H*_1_ : *β*≈ 0. We design a Bayesian testing strategy based on the Bayes factor and posterior probability of null association. Let *D* denote the complete data, including the gene expression and outcome trait in the transcriptome and GWAS data, respectively. The posterior odds of *H*_1_ versus *H*_0_ is 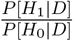 and the prior odds is 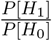. The Bayes factor (BF) for testing *H*_1_ versus *H*_0_ is given by:

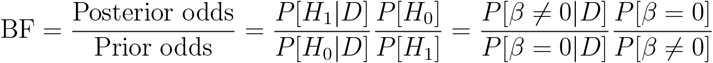

*P* [*H*_0_|*D*] = *P* [*β* = 0|*D*] stands for the posterior probability of null association (PPNA). The higher the value of the Bayes factor, the stronger the evidence against *H*_0_. The lower the value of PPNA, the stronger the evidence against *H*_0_. The expression of prior odds of *H*_1_ versus *H*_0_ is 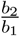. For the computation of posterior odds, we estimate the PPNA following the Rao-Blackwellization technique [39] as:

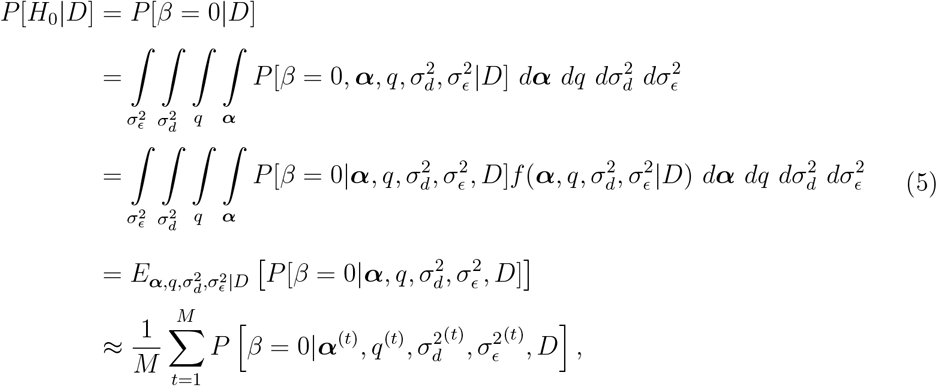

where 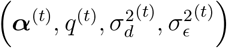 denotes the *t*^th^ posterior draw of (***α***, *q*, 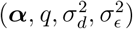) in the MCMC and *M* is the number of MCMC iterations. The term 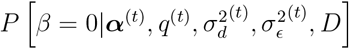 evaluated at the *t*^th^ step of MCMC is the full-conditional posterior probability that *β* = 0.

### 5.3 Hyperparamters

We discuss the choice of hyperparameters in the Bayesian model of glossTWAS. The prior distribution of *q* is Beta(*b*_1_, *b*_2_) with the shape parameters *b*_1_ and *b*_2_. We note that 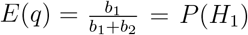, the prior probability that the gene is associated with the outcome trait. Since we expect only a small proportion of genes to be associated at a genome-wide level, we regard *P* (*H*_1_) = 0.1. For simplicity, we fix *b*_1_ = 1. We choose *b*_2_ such that *P* (*H*_1_) = 0.1. For this choice of (*b*_1_, *b*_2_), *P* (0 *< q <* 0.28) = 0.95. Next, we discuss the choices of the hyperparameters for the inverse gamma priors of the variance terms. Following previous literature [40], we consider flat and weakly informative priors for 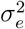 and 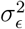. We choose *e*_1_ = *e*_2_ = *ϵ*_1_ = *ϵ*_2_ = 0.1. To determine the hyperparameters for the prior of 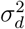, we adapt the method of moments approach implemented by Majumdar et al. [41]. Specifically, we equate 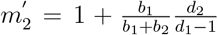, where 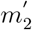 is the second-order sample raw moment of the *Z* (Wald) statistic for a standard TWAS method, e.g., PrediXcan, across all genes for an outcome trait. The right side of the equality is *E*(*Z*^2^) under a spike and slab prior assigned to a standardized effect size (multiplied by the square root of the GWAS data sample size) of the gene on the outcome [41]. Along with this equation, we further restrict the hyperparameters *d*_1_ and *d*_2_ such that the mode of the IG(*d*_1_, *d*_2_) distribution coincides with the mode of a weakly informative IG(0.1, 0.1) distribution. Besides the method of moments approach, we also consider the mode of the weakly informative IG prior following recent studies [40]. The mode of IG(0.1, 0.1) is 0.09. These provide two equations to solve to find *d*_1_ and *d*_2_. We obtain *d*_1_ = 1.003 and *d*_2_ = 0.18. This choice provides 90% prior probability that 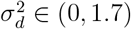. Thus, the hyper-parameter choices turn out to be in a sensible range. We note that sufficiently large sample sizes of TWAS data help achieve robustness concerning the hyper-paramer choices.

### 5.4 Multi-ancestry TWAS

We extend the single-ancestry glossTWAS framework to the multi-ancestry setup (Figure 12). We initially developed the model for two ancestries. We assume that a pair of transcriptome and GWAS data is available for each of the two ancestries. We consider a single gene and its expression in the same tissue or cell type relevant to the outcome trait in both ancestries. Denote **x**_*k*,ref_ as *n*_*k*_ *×* 1 vector of gene expression levels measured for *n*_*k*_ individuals in the *k*^th^ ancestry, *k* = 1, 2. Let *G*_*k*,ref_ denote *n*_*k*_ *× m* local SNPs’ genotype matrix in the transcriptome data for *k*^th^ ancestry, where *m* is the number of local SNPs. We consider the linear model for the ancestry-specific transcriptome data as follows:

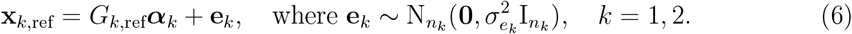

**Figure 11.**
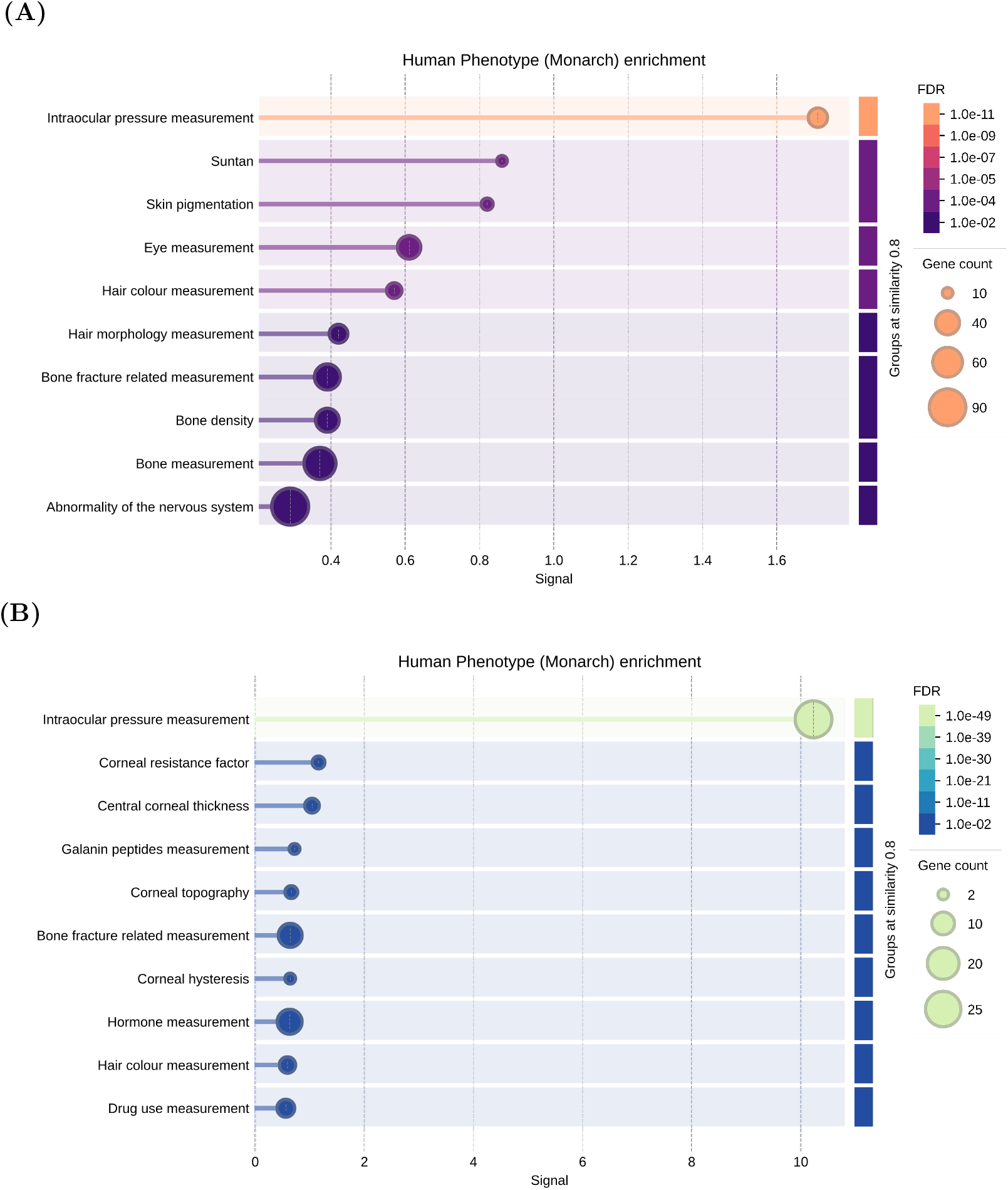
Human phenotype enrichment of multi-ancestry glossTWAS genes using Monarch Initiative annotations performed through the STRING web platform. (**A**) Top 10 phenotypes from Phenotype enrichment analysis using the complete set of multi-ancestry glossTWAS genes reveal significant over-representation (FDR *<* 0.05) of ocular phenotypes, with intraocular pressure measurement and eye measurement emerging as the top enriched terms. (**B**) Subnetwork analysis of genes associated with the top-enriched phenotype (intraocular pressure measurement) identifies additional, more specific cornea-related phenotypes.

**Figure 12.**
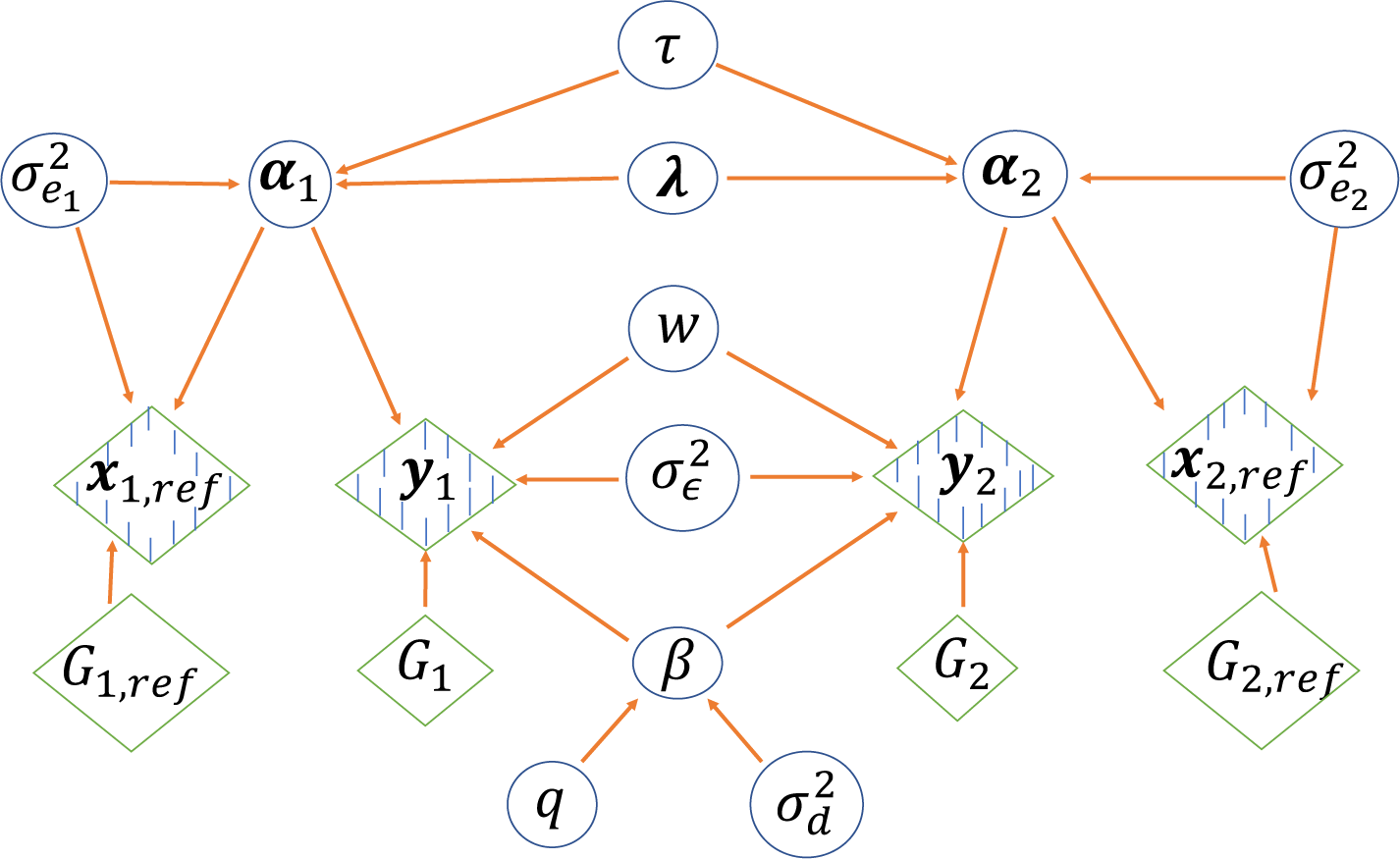
Graphical presentation of the hierarchical Bayesian model for glossTWAS in the multi-ancestry TWAS framework. Shaded boxes denote observed random variables, unshaded boxes indicate covariates, and the circles represent the unobserved random variables. Arrows denote the conditional dependencies between variables.

Here, ***α***_*k*_ represents the ancestry-specific effect sizes of the local SNPs on expression, and 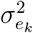 is the error variance for each component of the residual vector **e**_*k*_. We include different ***α***_*k*_s since true eQTLs can vary across populations with possibly distinct effect sizes. We also accommodate possible sharing of eQTL effects between ancestries, using a hierarchical Bayesian structure (discussed below). Since the heritability of gene expression due to local SNPs can differ among ancestries, we consider ancestry-specific error variance terms 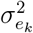.

Suppose **y**_*k*_ is the vector of outcome trait values for *N*_*k*_ individuals in the GWAS data for *k*^th^ ancestry, *k* = 1, 2. Let *G*_*k*_ stand for *N*_*k*_ *× m* local SNPs’ genotype matrix in the GWAS data for *k*^th^ ancestry. The local SNPs are consistent across transcriptome and GWAS data, as well as the ancestries. We model the gene-outcome trait relationship for the *k*^th^ GWAS data as:

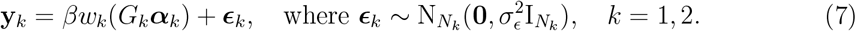

We consider: *w*_1_ *>* 0, *w*_2_ *>* 0, and *w*_1_ + *w*_2_ = 1. The term *G*_*k*_***α***_*k*_ represents the true GReX in the *k*^th^ ancestry. The ancestry-specific GReX effect size on the outcome is *βw*_*k*_. Here, *β* ∈ ℝ is the common effect size parameter that is multiplied by *w*_*k*_ to specify the *k*^*th*^ ancestry effect of the gene on the outcome. We adapt this setup inspired by the METRO method proposed in Li et al. [18]. *w*_*k*_ in Equation 7 reflects the relative contribution of the *k*^th^ ancestry to constructing the overall effect of the gene. We consider a spike and slab prior for *β* (described in the following). When *β* = 0, the gene is not associated with the outcome in both ancestries, which corresponds to the null hypothesis. When *β*≈ 0, various possibilities of *w*_1_, *w*_2_ allow different scenarios, such as similar effects of the gene across ancestries (*w*_1_ ≈ *w*_2_), unequal effects, limited effect in one ancestry (e.g., *w*_1_ ≈ 0), etc. Thus, if glossTWAS rejects a null hypothesis, the gene’s association may be mainly driven by one ancestry or both. Since a single gene explains a tiny proportion of the variability of a complex trait in each ancestry, we assume a common error variance 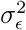.

We employ two horseshoe priors for ***α***_*k*_s in the two ancestries, but with the same shrinkage parameters across ancestries, implementing a hierarchical Bayesian framework:

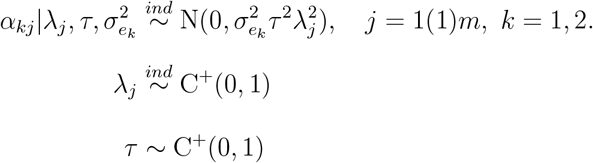

For *j* = 1, …, *m, α*_*kj*_ is the effect of the *j*^th^ local SNP on the gene expression in the *k*^th^ ancestry. We introduce a common local SNP-specific shrinkage parameter, *λ*_*j*_, which is independent of the ancestry. This implies a similar shrinkage of the local SNPs’ effect sizes on gene expression over the ancestries. We also regard the same global shrinkage parameter *τ* across the ancestries. The same *λ*_*j*_, *j* = 1, …, *m*, and *τ* authorize the possible sharing of eQTLs and a similarity between their effect sizes across populations. We note that the sharing is induced weakly because we assumed independence of the effect size distributions across the ancestries. The choice of the same shrinkage parameters across ancestries frames a hierarchical Bayesian model.

We again employed the Dirac spike and slab prior (Equation 3) to evaluate the overall effect of the gene on the trait, *β*:

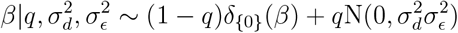

We denote *w*_1_ = *w*, hence *w*_2_ = 1 −*w*. We regard Beta(*γ*_1_, *γ*_2_) prior for *w* with *γ*_1_ = *γ*_2_ = 1, leading to a Uniform(0,1) prior distribution. As any information on *w* is generally not available a priori, a Uniform prior distribution ensures that *w* will be inferred based on the data. For the error variance terms, we consider inverse gamma priors. A complete hierarchical Bayesian structure of the multi-ancestry glossTWAS model is given by:

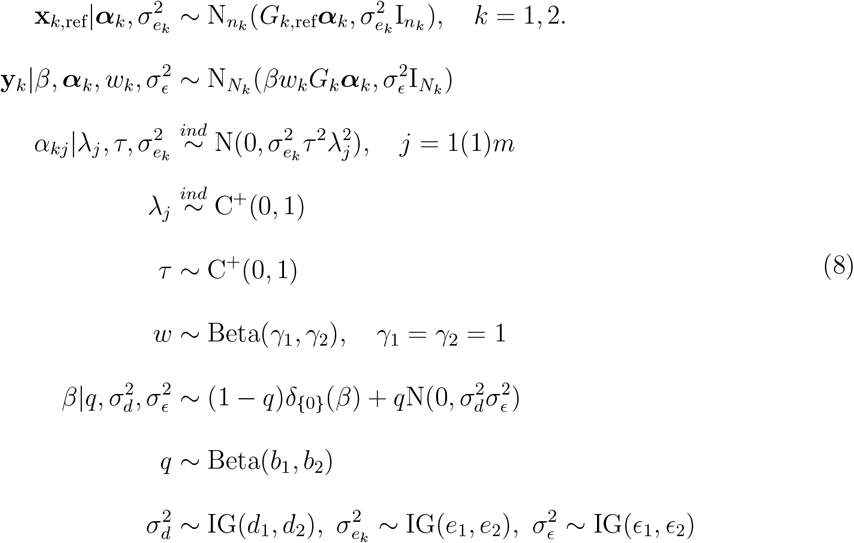

We perform a comprehensive Bayesian inference by implementing the MCMC based on the Metropolis-within-Gibbs sampler. The outline of the algorithm and the derivations of the full conditional posterior distributions are presented in the supplemental materials. The hyper-parameters in the multi-ancestry model are chosen analogously to single-ancestry TWAS (section 5.3). We primarily develop the multi-ancestry TWAS model for two ancestries. The method can be generalized to analyze more than two ancestries by adapting a Dirichlet prior for the vector of ancestry-specific weights *w*_1_, …, *w*_*k*_.

### 5.5 Estimation and testing

The estimation procedure remains similar for the multi-ancestry approach. We use the posterior means to estimate the ancestry-specific GReX effect sizes and the posterior SDs to assess their uncertainties. We evaluate *H*_0_ : *βw*_1_ = *βw*_2_ = 0 vs. *H*_1_ : *βw*_1_ ≈ 0, *βw*_2_ ≈ 0. It is equivalent to *H*_0_ : *β* = 0 versus *H*_1_ : *β*≈ 0, since 0 *< w*_1_, *w*_2_ *<* 1 and *w*_1_ + *w*_2_ = 1. Thus, *H*_0_ implies that the gene has no effect on the outcome trait in both ancestries. Although in the alternative hypothesis *H*_1_, the gene has a non-null effect in both ancestries in the same direction, a very small positive *w*_1_ will reflect a negligible effect in the first ancestry. Similarly, other combinations of (*w*_1_, *w*_2_) will characterize various patterns of the relative extent of the effect across ancestries. Also, a small |*β*| can reflect limited effects of the gene in both ancestries. Thus, if we find a gene to be associated, it may be due to substantial effects in both ancestries, moderate effects in both, or prominent in one but negligible in the other, depending on the relative values of *w*_1_ and *w*_2_. Hence, we interpret *H*_1_, the alternative hypothesis, as an association of the gene in either or both ancestries. We employ a Bayesian testing strategy and compute the Bayes factor and PPNA for the above hypothesis. We adopt a similar procedure, likewise the single-ancestry framework, to compute the prior and posterior odds (details provided in the supplementary materials).

### 5.6 Simulation Design

We evaluate the performance of our method, glossTWAS, by conducting extensive simulation studies and comparing it with some other existing approaches. We regard the accuracy of effect size estimation and the correct identification of null and non-null genes as the two main aspects of evaluation. For single-ancestry TWAS, we compare glossTWAS with the PMR-Egger [14], TIGAR [11], and PrediXcan [3]. For multi-ancestry TWAS, we contrast glossTWAS with METRO [18]. METRO does not explicitly model separate GWAS data for two ancestries. We combine the transcriptome datasets for two ancestries and run METRO for each GWAS data to get ancestry-specific p-values. Then, we combine the p-values implementing the aggregated Cauchy association test (ACAT) [42] to perform the multi-ancestry TWAS in a meta-analysis style. We choose ACAT primarily because the p-values can be dependent.

We consider the genetic region surrounding the DGCR8 gene located on chromosome 22. For single-ancestry TWAS, we chose the Japanese population from the 1000 Genome Project, which serves as a template population for simulating the genotype data. Surrounding DGCR8, we retain only those SNPs in the analysis that fall within the 500 kilobases of the upstream and downstream regions of the gene. We randomly select 300 SNPs from this set of local SNPs with minor allele frequencies (MAF) more than 0.05. To simulate genotype data for the individuals in the reference transcriptome data, we implement the HAPGEN software [43], which mimics the real LD pattern of the local SNPs in the Japanese population. HAPGEN is an established method based on meticulous theory for simulating genotype data that resembles a chosen real-world population. We run HAPGEN twice to simulate genotype data for the individuals in the transcriptome and GWAS data.

For *i*^th^ individual in the transcriptome data, we generate gene expression using the linear model:

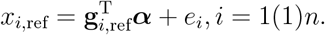

Here, *x*_*i*,ref_ is the expression level, and **g**_*i*,ref_ is the vector of genotype values at the local SNPs for the *i*^th^ individual. ***α*** denotes the effect size vector for local SNPs, and *e*_*i*_ is the random residual. Under the assumption that Var(*x*_*i*,ref_) = 1, we simulate *e*_*i*_ from 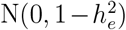, where 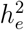 denotes the heritability of expression due to the local SNPs. We assume that 10% of local SNPs affect the gene expression. Let *m*_*c*_ stand for the number of eQTLs affecting the expression. The non-zero elements of ***α*** are generated from 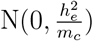. We assign zero value to the remaining components of ***α***.

Next, we discuss the steps to simulate the complex trait for the individuals in GWAS. The GWAS individuals do not overlap with those in the reference transcriptome data. HAPGEN generates the genotype data for the GWAS individuals for the same set of local SNPs considered in the transcriptome data. Let **g**_*i*_ denote the genotype vector for the *i*^th^ individual in the GWAS data, *i* = 1(1)*N* . Assuming that the effect of local SNPs on expression remains the same in both datasets, we first simulate the gene expression for the *i*^th^ GWAS individual using the linear model:

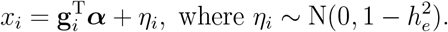

*x*_*i*_ is the gene expression level for *i*^th^ individual in GWAS. We generate the outcome trait in GWAS employing the following linear model:

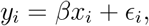

where *y*_*i*_ is the value of the trait for *i*^th^ individual and *β* denotes the effect of gene expression on the trait. Plugging in the linear model for *x*_*i*_ in the above equation, we can view *β* as the effect of the true GReX on the outcome. We simulate the random error term *ϵ*_*i*_ from N(0, 1). We assume Var(*y*_*i*_) = Var(*ϵ*_*i*_) = 1 because one gene is likely to explain a negligible proportion of the trait’s variance. We choose different non-zero values of *β* to create various non-null simulation scenarios. We standardize the vectors of gene expression, GWAS trait, and the columns of genotype matrices in both transcriptome and GWAS panels.

For multi-ancestry TWAS, we follow the simulation setup described in Li et al. [18]. We select the Japanese and British populations from the 1000 Genome data, which serve as the representative populations for East Asian Ancestry (EAS) and European Ancestry (EUR), respectively. We employ HAPGEN to simulate genotype data for the 300 local SNPs (MAF *>* 0.05) for the same DGCR8 gene for each ancestry. We implement the steps described above for a single-ancestry setup to simulate gene expression in separate transcriptome data for the two populations. We use the following linear model:

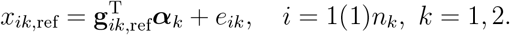

The notations *x*_*ik*,ref_ and 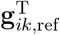 denote the gene expression level and genotype vector for *i*^th^ individual in the transcriptome data for *k*^th^ ancestry. ***α***_*k*_ is the vector of local SNPs’ effects on expression, and *e*_*ik*_ is the random noise specific to *k*^th^ ancestry. We assume Var(*x*_*ik*,ref_) = 1 and simulate *e*_*ik*_ from 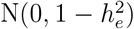 where 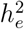 is the expression heritability. We consider 10% of the local SNPs to have a non-null effect on expression in each ancestry. The elements of ***α***_*k*_ that correspond to the non-null SNPs are simulated from 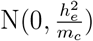 where *m*_*c*_ stands for the number of causal SNPs. We regard a fixed proportion, *f*_0_, of causal SNPs to be common between the two ancestries, leading to eQTL sharing or overlap. We obtain the gene expression level for *i*^th^ individual in GWAS in *k*^th^ ancestry using the linear model:

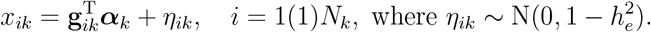

We generate the outcome trait for *N*_*k*_ GWAS individuals in *k*^th^ ancestry as follows:

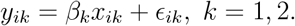

*β*_1_ and *β*_2_ denote the effects of the gene expression, equivalently of the GReX, on the outcome trait in EAS and EUR ancestries, respectively. We simulate *ϵ*_*ik*_ from N(0, 1) since a single gene explains a negligible proportion of the trait’s variance. We vary *β*_1_, *β*_2_ and *f*_0_ to create different simulation scenarios.

For single-ancestry TWAS, we choose the transcriptome data sample size as *n* = 500 and the GWAS data sample size as *N* = 10000. We consider both positive and negative values for the GReX effect size (*β* = 0.1, 0.2, 0.3, 0.4, 0.5 and *β* = −0.1, −0.2, −0.3, −0.4, −0.5). We fix the gene expression heritability, 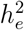, at 10%. In multi-ancestry TWAS, we set the sample sizes of the transcriptome data for EAS and EUR ancestries to be *n*_1_ = *n*_2_ = 500. We choose the corresponding GWAS data sample sizes as *N*_1_ = *N*_2_ = 10000. We make two specific choices of *f*_0_ as 0 and 50%. We regard equal (*β*_1_ = *β*_2_ = 0.1, 0.2, 0.3, 0.4, 0.5) and distinct (*β*_1_ = 0.1, *β*_2_ = 0.2 and *β*_1_ = 0.1, *β*_2_ = 0.5) GReX effect sizes for the two ancestries. We also consider the sign of the effect sizes to be negative for the above choices. We choose the same expression heritability, 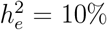, in two ancestries. We obtain posterior summaries for glossTWAS based on a sample of size 1000 obtained from 7000 MCMC iterations with a burn-in period of 2000 and a thinning period of 5.

To assess the accuracy of identifying non-null genes, we compare glossTWAS with PMR-Egger, TIGAR, and PrediXcan in the single-ancestry setting and with METRO in the multi-ancestry setting. Following the same procedure described in Coram et al. [44], we measure the accuracy by computing two quantities: false positive rate (FPR) and true positive rate (TPR). We define the FPR as the proportion of null genes declared significant, whereas the TPR denotes the proportion of non-null genes that are found to be associated with the outcome trait. Here we consider a set of 200 genes selected at random from chromosome 9. We assume 5% of these genes affect the trait. We simulate the effect sizes for the non-null genes from 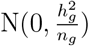, where 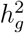 denotes the trait heritability due to the GreX of the non-null genes and *n*_*g*_ is the number of non-null genes. In single ancestry TWAS for each gene, we test whether it has any effect on the trait. We perform gene-trait association for each of these 200 hypotheses for each method. We obtain the p-values for classical approaches and the Bayes factor and PPNA for glossTWAS. In multi-ancestry TWAS, we test whether the gene affects the trait in either or both ancestries. In the simulation setting, we consider that half of the non-null genes have an effect on the outcome in the first ancestry, and the remaining in the second.

For a frequentist method, we arranged the p-values in increasing order. For glossTWAS, we arranged the log_10_(BF) in decreasing order. Then, we reject the null hypotheses for the corresponding ordered genes sequentially based on varying choices of significance thresholds of log_10_(BF) (p-values). For such a sequence of thresholds, we enumerate the actual number of correct and false rejections. For a given method, we calculate the FPR and TPR for each threshold and plot to create a partial receiver operating characteristic (ROC) curve. For a valid comparison, we vary the p-value and Bayes factor thresholds incrementally such that the competing methods obtain the same estimate of FPR. Then, we obtain the TPR of the methods for the same FPR. We estimate the area under the curve (AUC) for each method, and a method with a higher AUC is preferred. We implement this strategy because the p-value and Bayes factor are not directly comparable. We consider four different combinations of 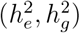 as (10%, 10%), (10%, 20%), (20%, 10%), and (20%, 20%) to create different simulation scenarios for the partial ROC curves.

In addition to partial ROC curves, we simultaneously check whether these methods control the error rate for testing multiple hypotheses in the above simulation scenarios. We employ two well-known measures: false discovery rate (FDR) for frequentist approaches, and Bayesian false discovery rate (BFDR) [45] for glossTWAS to assess the respective methods. FDR is defined as the expected proportion of false rejections among all rejections of null hypotheses. To control FDR at a pre-specified level, we adopt the Benjamini-Hochberg FDR controlling procedure [19] for the classical approaches. BFDR is defined as the expected proportion of false discoveries among all rejected null hypotheses conditioned on the observed data, where the expectation is taken with respect to the posterior distribution of the hypotheses’ truth or falsity [45]. To control BFDR for glossTWAS, we first compute the posterior probability of null association (PPNA) for each hypothesis. Suppose, we have *r* hypotheses and let *p*_*i*_ = *P* (*H*_0*i*_|*D*), *i* = 1, 2, · · ·, *r*. We order the *p*_*i*_s from smallest to largest and let *p*_(*i*)_ is the *i*^th^ smallest element in (*p*_1_, *p*_2_, · · ·, *p*_*r*_). To control BFDR at the level *α*, we reject all hypotheses associated with *p*_(1)_, *p*_(2)_, · · ·, *p*_(*l*)_, where *l* is the largest integer such that 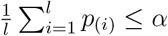. We regard the prior probability of *H*_0*i*_ being true as 0.9. We perform simulation studies for varying thresholds of FDR and BFDR, e.g., 0.01, 0.05, 0.1, etc, with varying choices of expression and trait heritability.

When comparing the accuracy of effect size estimation by glossTWAS with PMR-Egger, TIGAR, and PrediXcan, we base the comparison on a subset of effective replications within a given simulation scenario. Specifically, we compute log_10_(BF) for glossTWAS and the p-values for classical methods for testing *H*_0_ : *β* = 0 vs *H*_1_ : *β*≈ 0 in each replication. We include only those replications for which these methods show an association signal at a nominal level, i.e., p-value *<* 0.05 and log_10_(BF) *>* 1*/*2 [46]. We consider such replications because effect size estimation is of interest in settings where the gene shows at least a nominal evidence of association. In the multi-ancestry TWAS setup, we employ the same strategy to compare glossTWAS with METRO. We consider 300 such effective replications.

### 5.7 Real Data Application

We apply our approach, glossTWAS, to conduct a single-ancestry and multi-ancestry TWAS for intraocular pressure (IOP), which is also known as eye pressure. IOP is a measurement of the fluid pressure inside the eye. It is measured in units of millimeters of mercury (mmHg). An IOP measurement between 10 and 20 mmHg is considered normal. Too low or too high IOP can damage an individual’s vision. Elevated IOP without other symptoms is called ocular hypertension. Some people can have higher eye pressure with no damage. Other people may lose vision even if the pressure is in the normal range. When an individual has glaucoma, eye pressure damages the optic nerve, resulting in permanent vision loss. Without treatment, glaucoma can lead to complete blindness.

We obtain the reference transcriptome data from the Geuvadis expression study [47] and the GWAS data from the UK Biobank (UKB). RNA sequencing was performed for lymphoblastoid cell lines in the Geuvadis data, comprising 462 individuals from the 1000 Genomes Project. The mRNA expression levels are measured for individuals belonging to five different populations: CEPH (CEU), Finns (FIN), British (GBR), Toscani (TSI), and Yoruba (YRI). We combine the individuals from CEU, FIN, GBR, and TSI to form a broader European (EUR) ancestry with a sample size of 373. The remaining 89 individuals in the YRI population constitute an African (AFR) ancestry. We consider the EUR ancestry to perform a single-ancestry TWAS and the EUR and AFR ancestries to conduct a multi-ancestry TWAS for IOP.

We remove those genes from the analysis for which gene expression measurements are missing in at least 50% of the individuals. Among the remaining genes, we implement the KNNimpute algorithm [48] to impute the missing expression values for a gene. We annotated the protein-coding genes from the GENCODE (release v44) and retained a final set of 12517 genes, common with the Geuvadis data. Using multiple linear regression, the expression levels of a gene are adjusted for covariates such as age, sex, and the top 10 genetic principal components (PCs) separately within each ancestry, EUR and AFR. The genetic PCs are included to adjust for the effect of population stratification on the expression. We then apply the rank-based inverse normal transformation [49] on the adjusted expression so that the transformed residuals satisfy the normality assumption in the transcriptome data. We follow the standard quality control (QC) procedure to retain good-quality SNPs. We implement the following QC steps for the genotype data of the EUR and AFR ancestries separately. We filter out SNPs with MAF *<* 0.05. We test for the Hardy-Weinberg equilibrium (HWE) and remove the SNPs with p-value *<* 5 *×* 10^−8^. We also removed SNPs with a genotype missingness rate of more than 10%.

We obtain the GWAS genotype and trait data for IOP from the UK Biobank. We considered the Goldmann-correlated IOP measurement of the right eye (UKB Field ID = 5255). IOP is measured in units of millimeters of mercury (mmHg) in the UK Biobank. The two populations, White British (WB) and Black British (BB), constitute GWAS data for EUR and AFR ancestries in UKB. We remove a portion of the GWAS individuals in either ancestry who do not meet certain criteria. Specifically, we exclude individuals whose self-reported sex does not match their genetically determined sex, those not involved in the computation of genetic PCs available on the UKB research analysis platform (RAP), and those who have a genetic relatedness to any other participants. We finally retain 62147 unrelated individuals from the WB population and 3083 people from the BB population to analyze the phenotype IOP. Using multiple linear regression, IOP is adjusted for the following covariates: age, sex, and the top 10 genetic PCs separately for WB and BB ancestries. We then implement the inverse rank normal transformation to the adjusted IOP to induce normality. We employ a similar QC procedure outlined above for the Geuvadis genotype data. We filter out SNPs based on MAF *<* 0.05, HWE test p-value *<* 5*×*10^−8^, and retain a final set of 241977 SNPs that overlapped with the QCed SNPs in the Geuvadis data. We restrict our analysis to the genes on the autosomal chromosomes only. We standardize the complex trait, gene expression, and the columns of genotype matrices both in transcriptome and GWAS data. For the gene under consideration, we include local SNPs within 500 kilobases of the gene’s upstream and downstream regions beginning from the transcription start and end sites. We integrate the EUR Geuvadis data with the WB UKB data to conduct the single-ancestry TWAS, whereas we combine the EUR and AFR Geuvadis data with WB and BB UKB data for the multi-ancestry TWAS.

We compare glossTWAS with PMR-Egger and PrediXcan in single-ancestry TWAS of IOP. We exclude the approach TIGAR because it demonstrated comparatively suboptimal performance in our simulation studies, and implementing it on the UKB cloud computing platform is also computationally challenging. For the multi-ancestry TWAS, we evaluate the performance of glossTWAS in comparison with METRO. To adjust for multiple testing across 12517 protein-coding genes, we apply the Benjamini-Hochberg procedure to control the false discovery rate (FDR) for frequentist approaches, whereas for glossTWAS, we control the Bayesian FDR (BFDR). To limit the number of false discoveries, we set the thresholds for both FDR and BFDR at 0.01. As the BFDR depends on the prior probability that the null hypothesis for a given gene is true, we need to choose this probability in a sensible way. Let, *π*_0_ = *P* (*H*_0*i*_ is true), *i* = 1, 2, · · ·, 12517. Implementing TWAS, several genetic studies have discovered numerous genes associated with a wide range of complex traits. Majumdar et al. reported 18.6% polygenicity for height and 6.9% polygencity for body mass index, where polygenicity is defined as the proportion of non-null genes affecting the trait [41]. Li et al. identified 4.8% of protein-coding genes have non-zero effects on the total cholesterol [18]. Guided by these findings, we explore two different choices of *π*_0_, 0.9 and 0.95, in our TWAS analysis of IOP, which incorporates prior knowledge that 10% and 5% of the genes are expected to be associated with IOP, respectively.

## Notes

### Competing Interest Statement

The authors have declared no competing interest.

### Summary of Updates

Some revisions have been carried out thought the main text and supplementary materials to improve the earlier version of the manuscript. Author information has been updated too.

## References

[1] J. N. Hirschhorn and M. J. Daly, “Genome-wide association studies for common diseases and complex traits,” Nature reviews genetics, vol. 6, no. 2, pp. 95–108, 2005.

[2] D. L. Nicolae, E. Gamazon, W. Zhang, S. Duan, M. E. Dolan, and N. J. Cox, “Trait-associated snps are more likely to be eqtls: annotation to enhance discovery from gwas,” PLoS genetics, vol. 6, no. 4, p. e1000888, 2010.

[3] E. R. Gamazon, H. E. Wheeler, K. P. Shah, S. V. Mozaffari, K. Aquino-Michaels, R. J. Carroll, A. E. Eyler, J. C. Denny, G. Consortium, D. L. Nicolae et al., “A gene-based association method for mapping traits using reference transcriptome data,” Nature genetics, vol. 47, no. 9, pp. 1091–1098, 2015.

[4] C. Cao, B. Ding, Q. Li, D. Kwok, J. Wu, and Q. Long, “Power analysis of transcriptome-wide association study: Implications for practical protocol choice,” PLoS genetics, vol. 17, no. 2, p. e1009405, 2021.

[5] A. Gusev, A. Ko, H. Shi, G. Bhatia, W. Chung, B. W. Penninx, R. Jansen, E. J. De Geus, D. I. Boomsma, F. A. Wright et al., “Integrative approaches for large-scale transcriptome-wide association studies,” Nature genetics, vol. 48, no. 3, pp. 245–252, 2016.

[6] J. Mai, M. Lu, Q. Gao, J. Zeng, and J. Xiao, “Transcriptome-wide association studies: recent advances in methods, applications and available databases,” Communications Biology, vol. 6, no. 1, p. 899, 2023.

[7] R. Tibshirani, “Regression shrinkage and selection via the lasso,” Journal of the Royal Statistical Society Series B: Statistical Methodology, vol. 58, no. 1, pp. 267–288, 1996.

[8] C. de Leeuw, J. Werme, J. E. Savage, W. J. Peyrot, and D. Posthuma, “On the interpretation of transcriptome-wide association studies,” PLoS Genetics, vol. 19, no. 9, p. e1010921, 2023.

[9] H. Zou and T. Hastie, “Regularization and variable selection via the elastic net,” Journal of the Royal Statistical Society Series B: Statistical Methodology, vol. 67, no. 2, pp. 301–320, 2005.

[10] J. M. Luningham, J. Chen, S. Tang, P. L. De Jager, D. A. Bennett, A. S. Buchman, and J. Yang, “Bayesian genome-wide twas method to leverage both cis-and trans-eqtl information through summary statistics,” The American Journal of Human Genetics, vol. 107, no. 4, pp. 714–726, 2020.

[11] S. Nagpal, X. Meng, M. P. Epstein, L. C. Tsoi, M. Patrick, G. Gibson, P. L. De Jager, D. A. Bennett, A. P. Wingo, T. S. Wingo et al., “Tigar: an improved bayesian tool for transcriptomic data imputation enhances gene mapping of complex traits,” The American Journal of Human Genetics, vol. 105, no. 2, pp. 258–266, 2019.

[12] C. M. Carvalho, N. G. Polson, and J. G. Scott, “The horseshoe estimator for sparse signals,” Biometrika, vol. 97, no. 2, pp. 465–480, 2010.

[13] H. Ishwaran and J. S. Rao, “Spike and slab variable selection: Frequentist and bayesian strategies,” The Annals of Statistics, vol. 33, no. 2, pp. 730–773, 2005.

[14] Z. Yuan, H. Zhu, P. Zeng, S. Yang, S. Sun, C. Yang, J. Liu, and X. Zhou, “Testing and controlling for horizontal pleiotropy with probabilistic mendelian randomization in transcriptome-wide association studies,” Nature communications, vol. 11, no. 1, p. 3861, 2020.

[15] L. S. Mogil, A. Andaleon, A. Badalamenti, S. P. Dickinson, X. Guo, J. I. Rotter, W. C. Johnson, H. K. Im, Y. Liu, and H. E. Wheeler, “Genetic architecture of gene expression traits across diverse populations,” PLoS genetics, vol. 14, no. 8, p. e1007586, 2018.

[16] A. Bhadra, J. Datta, N. G. Polson, and B. Willard, “Lasso meets horseshoe,” Statistical Science, vol. 34, no. 3, pp. 405–427, 2019.

[17] Z. Lu, S. Gopalan, D. Yuan, D. V. Conti, B. Pasaniuc, A. Gusev, and N. Mancuso, “Multi-ancestry fine-mapping improves precision to identify causal genes in transcriptome-wide association studies,” The American Journal of Human Genetics, vol. 109, no. 8, pp. 1388–1404, 2022.

[18] Z. Li, W. Zhao, L. Shang, T. H. Mosley, S. L. Kardia, J. A. Smith, and X. Zhou, “Metro: Multi-ancestry transcriptome-wide association studies for powerful gene-trait association detection,” The American Journal of Human Genetics, vol. 109, no. 5, pp. 783–801, 2022.

[19] Y. Benjamini and Y. Hochberg, “Controlling the false discovery rate: a practical and powerful approach to multiple testing,” Journal of the Royal statistical society: series B (Methodological), vol. 57, no. 1, pp. 289–300, 1995.

[20] K. P. Burdon, S. Macgregor, A. W. Hewitt, S. Sharma, G. Chidlow, R. A. Mills, P. Danoy, R. Casson, A. C. Viswanathan, J. Z. Liu et al., “Genome-wide association study identifies susceptibility loci for open angle glaucoma at tmco1 and cdkn2b-as1,” Nature genetics, vol. 43, no. 6, pp. 574–578, 2011.

[21] X. R. Gao, H. Huang, D. R. Nannini, F. Fan, and H. Kim, “Genome-wide association analyses identify new loci influencing intraocular pressure,” Human molecular genetics, vol. 27, no. 12, pp. 2205–2213, 2018.

[22] H. Springelkamp, A. I. Iglesias, G. Cuellar-Partida, N. Amin, K. P. Burdon, E. M. van Leeuwen, P. Gharahkhani, A. Mishra, S. J. van der Lee, A. W. Hewitt et al., “Arhgef12 influences the risk of glaucoma by increasing intraocular pressure,” Human molecular genetics, vol. 24, no. 9, pp. 2689–2699, 2015.

[23] H. Choquet, K. K. Thai, J. Yin, T. J. Hoffmann, M. N. Kvale, Y. Banda, C. Schaefer, N. Risch, K. S. Nair, R. Melles et al., “A large multi-ethnic genome-wide association study identifies novel genetic loci for intraocular pressure,” Nature communications, vol. 8, no. 1, p. 2108, 2017.

[24] A. P. Khawaja, J. N. Cooke Bailey, N. J. Wareham, R. A. Scott, M. Simcoe, R. P. Igo Jr, Y. E. Song, R. Wojciechowski, C.-Y. Cheng, P. T. Khaw et al., “Genome-wide analyses identify 68 new loci associated with intraocular pressure and improve risk prediction for primary open-angle glaucoma,” Nature genetics, vol. 50, no. 6, pp. 778–782, 2018.

[25] L. Kolberg, U. Raudvere, I. Kuzmin, J. Vilo, and H. Peterson, “gprofiler2–an r package for gene list functional enrichment analysis and namespace conversion toolset g: Profiler,” F1000Research, vol. 9, pp. ELIXIR–709, 2020.

[26] G. O. Consortium, “The gene ontology (go) database and informatics resource,” Nucleic acids research, vol. 32, no. Suppl 1, pp. D258–D261, 2004.

[27] M. Kanehisa, M. Furumichi, Y. Sato, Y. Matsuura, and M. Ishiguro-Watanabe, “Kegg: biological systems database as a model of the real world,” Nucleic acids research, vol. 53, no. D1, pp. D672–D677, 2025.

[28] M. Milacic, D. Beavers, P. Conley, C. Gong, M. Gillespie, J. Griss, R. Haw, B. Jassal, L. Matthews, B. May et al., “The reactome pathway knowledgebase 2024,” Nucleic acids research, vol. 52, no. D1, pp. D672–D678, 2024.

[29] C. J. Mungall, J. A. McMurry, S. Köhler, J. P. Balhoff, C. Borromeo, M. Brush, S. Carbon, T. Conlin, N. Dunn, M. Engelstad et al., “The monarch initiative: an integrative data and analytic platform connecting phenotypes to genotypes across species,” Nucleic acids research, vol. 45, no. D1, pp. D712–D722, 2017.

[30] D. Szklarczyk, K. Nastou, M. Koutrouli, R. Kirsch, F. Mehryary, R. Hachilif, D. Hu, M. E. Peluso, Q. Huang, T. Fang et al., “The string database in 2025: protein networks with directionality of regulation,” Nucleic Acids Research, vol. 53, no. D1, pp. D730– D737, 2025.

[31] K. E. Keller and D. M. Peters, “Pathogenesis of glaucoma: Extracellular matrix dysfunction in the trabecular meshwork-a review,” Clinical & experimental ophthalmology, vol. 50, no. 2, pp. 163–182, 2022.

[32] T. Wang, H. R. Kimmel, C. Park, H. Ryoo, J. Liu, G. H. Underhill, and P. P. Pattabiraman, “Regulatory role of cholesterol in modulating actin dynamics and cell adhesive interactions in the trabecular meshwork,” BioRxiv, 2024.

[33] H. C. Webber, J. Y. Bermudez, J. C. Millar, W. Mao, and A. F. Clark, “The role of wnt/β-catenin signaling and k-cadherin in the regulation of intraocular pressure,” Investigative ophthalmology & visual science, vol. 59, no. 3, pp. 1454–1466, 2018.

[34] C. Knox, M. Wilson, C. M. Klinger, M. Franklin, E. Oler, A. Wilson, A. Pon, J. Cox, N. E. Chin, S. A. Strawbridge et al., “Drugbank 6.0: the drugbank knowledgebase for 2024,” Nucleic acids research, vol. 52, no. D1, pp. D1265–D1275, 2024.

[35] P. E. Tenconi, M. S. Echevarría, N. M. Giusto, and M. V. Mateos, “The phospholipase d pathway modulates oxidative stress in retinal pigment epithelium cells exposed to high glucose levels,” Investigative Ophthalmology & Visual Science, vol. 64, no. 8, pp. 4475–4475, 2023.

[36] G. L. Wojcik, M. Graff, K. K. Nishimura, R. Tao, J. Haessler, C. R. Gignoux, H. M. Highland, Y. M. Patel, E. P. Sorokin, C. L. Avery et al., “Genetic analyses of diverse populations improves discovery for complex traits,” Nature, vol. 570, no. 7762, pp. 514–518, 2019.

[37] S. F. Grant, G. Thorleifsson, I. Reynisdottir, R. Benediktsson, A. Manolescu, J. Sainz, A. Helgason, H. Stefansson, V. Emilsson, A. Helgadottir et al., “Variant of transcription factor 7-like 2 (tcf7l2) gene confers risk of type 2 diabetes,” Nature genetics, vol. 38, no. 3, pp. 320–323, 2006.

[38] K. A. Fawcett and I. Barroso, “The genetics of obesity: Fto leads the way,” Trends in genetics, vol. 26, no. 6, pp. 266–274, 2010.

[39] G. Casella and C. P. Robert, “Rao-blackwellisation of sampling schemes,” Biometrika, vol. 83, no. 1, pp. 81–94, 1996.

[40] J. Yang, L. G. Fritsche, X. Zhou, and G. Abecasis, “A scalable bayesian method for integrating functional information in genome-wide association studies,” The American Journal of Human Genetics, vol. 101, no. 3, pp. 404–416, 2017.

[41] A. Majumdar and B. Pasaniuc, “A bayesian method for estimating gene-level polygenicity under the framework of transcriptome-wide association study,” Statistics in Medicine, vol. 42, no. 26, pp. 4867–4885, 2023.

[42] Y. Liu, S. Chen, Z. Li, A. C. Morrison, E. Boerwinkle, and X. Lin, “Acat: a fast and powerful p value combination method for rare-variant analysis in sequencing studies,” The American Journal of Human Genetics, vol. 104, no. 3, pp. 410–421, 2019.

[43] Z. Su, J. Marchini, and P. Donnelly, “Hapgen2: simulation of multiple disease snps,” Bioinformatics, vol. 27, no. 16, pp. 2304–2305, 2011.

[44] M. A. Coram, S. I. Candille, Q. Duan, K. H. K. Chan, Y. Li, C. Kooperberg, A. P. Reiner, and H. Tang, “Leveraging multi-ethnic evidence for mapping complex traits in minority populations: an empirical bayes approach,” The American Journal of Human Genetics, vol. 96, no. 5, pp. 740–752, 2015.

[45] A. S. Whittemore, “A bayesian false discovery rate for multiple testing,” Journal of Applied Statistics, vol. 34, no. 1, pp. 1–9, 2007.

[46] R. E. Kass and A. E. Raftery, “Bayes factors,” Journal of the american statistical association, vol. 90, no. 430, pp. 773–795, 1995.

[47] T. Lappalainen, M. Sammeth, M. R. Friedländer, P.A. ‘t Hoen, J. Monlong, M. A. Rivas, M. Gonzalez-Porta, N. Kurbatova, T. Griebel, P. G. Ferreira et al., “Transcriptome and genome sequencing uncovers functional variation in humans,” Nature, vol. 501, no. 7468, pp. 506–511, 2013.

[48] O. Troyanskaya, M. Cantor, G. Sherlock, P. Brown, T. Hastie, R. Tibshirani, D. Botstein, and R. B. Altman, “Missing value estimation methods for dna microarrays,” Bioinformatics, vol. 17, no. 6, pp. 520–525, 2001.

[49] T. M. Beasley, S. Erickson, and D. B. Allison, “Rank-based inverse normal transformations are increasingly used, but are they merited?” Behavior genetics, vol. 39, pp. 580–595, 2009.

