## Supplementary material for "A unified hierarchical Bayesian approach to transcriptome-wide association study": Link for Supplementary Material

### 1 Derivation of posterior odds for glossTWAS in multi-ancestry TWAS setup

In the multi-ancestry TWAS framework, we regard the following hypothesis $H_0 : \beta = 0$ versus $H_1 : \beta \neq 0$. Let $D$ denote the complete dataset, comprising the gene expression and outcome from the transcriptome and GWAS data for the two ancestries, respectively. The posterior odds of $H_1$ versus $H_0$ is $\frac{P[H_1|D]}{P[H_0|D]}$ and the prior odds is $\frac{P[H_1]}{P[H_0]}$. The prior odds of $H_1$ versus $H_0$ is $\frac{b_2}{b_1}$. For the computation of posterior odds, we need to evaluate the posterior probability of null association, i.e., $P[H_0|D]$. Using the Rao-Blackwellization technique, an approximate expression for $P[H_0|D]$ is derived as follows:

$$
\begin{aligned}
& P\big[H_0|D\big] \\
&= P\big[\beta = 0|D\big] \\
&= \int\limits_{\sigma_\epsilon^2}\int\limits_{\sigma_d^2}\int\limits_{q}\int\limits_{w}\int\limits_{\boldsymbol{\alpha}_1}\int\limits_{\boldsymbol{\alpha}_2} P\Big[\beta = 0, \boldsymbol{\alpha}_1, \boldsymbol{\alpha}_2, w, q, \sigma_d^2, \sigma_\epsilon^2 \Big| D\Big]\ d\boldsymbol{\alpha}_2\ d\boldsymbol{\alpha}_1\ dw\ dq\ d\sigma_d^2\ d\sigma_\epsilon^2 \\
&= \int\limits_{\sigma_\epsilon^2}\int\limits_{\sigma_d^2}\int\limits_{q}\int\limits_{w}\int\limits_{\boldsymbol{\alpha}_1}\int\limits_{\boldsymbol{\alpha}_2} P\Big[\beta = 0 \Big| \boldsymbol{\alpha}_1, \boldsymbol{\alpha}_2, w, q, \sigma_d^2, \sigma_\epsilon^2, D\Big] f(\boldsymbol{\alpha}_1, \boldsymbol{\alpha}_2, w, q, \sigma_d^2, \sigma_\epsilon^2|D) d\boldsymbol{\alpha}_2\ d\boldsymbol{\alpha}_1\ dw\ dq\ d\sigma_d^2\ d\sigma_\epsilon^2 \\
&= E_{\boldsymbol{\alpha}_1, \boldsymbol{\alpha}_2, w, q, \sigma_d^2, \sigma_\epsilon^2|D}\Big[P\Big[\beta = 0 \Big| \boldsymbol{\alpha}_1, \boldsymbol{\alpha}_2, w, q, \sigma_d^2, \sigma_\epsilon^2, D\Big]\Big] \\
&\approx \frac{1}{M}\sum_{t=1}^{M} P\left[\beta = 0 \Big| \boldsymbol{\alpha}_1^{(t)}, \boldsymbol{\alpha}_2^{(t)}, w^{(t)}, q^{(t)}, \sigma_d^{2(t)}, \sigma_\epsilon^{2(t)}, D\right];
\end{aligned} \tag{1}
$$

where $(\boldsymbol{\alpha}_1^{(t)}, \boldsymbol{\alpha}_2^{(t)}, w^{(t)}, q^{(t)}, \sigma_d^{2(t)}, \sigma_\epsilon^{2(t)})$ is the $t^{\text{th}}$ posterior draw from $M$ MCMC iterations. The term $P\left[\beta = 0 \Big| \boldsymbol{\alpha}_1^{(t)}, \boldsymbol{\alpha}_2^{(t)}, w^{(t)}, q^{(t)}, \sigma_d^{2(t)}, \sigma_\epsilon^{2(t)}, D\right]$ represents the posterior probability that $\beta = 0$ in the corresponding full conditional distribution, evaluated at the $t^{\text{th}}$ step of MCMC. The explicit form of this quantity is provided in the Equation 11.

### 2 Derivation of the full conditional distributions for glossTWAS in single-ancestry TWAS setup

The hierarchical structure of glossTWAS in the single-ancestry setup is given by:

$$
\begin{aligned}
\mathbf{x}_{\text{ref}}|\boldsymbol{\alpha}, \sigma_e^2 &\sim \text{N}_n(G_{\text{ref}}\boldsymbol{\alpha}, \sigma_e^2 \text{I}_n) \\
\mathbf{y}|\beta, \boldsymbol{\alpha}, \sigma_\epsilon^2 &\sim \text{N}_N(\beta G\boldsymbol{\alpha}, \sigma_\epsilon^2 \text{I}_N) \\
\alpha_j|\lambda_j, \tau, \sigma_e^2 &\overset{ind}{\sim} \text{N}(0, \sigma_e^2\tau^2\lambda_j^2), \quad j = 1(1)m \\
\lambda_j &\overset{ind}{\sim} \text{C}^+(0,1) \\
\tau &\sim \text{C}^+(0,1) \\
\beta|q, \sigma_d^2, \sigma_\epsilon^2 &\sim (1-q)\delta_{\{0\}}(\beta) + q\text{N}(0, \sigma_d^2\sigma_\epsilon^2) \\
q &\sim \text{Beta}(b_1, b_2) \\
\sigma_d^2 &\sim \text{IG}(d_1, d_2) \\
\sigma_e^2 &\sim \text{IG}(e_1, e_2) \\
\sigma_\epsilon^2 &\sim \text{IG}(\epsilon_1, \epsilon_2)
\end{aligned}
\tag{2}
$$

Let us denote $\boldsymbol{\alpha} = (\alpha_1, \alpha_2, \cdots, \alpha_m)^{\text{T}}$, where $\alpha_j$ is the effect of $j^{\text{th}}$ cis-SNP on the gene expression for $j = 1(1)m$. We specify the SNP-specific shrinkage parameters in the form of a vector as $\boldsymbol{\lambda} = (\lambda_1, \lambda_2, \cdots, \lambda_m)^{\text{T}}$. Let us define $D_\lambda = diag(\lambda_1^2, \lambda_2^2, \cdots, \lambda_m^2)$, where $D_\lambda$ is a $m \times m$ diagonal matrix. Conditional on $\boldsymbol{\lambda}, \tau$ and $\sigma_e^2$, the prior distribution of $\boldsymbol{\alpha}$ is given by

$$\boldsymbol{\alpha}|\boldsymbol{\lambda}, \tau, \sigma_e^2 \sim \text{N}_m(\mathbf{0}, \sigma_e^2\tau^2 D_\lambda)$$

We now proceed to derive the full conditional distributions of the model parameters.

#### 2.1 Full conditional distribution of $\boldsymbol{\alpha}$

We use the notation $\boldsymbol{\alpha}|\cdot$ to denote the full conditional posterior distribution of $\boldsymbol{\alpha}$, with corresponding density $f(\boldsymbol{\alpha}|\cdot)$. Analogous notation is adopted for other model parameters.

The full conditional distribution of $\boldsymbol{\alpha}$ is given by

$$
\begin{aligned}
f(\boldsymbol{\alpha}|\cdot) &\propto f(\mathbf{x}_{\text{ref}}|\boldsymbol{\alpha},\sigma_e^2)f(\mathbf{y}|\beta,\boldsymbol{\alpha},\sigma_\epsilon^2)f(\boldsymbol{\alpha}|\boldsymbol{\lambda},\tau,\sigma_e^2)\\
&\propto \exp\left[-\frac{1}{2\sigma_e^2}(\mathbf{x}_{\text{ref}}-G_{\text{ref}}\boldsymbol{\alpha})^{\text{T}}(\mathbf{x}_{\text{ref}}-G_{\text{ref}}\boldsymbol{\alpha})\right]\exp\left[-\frac{1}{2\sigma_\epsilon^2}(\mathbf{y}-\beta G\boldsymbol{\alpha})^{\text{T}}(\mathbf{y}-\beta G\boldsymbol{\alpha})\right]\\
&\quad \exp\left[-\frac{\boldsymbol{\alpha}^{\text{T}}D_\lambda^{-1}\boldsymbol{\alpha}}{2\sigma_e^2\tau^2}\right]\\
&\propto \exp\left\{-\frac{1}{2}\left[\boldsymbol{\alpha}^{\text{T}}\left(\frac{1}{\sigma_e^2}G_{\text{ref}}^{\text{T}}G_{\text{ref}}+\frac{\beta^2}{\sigma_\epsilon^2}G^{\text{T}}G+\frac{1}{\sigma_e^2\tau^2}D_\lambda^{-1}\right)\boldsymbol{\alpha}-2\boldsymbol{\alpha}^{\text{T}}\left(\frac{1}{\sigma_e^2}G_{\text{ref}}^{\text{T}}\mathbf{x}_{\text{ref}}+\frac{\beta}{\sigma_\epsilon^2}G^{\text{T}}\mathbf{y}\right)\right]\right\}\\
&\propto \exp\left\{-\frac{1}{2}\left[\boldsymbol{\alpha}^{\text{T}}\Sigma\boldsymbol{\alpha}-2\boldsymbol{\alpha}^{\text{T}}\left(\frac{1}{\sigma_e^2}G_{\text{ref}}^{\text{T}}\mathbf{x}_{\text{ref}}+\frac{\beta}{\sigma_\epsilon^2}G^{\text{T}}\mathbf{y}\right)\right]\right\};\\
&\text{where } \Sigma=\frac{1}{\sigma_e^2}G_{\text{ref}}^{\text{T}}G_{\text{ref}}+\frac{\beta^2}{\sigma_\epsilon^2}G^{\text{T}}G+\frac{1}{\sigma_e^2\tau^2}D_\lambda^{-1}
\end{aligned}
$$

Hence, the full conditional distribution of $\boldsymbol{\alpha}$ is given by

$$
\boldsymbol{\alpha}|\cdot \sim \text{N}_m\left(\Sigma^{-1}\left(\frac{1}{\sigma_e^2}G_{\text{ref}}^{\text{T}}\mathbf{x}_{\text{ref}}+\frac{\beta}{\sigma_\epsilon^2}G^{\text{T}}\mathbf{y}\right),\Sigma^{-1}\right)
$$

#### 2.2 Full conditional distribution of $\boldsymbol{\lambda}$

As for $j = 1(1)m$, $\lambda_j$s are independently distributed, we update $\boldsymbol{\lambda}$ within the MCMC framework by sequentially updating its individual components. Specifically, for each $j = 1(1)m$, the full conditional distribution of $\lambda_j$ is given by

$$
\begin{aligned}
f(\lambda_j|\cdot) &\propto f(\alpha_j|\lambda_j,\tau,\sigma_e^2)f(\lambda_j)\\
&\propto \left\{\frac{1}{\sigma_e\lambda_j\tau}\ \exp\left[-\frac{\alpha_j^2}{2\sigma_e^2\tau^2\lambda_j^2}\right]\right\}\left\{\frac{1}{1+\lambda_j^2}\right\}\\
&\propto \frac{1}{\lambda_j(1+\lambda_j^2)}\ \exp\left[-\frac{\alpha_j^2}{2\sigma_e^2\tau^2\lambda_j^2}\right]
\end{aligned}
\tag{3}
$$

The resulting full conditional posterior distribution of $\lambda_j$ does not correspond to a standard distribution from which direct sampling is feasible. Consequently, we update $\lambda_j$ using a Metropolis-Hastings step within the MCMC algorithm, as described in section 3.1.

#### 2.3 Full conditional distribution of $\tau$

The full conditional distribution of $\tau$ is given by

$$\begin{aligned} f(\tau|\cdot) &\propto f(\boldsymbol{\alpha}|\boldsymbol{\lambda}, \tau, \sigma_e^2) f(\tau) \\ &\propto \left\{ \frac{1}{\tau^m} \exp\left[ -\frac{\boldsymbol{\alpha}^{\mathrm{T}} D_\lambda^{-1} \boldsymbol{\alpha}}{2\sigma_e^2 \tau^2} \right] \right\} \left\{ \frac{1}{1+\tau^2} \right\} \\ &\propto \frac{1}{\tau^m (1+\tau^2)} \exp\left[ -\frac{\boldsymbol{\alpha}^{\mathrm{T}} D_\lambda^{-1} \boldsymbol{\alpha}}{2\sigma_e^2 \tau^2} \right] \end{aligned} \tag{4}$$

Since the posterior distribution of $\tau$ is not conditionally conjugate, we draw samples from $f(\tau|\cdot)$ using the Metropolis-Hastings algorithm. The steps of the algorithm are provided in section 3.2.

#### 2.4 Full conditional distribution of $\sigma_e^2$

The full conditional distribution of $\sigma_e^2$ is given by

$$\begin{aligned} f(\sigma_e^2|\cdot) &\propto f(\mathbf{x}_{\mathrm{ref}}|\boldsymbol{\alpha}, \sigma_e^2) f(\boldsymbol{\alpha}|\boldsymbol{\lambda}, \tau, \sigma_e^2) f(\sigma_e^2) \\ &\propto \left\{ (\sigma_e^2)^{-\frac{n}{2}} \exp\left[ -\frac{1}{2\sigma_e^2} (\mathbf{x}_{\mathrm{ref}} - G_{\mathrm{ref}}\boldsymbol{\alpha})^{\mathrm{T}} (\mathbf{x}_{\mathrm{ref}} - G_{\mathrm{ref}}\boldsymbol{\alpha}) \right] \right\} \left\{ (\sigma_e^2)^{-\frac{m}{2}} \exp\left[ -\frac{\boldsymbol{\alpha}^{\mathrm{T}} D_\lambda^{-1} \boldsymbol{\alpha}}{2\sigma_e^2 \tau^2} \right] \right\} \\ &\quad \left\{ (\sigma_e^2)^{-(e_1+1)} \exp\left[ -\frac{e_2}{\sigma_e^2} \right] \right\} \\ &\propto \left(\sigma_e^2\right)^{-\left(e_1 + \frac{n}{2} + \frac{m}{2} + 1\right)} \exp\left[ -\frac{1}{\sigma_e^2} \left( e_2 + \frac{\boldsymbol{\alpha}^{\mathrm{T}} D_\lambda^{-1} \boldsymbol{\alpha}}{2\tau^2} + \frac{1}{2} (\mathbf{x}_{\mathrm{ref}} - G_{\mathrm{ref}}\boldsymbol{\alpha})^{\mathrm{T}} (\mathbf{x}_{\mathrm{ref}} - G_{\mathrm{ref}}\boldsymbol{\alpha}) \right) \right] \end{aligned}$$

Therefore, the full conditional distribution of $\sigma_e^2$ is given by

$$\sigma_e^2|\cdot \sim \mathrm{IG}\left( e_1 + \frac{n}{2} + \frac{m}{2},\ e_2 + \frac{\boldsymbol{\alpha}^{\mathrm{T}} D_\lambda^{-1} \boldsymbol{\alpha}}{2\tau^2} + \frac{1}{2} (\mathbf{x}_{\mathrm{ref}} - G_{\mathrm{ref}}\boldsymbol{\alpha})^{\mathrm{T}} (\mathbf{x}_{\mathrm{ref}} - G_{\mathrm{ref}}\boldsymbol{\alpha}) \right)$$

#### 2.5 Full conditional distribution of $\beta$

The full conditional distribution of $\beta$ is derived as follows:

$$
\begin{aligned}
f(\beta|\cdot) &\propto f(\mathbf{y}|\boldsymbol{\alpha},\beta,\sigma_\epsilon^2)f(\beta|q,\sigma_d^2,\sigma_\epsilon^2)\\
&\propto \exp\left[-\frac{1}{2\sigma_\epsilon^2}(\mathbf{y}-\beta G\boldsymbol{\alpha})^{\mathrm{T}}(\mathbf{y}-\beta G\boldsymbol{\alpha})\right]\left\{(1-q)\delta_{\{0\}}(\beta)+\frac{q}{\sqrt{2\pi}\sigma_d\sigma_\epsilon}\ \exp\left[-\frac{\beta^2}{2\sigma_d^2\sigma_\epsilon^2}\right]\right\}\\
&\propto \exp\left[-\frac{1}{2\sigma_\epsilon^2}\left(\beta^2\boldsymbol{\alpha}^{\mathrm{T}}G^{\mathrm{T}}G\boldsymbol{\alpha}-2\beta\boldsymbol{\alpha}^{\mathrm{T}}G^{\mathrm{T}}\mathbf{y}\right)\right]\left\{(1-q)\delta_{\{0\}}(\beta)+\frac{q}{\sqrt{2\pi}\sigma_d\sigma_\epsilon}\ \exp\left[-\frac{\beta^2}{2\sigma_d^2\sigma_\epsilon^2}\right]\right\}\\
&\propto (1-q)\delta_{\{0\}}(\beta)+\frac{q}{\sqrt{2\pi}\sigma_d\sigma_\epsilon}\ \exp\left[-\frac{\boldsymbol{\alpha}^{\mathrm{T}}G^{\mathrm{T}}G\boldsymbol{\alpha}+\frac{1}{\sigma_d^2}}{2\sigma_\epsilon^2}\left(\beta^2-2\beta\ \frac{\boldsymbol{\alpha}^{\mathrm{T}}G^{\mathrm{T}}\mathbf{y}}{\boldsymbol{\alpha}^{\mathrm{T}}G^{\mathrm{T}}G\boldsymbol{\alpha}+\frac{1}{\sigma_d^2}}\right)\right]\\
&\propto (1-q)\delta_{\{0\}}(\beta)+\frac{q}{\sqrt{2\pi}\sigma_d\sigma_\epsilon}\ \exp\left[-\frac{1}{2\widetilde{\sigma}^2}(\beta^2-2\beta\widetilde{\mu})\right]\\
&\propto (1-q)\delta_{\{0\}}(\beta)+\left(\frac{q\widetilde{\sigma}}{\sigma_d\sigma_\epsilon}\exp\left[\frac{\widetilde{\mu}^2}{2\widetilde{\sigma}^2}\right]\right)\left\{\frac{1}{\sqrt{2\pi}\widetilde{\sigma}}\exp\left[-\frac{1}{2\widetilde{\sigma}^2}(\beta-\widetilde{\mu})^2\right]\right\};\\
&\text{where}\quad \widetilde{\mu}=\frac{\boldsymbol{\alpha}^{\mathrm{T}}G^{\mathrm{T}}\mathbf{y}}{\boldsymbol{\alpha}^{\mathrm{T}}G^{\mathrm{T}}G\boldsymbol{\alpha}+\frac{1}{\sigma_d^2}}\quad\text{and}\quad \widetilde{\sigma}^2=\frac{\sigma_\epsilon^2}{\boldsymbol{\alpha}^{\mathrm{T}}G^{\mathrm{T}}G\boldsymbol{\alpha}+\frac{1}{\sigma_d^2}}
\end{aligned}
$$

Therefore, the full conditional distribution of $\beta$ can be expressed as a mixture of point mass function $\delta_{\{0\}}(\beta)$ and normal distribution $\mathrm{N}(\widetilde{\mu},\widetilde{\sigma}^2)$. The mixture weight corresponding to $\delta_{\{0\}}(\beta)$ is given by

$$
\widetilde{w}=\frac{1-q}{1-q+\frac{q\widetilde{\sigma}}{\sigma_d\sigma_\epsilon}\exp\left(\frac{\widetilde{\mu}^2}{2\widetilde{\sigma}^2}\right)} \tag{5}
$$

Thus, the conditional posterior distribution of $\beta$ is given by

$$
\beta|\cdot\sim\begin{cases}\delta_{\{0\}}(\beta), & \text{with probability}\quad \widetilde{w}=\frac{1-q}{1-q+\frac{q\widetilde{\sigma}}{\sigma_d\sigma_\epsilon}\exp\left(\frac{\widetilde{\mu}^2}{2\widetilde{\sigma}^2}\right)}\\[2ex] \mathrm{N}(\widetilde{\mu},\widetilde{\sigma}^2), & \text{with probability}\quad 1-\widetilde{w}\end{cases}
$$

#### 2.6 Full conditional distribution of $q$

We now derive the full conditional distribution of $q$. Let $\phi(x;\mu,\sigma^2)$ denote the density of the normal distribution evaluated at the point $x$ with mean $\mu$ and variance $\sigma^2$. The beta function is defined as $B(s,t)=\frac{\Gamma(s)\Gamma(t)}{\Gamma(s+t)}$ for $s,t>0$, where $\Gamma(\cdot)$ denotes the gamma function.

The full conditional distribution of $q$ is given by

$$
\begin{aligned}
f(q|\cdot) &\propto f(\beta|q,\sigma_d^2,\sigma_\epsilon^2)f(q) \\
&\propto \left\{(1-q)\delta_{\{0\}}(\beta) + q\ \phi(\beta;0,\sigma_d^2\sigma_\epsilon^2)\right\}\left\{q^{b_1-1}(1-q)^{b_2-1}\right\} \\
&\propto \left\{\delta_{\{0\}}(\beta)q^{b_1-1}(1-q)^{b_2} + \phi(\beta;0,\sigma_d^2\sigma_\epsilon^2)q^{b_1}(1-q)^{b_2-1}\right\} \\
&\propto \left\{\delta_{\{0\}}(\beta)B(b_1,b_2+1)\text{Beta}(b_1,b_2+1) + \phi(\beta;0,\sigma_d^2\sigma_\epsilon^2)B(b_1+1,b_2)\text{Beta}(b_1+1,b_2)\right\}
\end{aligned}
$$

Therefore, the full conditional distribution of q is a two-component mixture of Beta distributions, given by

$$
q|\cdot \sim \begin{cases} \text{Beta}(b_1,b_2+1), & \text{with probability} \quad \widetilde{w} = \frac{\delta_{\{0\}}(\beta)B(b_1,b_2+1)}{\delta_{\{0\}}(\beta)B(b_1,b_2+1)+\phi(\beta;0,\sigma_d^2\sigma_\epsilon^2)B(b_1+1,b_2)} \\ \text{Beta}(b_1+1,b_2), & \text{with probability} \quad 1-\widetilde{w} \end{cases} \tag{6}
$$

#### 2.7 Full conditional distribution of $\sigma_d^2$

The full conditional distribution of $\sigma_d^2$ is given by

$$
\begin{aligned}
f(\sigma_d^2|\cdot) &\propto f(\beta|q,\sigma_d^2,\sigma_\epsilon^2)f(\sigma_d^2) \\
&\propto \left\{(1-q)\delta_{\{0\}}(\beta) + \frac{q}{\sqrt{2\pi}\sigma_d\sigma_\epsilon}\ \exp\left[-\frac{\beta^2}{2\sigma_d^2\sigma_\epsilon^2}\right]\right\}\left\{(\sigma_d^2)^{-(d_1+1)}\exp\left[-\frac{d_2}{\sigma_d^2}\right]\right\} \\
&\propto \left\{(1-q)\delta_{\{0\}}(\beta)(\sigma_d^2)^{-(d_1+1)}\exp\left[-\frac{d_2}{\sigma_d^2}\right] + \frac{q}{\sqrt{2\pi}\sigma_\epsilon}(\sigma_d^2)^{-(d_1+\frac{1}{2}+1)}\exp\left[-\frac{1}{\sigma_d^2}\left(d_2+\frac{\beta^2}{2\sigma_\epsilon^2}\right)\right]\right\} \\
&\propto \frac{(1-q)\delta_{\{0\}}(\beta)\Gamma(d_1)}{d_2^{d_1}}\ \text{IG}(d_1,d_2) + \frac{q\Gamma(d_1+\frac{1}{2})}{\sqrt{2\pi}\sigma_\epsilon\left(d_2+\frac{\beta^2}{2\sigma_\epsilon^2}\right)^{d_1+\frac{1}{2}}}\ \text{IG}\left(d_1+\frac{1}{2},\ d_2+\frac{\beta^2}{2\sigma_\epsilon^2}\right)
\end{aligned}
$$

Let us define

$$
\widetilde{w}_1 = \frac{(1-q)\delta_{\{0\}}(\beta)\Gamma(d_1)}{d_2^{d_1}} \quad \text{and} \quad \widetilde{w}_2 = \frac{q\Gamma(d_1+\frac{1}{2})}{\sqrt{2\pi}\sigma_\epsilon\left(d_2+\frac{\beta^2}{2\sigma_\epsilon^2}\right)^{d_1+\frac{1}{2}}}
$$

Then, $\sigma_d^2|\cdot$ follows a mixture of inverse gamma distributions, given by

$$
\sigma_d^2|\cdot \sim \begin{cases} \text{IG}(d_1,d_2), & \text{with probability} \quad \frac{\widetilde{w}_1}{\widetilde{w}_1+\widetilde{w}_2} \\ \text{IG}\left(d_1+\frac{1}{2},\ d_2+\frac{\beta^2}{2\sigma_\epsilon^2}\right), & \text{with probability} \quad \frac{\widetilde{w}_2}{\widetilde{w}_1+\widetilde{w}_2} \end{cases} \tag{7}
$$

#### 2.8 Full conditional distribution of $\sigma_\epsilon^2$

The full conditional distribution of $\sigma_\epsilon^2$ is given by

$$
\begin{aligned}
f(\sigma_\epsilon^2|\cdot) &\propto f(\mathbf{y}|\boldsymbol{\alpha},\beta,\sigma_\epsilon^2)f(\beta|q,\sigma_d^2,\sigma_\epsilon^2)f(\sigma_\epsilon^2)\\
&\propto \left\{\left(\sigma_\epsilon^2\right)^{-\frac{N}{2}}\exp\left[-\frac{1}{2\sigma_\epsilon^2}(\mathbf{y}-\beta G\boldsymbol{\alpha})^{\mathrm{T}}(\mathbf{y}-\beta G\boldsymbol{\alpha})\right]\right\}\left\{(1-q)\delta_{\{0\}}(\beta)+\frac{q}{\sqrt{2\pi}\sigma_d\sigma_\epsilon}\exp\left[-\frac{\beta^2}{2\sigma_d^2\sigma_\epsilon^2}\right]\right\}\\
&\quad \left\{(\sigma_\epsilon^2)^{-(\epsilon_1+1)}\exp\left[-\frac{\epsilon_2}{\sigma_\epsilon^2}\right]\right\}\\
&\propto \left\{(1-q)\delta_{\{0\}}(\beta)(\sigma_\epsilon^2)^{-(\frac{N}{2}+\epsilon_1+1)}\exp\left[-\frac{1}{\sigma_\epsilon^2}\left(\epsilon_2+\frac{1}{2}(\mathbf{y}-\beta G\boldsymbol{\alpha})^{\mathrm{T}}(\mathbf{y}-\beta G\boldsymbol{\alpha})\right)\right]+\right.\\
&\quad \left.\frac{q}{\sqrt{2\pi}\sigma_d}(\sigma_\epsilon^2)^{-(\frac{N+1}{2}+\epsilon_1+1)}\exp\left[-\frac{1}{\sigma_\epsilon^2}\left(\epsilon_2+\frac{\beta^2}{2\sigma_d^2}+\frac{1}{2}(\mathbf{y}-\beta G\boldsymbol{\alpha})^{\mathrm{T}}(\mathbf{y}-\beta G\boldsymbol{\alpha})\right)\right]\right\}\\
&\propto \left\{\widetilde{w}_1\ \mathrm{IG}\left(\frac{N}{2}+\epsilon_1,\ \epsilon_2+\frac{1}{2}(\mathbf{y}-\beta G\boldsymbol{\alpha})^{\mathrm{T}}(\mathbf{y}-\beta G\boldsymbol{\alpha})\right)+\right.\\
&\qquad\qquad \left.\widetilde{w}_2\ \mathrm{IG}\left(\frac{N+1}{2}+\epsilon_1,\ \epsilon_2+\frac{\beta^2}{2\sigma_d^2}+\frac{1}{2}(\mathbf{y}-\beta G\boldsymbol{\alpha})^{\mathrm{T}}(\mathbf{y}-\beta G\boldsymbol{\alpha})\right)\right\};
\end{aligned}
$$

where

$$
\begin{aligned}
\widetilde{w}_1 &= \frac{(1-q)\delta_{\{0\}}(\beta)\Gamma(\frac{N}{2}+\epsilon_1)}{\left(\epsilon_2+\frac{1}{2}(\mathbf{y}-\beta G\boldsymbol{\alpha})^{\mathrm{T}}(\mathbf{y}-\beta G\boldsymbol{\alpha})\right)^{\frac{N}{2}+\epsilon_1}}\\
\widetilde{w}_2 &= \frac{q\Gamma(\frac{N+1}{2}+\epsilon_1)}{\sqrt{2\pi}\sigma_d\left(\epsilon_2+\frac{\beta^2}{2\sigma_d^2}+\frac{1}{2}(\mathbf{y}-\beta G\boldsymbol{\alpha})^{\mathrm{T}}(\mathbf{y}-\beta G\boldsymbol{\alpha})\right)^{\frac{N+1}{2}+\epsilon_1}}
\end{aligned}
$$

Thus $\sigma_\epsilon^2|\cdot$ follows a mixture of inverse gamma distributions, given by

$$
\sigma_\epsilon^2|\cdot \sim \begin{cases}\mathrm{IG}\left(\frac{N}{2}+\epsilon_1,\ \epsilon_2+\frac{1}{2}(\mathbf{y}-\beta G\boldsymbol{\alpha})^{\mathrm{T}}(\mathbf{y}-\beta G\boldsymbol{\alpha})\right), & \text{with probability } \frac{\widetilde{w}_1}{\widetilde{w}_1+\widetilde{w}_2}\\ \mathrm{IG}\left(\frac{N+1}{2}+\epsilon_1,\ \epsilon_2+\frac{\beta^2}{2\sigma_d^2}+\frac{1}{2}(\mathbf{y}-\beta G\boldsymbol{\alpha})^{\mathrm{T}}(\mathbf{y}-\beta G\boldsymbol{\alpha})\right), & \text{with probability } \frac{\widetilde{w}_2}{\widetilde{w}_1+\widetilde{w}_2}\end{cases}
$$

### 3 Metropolis-Hastings algorithm for shrinkage parameters in single-ancestry setup

We make inferences on the model parameters by implementing the Gibbs sampler. However, we cannot sample directly from the full conditional distributions of shrinkage parameters $\boldsymbol{\lambda}$ and $\tau$ as these distributions are not of any well-known form. As a remedy, we implement the

Metropolis-Hastings steps within the Gibbs sampler to draw samples from these conditional posterior distributions.

#### 3.1 Metropolis-Hastings algorithm for $\boldsymbol{\lambda}$

We derive the Metropolis-Hastings algorithm for $\lambda_j$, the $j^{\text{th}}$ component of the vector $\boldsymbol{\lambda}$. The derivations of the other components can be done similarly. The full conditional posterior distribution of $\lambda_j$ (Equation 3) is given by

$$f(\lambda_j|\cdot) \propto \frac{1}{\lambda_j(1+\lambda_j^2)} \ \exp\left[-\frac{\alpha_j^2}{2\sigma_e^2\tau^2\lambda_j^2}\right], \quad \lambda_j > 0$$

Consider the transformation $Z_j = \lambda_j^2$. Then the conditional posterior distribution of $Z_j$ turns out to be

$$f(Z_j|\cdot) \propto \frac{1}{Z_j(1+Z_j)} \ \exp\left[-\frac{\alpha_j^2}{2\sigma_e^2\tau^2 Z_j}\right], \quad Z_j > 0$$

The transformation $Z_j = \lambda_j^2$ is one-to-one since $\lambda_j > 0$. Hence, updating the parameter $\lambda_j$ is equivalent to updating $Z_j$ in an MCMC step. Let us denote the density of $Z_j|\cdot$ by $g(\cdot)$. We choose the candidate proposal distribution to be IG $\left(1, \frac{\alpha_j^2}{2\sigma_e^2\tau^2}\right)$ and denote its density by $h(\cdot)$. This proposal density approximates the target distribution well for large values of $Z_j$. Because the proposal distribution does not depend on previously generated samples from the target distribution at any step, an independent Metropolis-Hastings algorithm is used to update $Z_j$ in the MCMC.

Let ${z_j}^{(l-1)}$ be the sample generated from the target distribution $Z_j|\cdot$ at the $(l-1)^{\text{th}}$ step. We consider the following steps to draw a sample from the target distribution at the $l^{\text{th}}$ step.

- Generate a random sample $v^{(l)}$ from the IG $\left(1, \frac{\alpha_j^2}{2\sigma_e^2\tau^2}\right)$ distribution.

- Draw a random number $U \sim \text{Uniform}(0,1)$.

- Set

$$z_j{}^{(l)} = \begin{cases} v^{(l)}, & \text{if } 0 < U < r(z_j{}^{(l-1)}, v^{(l)}) \\ z_j{}^{(l-1)}, & \text{if } U > r(z_j{}^{(l-1)}, v^{(l)}) \end{cases}$$

Then $z_j{}^{(l)}$ is the sample generated from the target distribution at the $l^{\text{th}}$ step. The quantity $r(z_j{}^{(l-1)}, v^{(l)})$ is the acceptance ratio of the Metropolis-Hastings algorithm, whose analytic form is given by

$$\begin{aligned} r(z_j{}^{(l-1)}, v^{(l)}) &= \min\left\{1, \frac{g(v^{(l)})h(z_j{}^{(l-1)})}{g(z_j{}^{(l-1)})h(v^{(l)})}\right\} \\ &= \min\left\{1, \frac{\frac{1}{v^{(l)}\left(1+v^{(l)}\right)} \exp\left[-\frac{\alpha_j^2}{2\sigma_e^2\tau^2 v^{(l)}}\right] \frac{1}{\left(z_j{}^{(l-1)}\right)^2} \exp\left[-\frac{\alpha_j^2}{2\sigma_e^2\tau^2 z_j{}^{(l-1)}}\right]}{\frac{1}{z_j{}^{(l-1)}\left(1+z_j{}^{(l-1)}\right)} \exp\left[-\frac{\alpha_j^2}{2\sigma_e^2\tau^2 z_j{}^{(l-1)}}\right] \frac{1}{\left(v^{(l)}\right)^2} \exp\left[-\frac{\alpha_j^2}{2\sigma_e^2\tau^2 v^{(l)}}\right]}\right\} \\ &= \min\left\{1, \frac{1+z_j{}^{(l-1)}}{z_j{}^{(l-1)}} \frac{v^{(l)}}{1+v^{(l)}}\right\} \end{aligned}$$

At the $l^{\text{th}}$ step, we either draw a new sample $v^{(l)}$ from the target distribution with probability $r(z_j{}^{(l-1)}, v^{(l)})$ or we retain the previous sample $z_j{}^{(l-1)}$ with probability $1 - r(z_j{}^{(l-1)}, v^{(l)})$. Thus $\sqrt{z_j{}^{(l)}}$ is the sample drawn from the full conditional posterior distribution of $\lambda_j|\cdot$ at the $l^{\text{th}}$ step.

#### 3.2 Metropolis-Hastings algorithm for $\tau$

The full conditional posterior distribution of $\tau$ (Equation 4) is given by

$$f(\tau|\cdot) \propto \frac{1}{\tau^m(1+\tau^2)} \exp\left[-\frac{\boldsymbol{\alpha}^{\mathrm{T}} D_\lambda^{-1} \boldsymbol{\alpha}}{2\sigma_e^2\tau^2}\right], \quad \tau > 0$$

Consider the transformation $Z = \tau^2$. Then the conditional posterior distribution of $Z$ becomes

$$f(Z|\cdot) \propto \frac{1}{Z^{\frac{m+1}{2}}(1+Z)} \exp\left[-\frac{\boldsymbol{\alpha}^{\mathrm{T}} D_\lambda^{-1} \boldsymbol{\alpha}}{2\sigma_e^2 Z}\right], \quad Z > 0$$

Since the parameters $\tau$ and $Z$ are related through the one-to-one transformation $Z = \tau^2$, updating $\tau$ is equivalent to updating $Z$. Let us denote the density of $Z|\cdot$ by $g(\cdot)$. We choose the candidate distribution as IG $\left(\frac{m+1}{2}, \frac{\boldsymbol{\alpha}^{\mathrm{T}} D_{\lambda}^{-1} \boldsymbol{\alpha}}{2\sigma_e^2}\right)$ and denote its density by $h(\cdot)$. The independent Metropolis-Hastings algorithm is then implemented to update $Z$ within an MCMC step.

Let $z^{(l-1)}$ be the sample drawn from the target distribution $Z|\cdot$ at the $(l-1)^{\text{th}}$ step. We perform the following steps to generate a sample from the target distribution at the $l^{\text{th}}$ step.

- Generate a random sample $v^{(l)}$ from the IG $\left(\frac{m+1}{2}, \frac{\boldsymbol{\alpha}^{\mathrm{T}} D_{\lambda}^{-1} \boldsymbol{\alpha}}{2\sigma_e^2}\right)$ distribution.
- Draw a random number $U \sim \text{Uniform}(0, 1)$.
- Set

$$z^{(l)} = \begin{cases} v^{(l)}, & \text{if } 0 < U < r(z^{(l-1)}, v^{(l)}) \\ z^{(l-1)}, & \text{if } U > r(z^{(l-1)}, v^{(l)}) \end{cases}$$

Then $z^{(l)}$ denotes the sample generated from the target distribution at the $l^{\text{th}}$ step. $r(z^{(l-1)}, v^{(l)})$ represents the Metropolis-Hastings acceptance ratio, whose analytic form is given below.

$$\begin{aligned} r(z^{(l-1)}, v^{(l)}) &= \min\left\{1, \frac{g(v^{(l)})h(z^{(l-1)})}{g(z^{(l-1)})h(v^{(l)})}\right\} \\ &= \min\left\{1, \frac{\frac{1}{\left(v^{(l)}\right)^{\frac{m+1}{2}}\left(1+v^{(l)}\right)} \exp\left[-\frac{\boldsymbol{\alpha}^{\mathrm{T}} D_{\lambda}^{-1} \boldsymbol{\alpha}}{2\sigma_e^2 v^{(l)}}\right] \frac{1}{\left(z^{(l-1)}\right)^{\frac{m+1}{2}+1}} \exp\left[-\frac{\boldsymbol{\alpha}^{\mathrm{T}} D_{\lambda}^{-1} \boldsymbol{\alpha}}{2\sigma_e^2 z^{(l-1)}}\right]}{\frac{1}{\left(z^{(l-1)}\right)^{\frac{m+1}{2}}\left(1+z^{(l-1)}\right)} \exp\left[-\frac{\boldsymbol{\alpha}^{\mathrm{T}} D_{\lambda}^{-1} \boldsymbol{\alpha}}{2\sigma_e^2 z^{(l-1)}}\right] \frac{1}{\left(v^{(l)}\right)^{\frac{m+1}{2}+1}} \exp\left[-\frac{\boldsymbol{\alpha}^{\mathrm{T}} D_{\lambda}^{-1} \boldsymbol{\alpha}}{2\sigma_e^2 v^{(l)}}\right]}\right\} \\ &= \min\left\{1, \frac{1+z^{(l-1)}}{z^{(l-1)}} \frac{v^{(l)}}{1+v^{(l)}}\right\} \end{aligned}$$

Then $\sqrt{z^{(l)}}$ is the random sample drawn from the full conditional distribution of $\tau|\cdot$ at the $l^{\text{th}}$ step.

### 4 Derivation of the full conditional distributions for glossTWAS in multi-ancestry TWAS setup

The hierarchical structure of glossTWAS in the multi-ancestry TWAS setting is given by

$$
\begin{aligned}
\mathbf{x}_{k,\text{ref}}|\boldsymbol{\alpha}_k, \sigma^2_{e_k} &\sim \text{N}_{n_k}(G_{k,\text{ref}}\boldsymbol{\alpha}_k, \sigma^2_{e_k}\text{I}_{n_k}), \quad k = 1, 2. \\
\mathbf{y}_k|\beta, \boldsymbol{\alpha}_k, w_k, \sigma^2_\epsilon &\sim \text{N}_{N_k}(\beta w_k G_k \boldsymbol{\alpha}_k, \sigma^2_\epsilon \text{I}_{N_k}) \\
\alpha_{kj}|\lambda_j, \tau, \sigma^2_{e_k} &\overset{ind}{\sim} \text{N}(0, \sigma^2_{e_k}\tau^2\lambda^2_j), \quad j = 1(1)m \\
\lambda_j &\overset{ind}{\sim} \text{C}^+(0, 1) \\
\tau &\sim \text{C}^+(0, 1) \\
w &\sim \text{Beta}(\gamma_1, \gamma_2), \quad \gamma_1 = \gamma_2 = 1 \\
\beta|q, \sigma^2_d, \sigma^2_\epsilon &\sim (1-q)\delta_{\{0\}}(\beta) + q\text{N}(0, \sigma^2_d\sigma^2_\epsilon) \\
q &\sim \text{Beta}(b_1, b_2) \\
\sigma^2_d &\sim \text{IG}(d_1, d_2) \\
\sigma^2_{e_k} &\sim \text{IG}(e_1, e_2) \\
\sigma^2_\epsilon &\sim \text{IG}(\epsilon_1, \epsilon_2)
\end{aligned}
\tag{8}
$$

The term $w_k$ in the Equation 8 equals $w$ for $k = 1$ and $(1 - w)$ for $k = 2$. For $k = 1, 2$, let us denote $\boldsymbol{\alpha}_k = (\alpha_{k1}, \alpha_{k2}, \cdots, \alpha_{km})^\text{T}$. For $j = 1(1)m$, $\alpha_{kj}$ denotes the effect of $j^\text{th}$ cis-SNP on the gene expression in the $k^\text{th}$ ancestry. We specify the SNP-specific shrinkage parameters across all ancestries in the form of a vector as $\boldsymbol{\lambda} = (\lambda_1, \lambda_2, \cdots, \lambda_m)^\text{T}$. Let us define $D_\lambda = diag(\lambda^2_1, \lambda^2_2, \cdots, \lambda^2_m)$, where $D_\lambda$ is a $m \times m$ diagonal matrix. The prior distribution of $\boldsymbol{\alpha}_k$ conditional on $\boldsymbol{\lambda}, \tau$ and $\sigma^2_{e_k}$ is given by

$$\boldsymbol{\alpha}_k|\boldsymbol{\lambda}, \tau, \sigma^2_{e_k} \sim \text{N}_m(\mathbf{0}, \sigma^2_{e_k}\tau^2 D_\lambda)$$

We now derive the full conditional posterior distributions of the model parameters.

#### 4.1 Full conditional distribution of $\boldsymbol{\alpha}_k$

The full conditional distribution of $\boldsymbol{\alpha}_k$ is given by

$$
\begin{aligned}
f(\boldsymbol{\alpha}_k|\cdot) &\propto f(\mathbf{x}_{k,\text{ref}}|\boldsymbol{\alpha}_k, \sigma^2_{e_k}) f(\mathbf{y}_k|\beta, w_k, \boldsymbol{\alpha}_k, \sigma^2_\epsilon) f(\boldsymbol{\alpha}_k|\boldsymbol{\lambda}, \tau, \sigma^2_{e_k}) \\
&\propto \exp\left[-\frac{1}{2\sigma^2_{e_k}}(\mathbf{x}_{k,\text{ref}} - G_{k,\text{ref}}\boldsymbol{\alpha}_k)^{\mathrm{T}}(\mathbf{x}_{k,\text{ref}} - G_{k,\text{ref}}\boldsymbol{\alpha}_k)\right] \exp\left[-\frac{1}{2\sigma^2_\epsilon}(\mathbf{y}_k - \beta w_k G_k \boldsymbol{\alpha}_k)^{\mathrm{T}}(\mathbf{y}_k - \beta w_k G_k \boldsymbol{\alpha}_k)\right] \\
&\quad \exp\left[-\frac{\boldsymbol{\alpha}_k^{\mathrm{T}} D_\lambda^{-1}\boldsymbol{\alpha}_k}{2\sigma^2_{e_k}\tau^2}\right] \\
&\propto \exp\left\{-\frac{1}{2}\left[\boldsymbol{\alpha}_k^{\mathrm{T}}\left(\frac{1}{\sigma^2_{e_k}}G^{\mathrm{T}}_{k,\text{ref}}G_{k,\text{ref}} + \frac{\beta^2 w_k^2}{\sigma^2_\epsilon}G_k^{\mathrm{T}}G_k + \frac{1}{\sigma^2_{e_k}\tau^2}D_\lambda^{-1}\right)\boldsymbol{\alpha}_k - 2\boldsymbol{\alpha}_k^{\mathrm{T}}\left(\frac{1}{\sigma^2_{e_k}}G^{\mathrm{T}}_{k,\text{ref}}\mathbf{x}_{k,\text{ref}} + \frac{\beta w_k}{\sigma^2_\epsilon}G_k^{\mathrm{T}}\mathbf{y}_k\right)\right]\right\} \\
&\propto \exp\left\{-\frac{1}{2}\left[\boldsymbol{\alpha}_k^{\mathrm{T}}\Sigma_k\boldsymbol{\alpha}_k - 2\boldsymbol{\alpha}_k^{\mathrm{T}}\left(\frac{1}{\sigma^2_{e_k}}G^{\mathrm{T}}_{k,\text{ref}}\mathbf{x}_{k,\text{ref}} + \frac{\beta w_k}{\sigma^2_\epsilon}G_k^{\mathrm{T}}\mathbf{y}_k\right)\right]\right\};
\end{aligned}
$$

where $\Sigma_k = \frac{1}{\sigma^2_{e_k}}G^{\mathrm{T}}_{k,\text{ref}}G_{k,\text{ref}} + \frac{\beta^2 w_k^2}{\sigma^2_\epsilon}G_k^{\mathrm{T}}G_k + \frac{1}{\sigma^2_{e_k}\tau^2}D_\lambda^{-1}$

Hence, the full conditional distribution of $\boldsymbol{\alpha}_k$ is given by

$$
\boldsymbol{\alpha}_k|\cdot \sim \mathrm{N}_m\left(\Sigma_k^{-1}\left(\frac{1}{\sigma^2_{e_k}}G^{\mathrm{T}}_{k,\text{ref}}\mathbf{x}_{k,\text{ref}} + \frac{\beta w_k}{\sigma^2_\epsilon}G_k^{\mathrm{T}}\mathbf{y}_k\right), \Sigma_k^{-1}\right), \quad k = 1, 2.
$$

#### 4.2 Full conditional distribution of $\boldsymbol{\lambda}$

For each $j = 1(1)m$, since the $\lambda_j$s are independently distributed under the prior, it is convenient to update the components of $\boldsymbol{\lambda}$ sequentially within the MCMC step. The conditional posterior distribution of $\lambda_j$ is given by

$$
\begin{aligned}
f(\lambda_j|\cdot) &\propto \left\{\prod_{k=1}^{2} f(\alpha_{kj}|\lambda_j, \tau, \sigma^2_e)\right\} f(\lambda_j), \quad j = 1(1)m \\
&\propto \left[\prod_{k=1}^{2}\left\{\frac{1}{\sigma_{e_k}\lambda_j\tau}\ \exp\left[-\frac{\alpha^2_{kj}}{2\sigma^2_{e_k}\tau^2\lambda_j^2}\right]\right\}\right]\left\{\frac{1}{1+\lambda_j^2}\right\} \qquad (9) \\
&\propto \frac{1}{\lambda_j^2(1+\lambda_j^2)}\ \exp\left[-\frac{1}{2\tau^2\lambda_j^2}\sum_{k=1}^{2}\frac{\alpha^2_{kj}}{\sigma^2_{e_k}}\right]
\end{aligned}
$$

Since this full conditional posterior distribution of $\lambda_j$ is not conjugate, we employ the Metropolis-Hastings algorithm to generate posterior samples. A brief description of the algorithm is provided in section 5.1.

#### 4.3 Full conditional distribution of $\tau$

The full conditional distribution of $\tau$ is given by

$$\begin{aligned} f(\tau|\cdot) &\propto \left\{\prod_{k=1}^{2} f(\boldsymbol{\alpha}_k|\boldsymbol{\lambda},\tau,\sigma^2_{e_k})\right\} f(\tau) \\ &\propto \left\{\prod_{k=1}^{2} \frac{1}{\tau^m}\ \exp\left[-\frac{\boldsymbol{\alpha}_k^{\mathrm{T}} D_\lambda^{-1}\boldsymbol{\alpha}_k}{2\sigma^2_{e_k}\tau^2}\right]\right\}\left\{\frac{1}{1+\tau^2}\right\} \\ &\propto \frac{1}{\tau^{2m}(1+\tau^2)}\ \exp\left[-\frac{1}{2\tau^2}\sum_{k=1}^{2}\frac{\boldsymbol{\alpha}_k^{\mathrm{T}} D_\lambda^{-1}\boldsymbol{\alpha}_k}{\sigma^2_{e_k}}\right] \end{aligned} \tag{10}$$

We employ the Metropolis-Hastings algorithm to draw samples from the conditional posterior distribution of $\tau$. The steps of the algorithm are described in section 5.2.

#### 4.4 Full conditional distribution of $\sigma^2_{e_k}$

For $k = 1, 2$, the full conditional distribution of $\sigma^2_{e_k}$ is given by

$$\begin{aligned} f(\sigma^2_{e_k}|\cdot) &\propto f(\mathbf{x}_{k,\mathrm{ref}}|\boldsymbol{\alpha}_k,\sigma^2_{e_k}) f(\boldsymbol{\alpha}_k|\boldsymbol{\lambda},\tau,\sigma^2_{e_k}) f(\sigma^2_{e_k}) \\ &\propto \left\{(\sigma^2_{e_k})^{-\frac{n_k}{2}} \exp\left[-\frac{1}{2\sigma^2_{e_k}}(\mathbf{x}_{k,\mathrm{ref}}-G_{k,\mathrm{ref}}\boldsymbol{\alpha}_k)^{\mathrm{T}}(\mathbf{x}_{k,\mathrm{ref}}-G_{k,\mathrm{ref}}\boldsymbol{\alpha}_k)\right]\right\}\left\{(\sigma^2_{e_k})^{-\frac{m}{2}}\exp\left[-\frac{\boldsymbol{\alpha}_k^{\mathrm{T}} D_\lambda^{-1}\boldsymbol{\alpha}_k}{2\sigma^2_{e_k}\tau^2}\right]\right\} \\ &\quad \left\{(\sigma^2_{e_k})^{-(e_1+1)}\exp\left[-\frac{e_2}{\sigma^2_{e_k}}\right]\right\} \\ &\propto (\sigma^2_{e_k})^{-\left(e_1+\frac{n_k}{2}+\frac{m}{2}+1\right)} \exp\left[-\frac{1}{\sigma^2_{e_k}}\left(e_2+\frac{\boldsymbol{\alpha}_k^{\mathrm{T}} D_\lambda^{-1}\boldsymbol{\alpha}_k}{2\tau^2}+\frac{1}{2}(\mathbf{x}_{k,\mathrm{ref}}-G_{k,\mathrm{ref}}\boldsymbol{\alpha}_k)^{\mathrm{T}}\left(\mathbf{x}_{k,\mathrm{ref}}-G_{k,\mathrm{ref}}\boldsymbol{\alpha}_k\right)\right)\right] \end{aligned}$$

Thus the full conditional distribution of $\sigma^2_{e_k}$ is given by

$$\sigma^2_{e_k}|\cdot \sim \mathrm{IG}\left(e_1+\frac{n_k}{2}+\frac{m}{2},\ e_2+\frac{\boldsymbol{\alpha}_k^{\mathrm{T}} D_\lambda^{-1}\boldsymbol{\alpha}_k}{2\tau^2}+\frac{1}{2}(\mathbf{x}_{k,\mathrm{ref}}-G_{k,\mathrm{ref}}\boldsymbol{\alpha}_k)^{\mathrm{T}}\left(\mathbf{x}_{k,\mathrm{ref}}-G_{k,\mathrm{ref}}\boldsymbol{\alpha}_k\right)\right), \quad k=1,2.$$

##### 4.5 Full conditional distribution of $\beta$

The full conditional distribution of $\beta$ is given by

$$
\begin{aligned}
f(\beta|\cdot) &\propto \left\{\prod_{k=1}^{2} f(\mathbf{y}_k|\beta, w_k, \boldsymbol{\alpha}_k, \sigma_\epsilon^2)\right\} f(\beta|q, \sigma_d^2, \sigma_\epsilon^2) \\
&\propto \left\{\prod_{k=1}^{2} \exp\left[-\frac{1}{2\sigma_\epsilon^2}(\mathbf{y}_k - \beta w_k G_k \boldsymbol{\alpha}_k)^{\mathrm{T}}(\mathbf{y}_k - \beta w_k G_k \boldsymbol{\alpha}_k)\right]\right\} \left\{(1-q)\delta_{\{0\}}(\beta) + \frac{q}{\sqrt{2\pi}\sigma_d\sigma_\epsilon} \exp\left[-\frac{\beta^2}{2\sigma_d^2\sigma_\epsilon^2}\right]\right\} \\
&\propto \exp\left[-\frac{1}{2\sigma_\epsilon^2}\sum_{k=1}^{2}\left(\beta^2 w_k^2 \boldsymbol{\alpha}_k^{\mathrm{T}} G_k^{\mathrm{T}} G_k \boldsymbol{\alpha}_k - 2\beta w_k \boldsymbol{\alpha}_k^{\mathrm{T}} G_k^{\mathrm{T}} \mathbf{y}_k\right)\right] \left\{(1-q)\delta_{\{0\}}(\beta) + \frac{q}{\sqrt{2\pi}\sigma_d\sigma_\epsilon} \exp\left[-\frac{\beta^2}{2\sigma_d^2\sigma_\epsilon^2}\right]\right\} \\
&\propto (1-q)\delta_{\{0\}}(\beta) + \frac{q}{\sqrt{2\pi}\sigma_d\sigma_\epsilon} \exp\left[-\frac{\sum_{k=1}^{2} w_k^2 \boldsymbol{\alpha}_k^{\mathrm{T}} G_k^{\mathrm{T}} G_k \boldsymbol{\alpha}_k + \frac{1}{\sigma_d^2}}{2\sigma_\epsilon^2}\left(\beta^2 - 2\beta\,\frac{\sum_{k=1}^{2} w_k \boldsymbol{\alpha}_k^{\mathrm{T}} G_k^{\mathrm{T}} \mathbf{y}_k}{\sum_{k=1}^{2} w_k^2 \boldsymbol{\alpha}_k^{\mathrm{T}} G_k^{\mathrm{T}} G_k \boldsymbol{\alpha}_k + \frac{1}{\sigma_d^2}}\right)\right] \\
&\propto (1-q)\delta_{\{0\}}(\beta) + \frac{q}{\sqrt{2\pi}\sigma_d\sigma_\epsilon} \exp\left[-\frac{1}{2\widetilde{\sigma}^2}(\beta^2 - 2\beta\widetilde{\mu})\right] \\
&\propto (1-q)\delta_{\{0\}}(\beta) + \left(\frac{q\widetilde{\sigma}}{\sigma_d\sigma_\epsilon}\exp\left[\frac{\widetilde{\mu}^2}{2\widetilde{\sigma}^2}\right]\right)\left\{\frac{1}{\sqrt{2\pi}\widetilde{\sigma}}\exp\left[-\frac{1}{2\widetilde{\sigma}^2}(\beta - \widetilde{\mu})^2\right]\right\};
\end{aligned}
$$

where $\widetilde{\mu} = \dfrac{\sum_{k=1}^{2} w_k \boldsymbol{\alpha}_k^{\mathrm{T}} G_k^{\mathrm{T}} \mathbf{y}_k}{\sum_{k=1}^{2} w_k^2 \boldsymbol{\alpha}_k^{\mathrm{T}} G_k^{\mathrm{T}} G_k \boldsymbol{\alpha}_k + \frac{1}{\sigma_d^2}}$ and $\widetilde{\sigma}^2 = \dfrac{\sigma_\epsilon^2}{\sum_{k=1}^{2} w_k^2 \boldsymbol{\alpha}_k^{\mathrm{T}} G_k^{\mathrm{T}} G_k \boldsymbol{\alpha}_k + \frac{1}{\sigma_d^2}}$

Hence, the full conditional distribution of $\beta$ is a mixture of point mass function $\delta_{\{0\}}(\beta)$ and normal distribution $\mathrm{N}(\widetilde{\mu}, \widetilde{\sigma}^2)$. Let us denote the mixing probability attached to the point mass function $\delta_{\{0\}}(\beta)$ by

$$
\widetilde{w} = \frac{1-q}{1-q+\frac{q\widetilde{\sigma}}{\sigma_d\sigma_\epsilon}\exp\left(\frac{\widetilde{\mu}^2}{2\widetilde{\sigma}^2}\right)} \tag{11}
$$

Hence, the conditional posterior distribution of $\beta$ becomes

$$
\beta|\cdot \sim \begin{cases} \delta_{\{0\}}(\beta), & \text{with probability} \quad \widetilde{w} \\ \mathrm{N}(\widetilde{\mu}, \widetilde{\sigma}^2), & \text{with probability} \quad 1-\widetilde{w} \end{cases}
$$

#### 4.6 Full conditional distribution of $\sigma_\epsilon^2$

Let us denote $N_c = \sum_{k=1}^2 N_k$. The full conditional distribution of $\sigma_\epsilon^2$ is given by

$$
\begin{aligned}
f(\sigma_\epsilon^2|\cdot) &\propto \left\{\prod_{k=1}^{2} f(\mathbf{y}_k|\beta, w_k, \boldsymbol{\alpha}_k, \sigma_\epsilon^2)\right\} f(\beta|q,\sigma_d^2,\sigma_\epsilon^2) f(\sigma_\epsilon^2) \\
&\propto \left\{\prod_{k=1}^{2} (\sigma_\epsilon^2)^{-\frac{N_k}{2}} \exp\left[-\frac{1}{2\sigma_\epsilon^2}(\mathbf{y}_k - \beta w_k G_k \boldsymbol{\alpha}_k)^{\mathrm{T}}(\mathbf{y}_k - \beta w_k G_k \boldsymbol{\alpha}_k)\right]\right\} \\
&\quad \left\{(1-q)\delta_{\{0\}}(\beta) + \frac{q}{\sqrt{2\pi}\sigma_d\sigma_\epsilon} \exp\left[-\frac{\beta^2}{2\sigma_d^2\sigma_\epsilon^2}\right]\right\}\left\{(\sigma_\epsilon^2)^{-(\epsilon_1+1)}\exp\left[-\frac{\epsilon_2}{\sigma_\epsilon^2}\right]\right\} \\
&\propto \left\{(1-q)\delta_{\{0\}}(\beta)(\sigma_\epsilon^2)^{-(\frac{N_c}{2}+\epsilon_1+1)} \exp\left[-\frac{1}{\sigma_\epsilon^2}\left(\epsilon_2 + \frac{1}{2}\sum_{k=1}^{2}(\mathbf{y}_k - \beta w_k G_k \boldsymbol{\alpha}_k)^{\mathrm{T}}(\mathbf{y}_k - \beta w_k G_k \boldsymbol{\alpha}_k)\right)\right] + \right. \\
&\quad \left. \frac{q}{\sqrt{2\pi}\sigma_d}(\sigma_\epsilon^2)^{-(\frac{N_c+1}{2}+\epsilon_1+1)} \exp\left[-\frac{1}{\sigma_\epsilon^2}\left(\epsilon_2 + \frac{\beta^2}{2\sigma_d^2} + \frac{1}{2}\sum_{k=1}^{2}(\mathbf{y}_k - \beta w_k G_k \boldsymbol{\alpha}_k)^{\mathrm{T}}(\mathbf{y}_k - \beta w_k G_k \boldsymbol{\alpha}_k)\right)\right]\right\} \\
&\propto \left\{\widetilde{w}_1 \ \mathrm{IG}\left(\frac{N_c}{2} + \epsilon_1,\ \epsilon_2 + \frac{1}{2}\sum_{k=1}^{2}(\mathbf{y}_k - \beta w_k G_k \boldsymbol{\alpha}_k)^{\mathrm{T}}(\mathbf{y}_k - \beta w_k G_k \boldsymbol{\alpha}_k)\right) + \right. \\
&\quad \left. \widetilde{w}_2 \ \mathrm{IG}\left(\frac{N_c+1}{2} + \epsilon_1,\ \epsilon_2 + \frac{\beta^2}{2\sigma_d^2} + \frac{1}{2}\sum_{k=1}^{2}(\mathbf{y}_k - \beta w_k G_k \boldsymbol{\alpha}_k)^{\mathrm{T}}(\mathbf{y}_k - \beta w_k G_k \boldsymbol{\alpha}_k)\right)\right\};
\end{aligned}
$$

where

$$
\begin{aligned}
\widetilde{w}_1 &= \frac{(1-q)\delta_{\{0\}}(\beta)\Gamma(\frac{N_c}{2}+\epsilon_1)}{\left(\epsilon_2 + \frac{1}{2}\sum_{k=1}^{2}(\mathbf{y}_k - \beta w_k G_k \boldsymbol{\alpha}_k)^{\mathrm{T}}(\mathbf{y}_k - \beta w_k G_k \boldsymbol{\alpha}_k)\right)^{\frac{N_c}{2}+\epsilon_1}} \\
\widetilde{w}_2 &= \frac{q\ \Gamma(\frac{N_c+1}{2}+\epsilon_1)}{\sqrt{2\pi}\sigma_d\left(\epsilon_2 + \frac{\beta^2}{2\sigma_d^2} + \frac{1}{2}\sum_{k=1}^{2}(\mathbf{y}_k - \beta w_k G_k \boldsymbol{\alpha}_k)^{\mathrm{T}}(\mathbf{y}_k - \beta w_k G_k \boldsymbol{\alpha}_k)\right)^{\frac{N_c+1}{2}+\epsilon_1}}
\end{aligned}
$$

Thus, the full conditional distribution of $\sigma_\epsilon^2$ can be represented as a mixture of inverse gamma distributions as follows

$$
\sigma_\epsilon^2|\cdot \sim \begin{cases} \mathrm{IG}\left(\frac{N_c}{2}+\epsilon_1,\ \epsilon_2 + \frac{1}{2}\sum_{k=1}^{2}(\mathbf{y}_k - \beta w_k G_k \boldsymbol{\alpha}_k)^{\mathrm{T}}(\mathbf{y}_k - \beta w_k G_k \boldsymbol{\alpha}_k)\right), & \text{with probability } \frac{\widetilde{w}_1}{\widetilde{w}_1+\widetilde{w}_2} \\ \mathrm{IG}\left(\frac{N_c+1}{2}+\epsilon_1,\ \epsilon_2 + \frac{\beta^2}{2\sigma_d^2} + \frac{1}{2}\sum_{k=1}^{2}(\mathbf{y}_k - \beta w_k G_k \boldsymbol{\alpha}_k)^{\mathrm{T}}(\mathbf{y}_k - \beta w_k G_k \boldsymbol{\alpha}_k)\right), & \text{with probability } \frac{\widetilde{w}_2}{\widetilde{w}_1+\widetilde{w}_2} \end{cases}
$$

#### 4.7 Full conditional distribution of $w$

The full conditional distribution of $w$ is given by

$$
\begin{aligned}
f(w|\cdot) &\propto f(\mathbf{y_1}|\beta, w, \boldsymbol{\alpha}_1, \sigma_\epsilon^2) f(\mathbf{y_2}|\beta, w, \boldsymbol{\alpha}_2, \sigma_\epsilon^2) f(w) \\
&\propto \exp\left[-\frac{1}{2\sigma_\epsilon^2}(\mathbf{y}_1 - \beta w G_1\boldsymbol{\alpha}_1)^{\mathrm{T}}(\mathbf{y}_1 - \beta w G_1\boldsymbol{\alpha}_1)\right] \exp\left[-\frac{1}{2\sigma_\epsilon^2}(\mathbf{y}_2 - \beta(1-w)G_2\boldsymbol{\alpha}_2)^{\mathrm{T}}(\mathbf{y}_2 - \beta(1-w)G_2\boldsymbol{\alpha}_2)\right] \\
&\propto \exp\left[-\frac{1}{2\sigma_\epsilon^2}\left\{\beta^2\Big(w^2\boldsymbol{\alpha}_1^{\mathrm{T}}G_1^{\mathrm{T}}G_1\boldsymbol{\alpha}_1 + (1-w)^2\boldsymbol{\alpha}_2^{\mathrm{T}}G_2^{\mathrm{T}}G_2\boldsymbol{\alpha}_2\Big) - 2\beta\Big(w\boldsymbol{\alpha}_1^{\mathrm{T}}G_1^{\mathrm{T}}\mathbf{y}_1 + (1-w)\boldsymbol{\alpha}_2^{\mathrm{T}}G_2^{\mathrm{T}}\mathbf{y}_2\Big)\right\}\right] \qquad (12)
\end{aligned}
$$

We implement the random walk Metropolis-Hastings algorithm to draw samples from the conditional distribution of $w$ outlined in section 5.3.

The full conditional posterior distributions of $q$ and $\sigma_d^2$ remain the same as those given in Equations 6 and 7.

### 5 Metropolis-Hastings algorithm for shrinkage parameters in multi-ancestry TWAS setup

In our multi-ancestry TWAS model, the shrinkage parameters in the horseshoe prior are given by $\boldsymbol{\lambda}$ and $\tau$. As the posterior distributions of these shrinkage parameters are not conjugate, direct sampling from them is not feasible. Therefore, we use the Metropolis-Hastings algorithm to generate samples from these conditional distributions.

#### 5.1 Metropolis-Hastings algorithm for $\boldsymbol{\lambda}$

We derive the Metropolis-Hastings algorithm for $\lambda_j$, the $j^{\text{th}}$ component of the shrinkage parameter vector $\boldsymbol{\lambda}$. The derivation for the other components can be done analogously. The full conditional posterior distribution of $\lambda_j$ (Equation 9) is given by

$$
f(\lambda_j|\cdot) \propto \frac{1}{\lambda_j^2(1+\lambda_j^2)} \exp\left[-\frac{1}{2\tau^2\lambda_j^2}\sum_{k=1}^{2}\frac{\alpha_{kj}^2}{\sigma_{e_k}^2}\right], \quad \lambda_j > 0
$$

Consider the transformation $Z_j = \lambda_j^2$. Then the conditional posterior distribution of $Z_j$ becomes

$$f(Z_j|\cdot) \propto \frac{1}{Z_j^{\frac{3}{2}}(1+Z_j)} \; \exp\left[-\frac{1}{2\tau^2 Z_j}\sum_{k=1}^{2}\frac{\alpha_{kj}^2}{\sigma_{e_k}^2}\right], \quad Z_j > 0$$

Here, updating the parameter $\lambda_j$ is equivalent to updating $Z_j$, since they are related through the one-to-one transformation $Z_j = \lambda_j^2$. Let us denote the density of $Z_j|\cdot$ by $g(\cdot)$. We choose the candidate proposal distribution to be IG $\left(\frac{3}{2}, \frac{1}{2\tau^2}\sum_{k=1}^{2}\frac{\alpha_{kj}^2}{\sigma_{e_k}^2}\right)$ and denote its density by $h(\cdot)$. We follow the same procedure to generate a sample from the target distribution as discussed in the section 3.1. The resulting expression for the Metropolis-Hastings acceptance ratio is

$$r({z_j}^{(l-1)}, v^{(l)}) = \min\left\{1, \frac{g(v^{(l)})h({z_j}^{(l-1)})}{g({z_j}^{(l-1)})h(v^{(l)})}\right\} = \min\left\{1, \frac{1+{z_j}^{(l-1)}}{{z_j}^{(l-1)}}\frac{v^{(l)}}{1+v^{(l)}}\right\}; \qquad (13)$$

where ${z_j}^{(l-1)}$ is the generated sample from the target distribution at the $(l-1)^{\text{th}}$ step and $v^{(l)}$ is the sample from the proposal distribution at the $l^{\text{th}}$ step.

##### 5.2 Metropolis-Hastings algorithm for $\tau$

The full conditional posterior distribution of $\tau$ (Equation 10) is given by

$$f(\tau|\cdot) \propto \frac{1}{\tau^{2m}(1+\tau^2)} \; \exp\left[-\frac{1}{2\tau^2}\sum_{k=1}^{2}\frac{\boldsymbol{\alpha}_k^{\mathrm{T}} D_\lambda^{-1}\boldsymbol{\alpha}_k}{\sigma_{e_k}^2}\right], \quad \tau > 0$$

Consider the transformation $Z = \tau^2$. Then the conditional posterior distribution of $Z$ becomes

$$f(Z|\cdot) \propto \frac{1}{Z^{(m+\frac{1}{2})}(1+Z)} \; \exp\left[-\frac{1}{2Z}\sum_{k=1}^{2}\frac{\boldsymbol{\alpha}_k^{\mathrm{T}} D_\lambda^{-1}\boldsymbol{\alpha}_k}{\sigma_{e_k}^2}\right], \quad Z > 0$$

We choose the candidate proposal distribution as IG $\left(m+\frac{1}{2}, \frac{1}{2}\sum_{k=1}^{2}\frac{\boldsymbol{\alpha}_k^{\mathrm{T}} D_\lambda^{-1}\boldsymbol{\alpha}_k}{\sigma_{e_k}^2}\right)$. The expression for Metropolis-Hastings acceptance ratio coincides with the term presented in Equation 13 and the sampling procedure follows the same steps outlined in section 3.2.

##### 5.3 Metropolis-Hastings algorithm for $w$

The full conditional distribution of $w$ (Equation 12) is given by

$$f(w|\cdot) \propto \exp\left[-\frac{1}{2\sigma_\epsilon^2}\left\{\beta^2\Big(w^2\boldsymbol{\alpha}_1^{\mathrm{T}}G_1^{\mathrm{T}}G_1\boldsymbol{\alpha}_1 + (1-w)^2\boldsymbol{\alpha}_2^{\mathrm{T}}G_2^{\mathrm{T}}G_2\boldsymbol{\alpha}_2\Big) - 2\beta\Big(w\boldsymbol{\alpha}_1^{\mathrm{T}}G_1^{\mathrm{T}}\mathbf{y}_1 + (1-w)\boldsymbol{\alpha}_2^{\mathrm{T}}G_2^{\mathrm{T}}\mathbf{y}_2\Big)\right\}\right], \quad 0 < w < 1$$

We consider the logit transformation of $w$, i.e., $\theta = \log_e(\frac{w}{1-w})$. Then the conditional distribution of $\theta$ is given by

$$f(\theta|\cdot) \propto \exp\left[-\frac{1}{2\sigma_\epsilon^2}\left\{\beta^2\left(\left(\frac{e^\theta}{1+e^\theta}\right)^2\boldsymbol{\alpha}_1^{\mathrm{T}}G_1^{\mathrm{T}}G_1\boldsymbol{\alpha}_1 + \left(\frac{1}{1+e^\theta}\right)^2\boldsymbol{\alpha}_2^{\mathrm{T}}G_2^{\mathrm{T}}G_2\boldsymbol{\alpha}_2\right)\right.\right.$$
$$\left.\left. -2\beta\left(\frac{e^\theta}{1+e^\theta}\boldsymbol{\alpha}_1^{\mathrm{T}}G_1^{\mathrm{T}}\mathbf{y}_1 + \frac{1}{1+e^\theta}\boldsymbol{\alpha}_2^{\mathrm{T}}G_2^{\mathrm{T}}\mathbf{y}_2\right)\right\}\right]\frac{e^\theta}{(1+e^\theta)^2}, \quad -\infty < \theta < \infty$$

Since $\theta$ is related to $w$ through a one-to-one transformation, we update $\theta$ in an MCMC step. As the transformed variable $\theta$ can take any value on the real line, we use a $\mathrm{N}(\mu, \sigma_0^2)$ distribution as the proposal for the random walk Metropolis-Hastings algorithm. The proposal distribution has the desirable property of symmetry. Let $h(x;\mu,\sigma_0^2)$ denote the density of $\mathrm{N}(\mu,\sigma_0^2)$ distribution at the point $x$. We choose $\sigma_0^2 = \frac{1}{4}$. Let $\theta^{(l-1)}$ be the sample drawn from $f(\theta|\cdot)$ at the $(l-1)^{\text{th}}$ step of MCMC. The $l^{\text{th}}$ step of the random walk Metropolis-Hastings algorithm is as follows:

- Generate a random sample $\theta^{(l)}$ from the $\mathrm{N}(\theta^{(l-1)}, \sigma_0^2)$ distribution.
- Draw a random number $U \sim \text{Uniform}(0,1)$.
- Set

$$v^{(l)} = \begin{cases} \theta^{(l)}, & \text{if } 0 < U < r(\theta^{(l-1)}, \theta^{(l)}) \\ \theta^{(l-1)}, & \text{if } U > r(\theta^{(l-1)}, \theta^{(l)}) \end{cases}$$

Then $v^{(l)}$ is the generated sample from $f(\theta|\cdot)$ at the $l^{\text{th}}$ step. $r(\theta^{(l-1)}, \theta^{(l)})$ is the Metropolis-Hastings acceptance ratio with the following analytic form

$$\begin{aligned} r(\theta^{(l-1)}, \theta^{(l)}) &= \min\left\{1, \frac{f(\theta^{(l)})h(\theta^{(l-1)}; \theta^{(l)}, \sigma_0^2)}{f(\theta^{(l-1)})h(\theta^{(l)}; \theta^{(l-1)}, \sigma_0^2)}\right\} \\ &= \min\left\{1, \frac{f(\theta^{(l)})}{f(\theta^{(l-1)})}\right\}; \end{aligned}$$

since $h(\theta^{(l-1)}; \theta^{(l)}, \sigma_0^2) = h(\theta^{(l)}; \theta^{(l-1)}, \sigma_0^2)$.

### 6 MCMC algorithm for glossTWAS

We describe the Metropolis within Gibbs sampler used for glossTWAS. We implement the algorithm by integrating out the parameter $q$ from the MCMC procedure in both single-ancestry and multi-ancestry TWAS settings. This integration does not alter the previously derived full conditional distributions. However, it modifies the mixing probabilities associated with the full conditional posterior distributions of $\beta, \sigma_d^2$, and $\sigma_\epsilon^2$. In single-ancestry TWAS, the mixing weight corresponding to $\delta_{\{0\}}(\beta)$ in the full conditional distribution of $\beta$ is $\widetilde{w} = \frac{1-q}{1-q+\frac{q\widetilde{\sigma}}{\sigma_d \sigma_\epsilon}\exp\left(\frac{\widetilde{\mu}^2}{2\widetilde{\sigma}^2}\right)}$ (Equation 5). When $q$ is integrated out of MCMC, this weight revised to $\widetilde{w} = \frac{1-E(q)}{1-E(q)+\frac{E(q)\widetilde{\sigma}}{\sigma_d \sigma_\epsilon}\exp\left(\frac{\widetilde{\mu}^2}{2\widetilde{\sigma}^2}\right)}$, where $E(q) = \frac{b_1}{b_1+b_2}$. The similar modifications apply to the mixing probabilities in the full conditional distribution of $\sigma_d^2$ and $\sigma_\epsilon^2$, where the parameter $q$ is replaced by $E(q)$. These modifications are outlined in the algorithmic steps described below.

**Algorithm S1** Metropolis within Gibbs sampler for single-ancestry TWAS

1: Start

2: Choose initial values for parameters $\sigma_e^2$, $\boldsymbol{\lambda}$, $\tau$, $\beta$, $\sigma_d^2$, $\sigma_\epsilon^2$. Set $E(q) = \frac{b_1}{b_1+b_2}$.

3: loop:

4: Update $\boldsymbol{\alpha}$ by drawing a random vector from $\mathrm{N}_m\left(\Sigma^{-1}\left(\frac{1}{\sigma_e^2}G_{\mathrm{ref}}^{\mathrm{T}}\mathbf{x}_{\mathrm{ref}} + \frac{\beta}{\sigma_\epsilon^2}G^{\mathrm{T}}\mathbf{y}\right), \Sigma^{-1}\right)$ distribution where $\Sigma = \frac{1}{\sigma_e^2}G_{\mathrm{ref}}^{\mathrm{T}}G_{\mathrm{ref}} + \frac{\beta^2}{\sigma_\epsilon^2}G^{\mathrm{T}}G + \frac{1}{\sigma_e^2\tau^2}D_\lambda^{-1}$.

5: The conditional posterior distribution of $\lambda_j$ is given by $f(\lambda_j|\cdot) \propto \frac{1}{\lambda_j(1+\lambda_j^2)} \exp\left[-\frac{\alpha_j^2}{2\sigma_e^2\tau^2\lambda_j^2}\right], \quad j = 1(1)m$. We update each $\lambda_j$ by the Metropolis-Hastings algorithm outlined in the section 3.1.

6: The full conditional distribution of $\tau$ is given by $f(\tau|\cdot) \propto \frac{1}{\tau^m(1+\tau^2)} \exp\left[-\frac{\boldsymbol{\alpha}^{\mathrm{T}}D_\lambda^{-1}\boldsymbol{\alpha}}{2\sigma_e^2\tau^2}\right]$. We update $\tau$ by the Metropolis-Hastings algorithm discussed in the section 3.2.

7: We update $\sigma_e^2$ in the Gibbs sampler by drawing a random number from $\mathrm{IG}\left(e_1 + \frac{n}{2} + \frac{m}{2},\ e_2 + \frac{\boldsymbol{\alpha}^{\mathrm{T}}D_\lambda^{-1}\boldsymbol{\alpha}}{2\tau^2} + \frac{1}{2}(\mathbf{x}_{\mathrm{ref}} - G_{\mathrm{ref}}\boldsymbol{\alpha})^{\mathrm{T}}(\mathbf{x}_{\mathrm{ref}} - G_{\mathrm{ref}}\boldsymbol{\alpha})\right)$.

8: To update $\beta$, we first compute the probability corresponding to the point mass function $\delta_{\{0\}}(\beta)$ in the full conditional distribution of $\beta$ as $\widetilde{w} = \frac{1-E(q)}{1-E(q)+\frac{E(q)\widetilde{\sigma}}{\sigma_d\sigma_\epsilon}\exp\left(\frac{\widetilde{\mu}^2}{2\widetilde{\sigma}^2}\right)}$, where $\widetilde{\mu} = \frac{\boldsymbol{\alpha}^{\mathrm{T}}G^{\mathrm{T}}\mathbf{y}}{\boldsymbol{\alpha}^{\mathrm{T}}G^{\mathrm{T}}G\boldsymbol{\alpha}+\frac{1}{\sigma_d^2}}$ and $\widetilde{\sigma}^2 = \frac{\sigma_\epsilon^2}{\boldsymbol{\alpha}^{\mathrm{T}}G^{\mathrm{T}}G\boldsymbol{\alpha}+\frac{1}{\sigma_d^2}}$. Generate $U \sim$ Uniform$(0,1)$ distribution. **if** $U < \widetilde{w}$, **then** assign $\beta = 0$; **else** simulate $\beta$ from $\mathrm{N}(\widetilde{\mu}, \widetilde{\sigma}^2)$ distribution.

9: To update $\sigma_d^2$ first compute two quantities $\widetilde{w}_1 = \frac{(1-E(q))\delta_{\{0\}}(\beta)\Gamma(d_1)}{d_2^{d_1}}$ and $\widetilde{w}_2 = \frac{E(q)\Gamma(d_1+\frac{1}{2})}{\sqrt{2\pi}\sigma_\epsilon\left(d_2+\frac{\beta^2}{2\sigma_\epsilon^2}\right)^{d_1+\frac{1}{2}}}$, where $\Gamma(\cdot)$ denotes the gamma function. Generate $U \sim$ Uniform$(0,1)$ distribution. **if** $U < \frac{\widetilde{w}_1}{\widetilde{w}_1+\widetilde{w}_2}$, **then** draw a sample from $\mathrm{IG}(d_1, d_2)$ distribution; **else** draw from $\mathrm{IG}\left(d_1 + \frac{1}{2},\ d_2 + \frac{\beta^2}{2\sigma_\epsilon^2}\right)$ distribution.

Continuation of Algorithm S1

10: The full conditional distribution of $\sigma_\epsilon^2$ is a mixture of two inverse gamma distributions and the weights corresponding to two components are $\widetilde{w}_1 = \frac{(1-E(q))\delta_{\{0\}}(\beta)\Gamma(\frac{N}{2}+\epsilon_1)}{\left(\epsilon_2+\frac{1}{2}(\mathbf{y}-\beta G\boldsymbol{\alpha})^{\mathrm{T}}(\mathbf{y}-\beta G\boldsymbol{\alpha})\right)^{\frac{N}{2}+\epsilon_1}}$ and $\widetilde{w}_2 = \frac{E(q)\Gamma(\frac{N+1}{2}+\epsilon_1)}{\sqrt{2\pi}\sigma_d\left(\epsilon_2+\frac{\beta^2}{2\sigma_d^2}+\frac{1}{2}(\mathbf{y}-\beta G\boldsymbol{\alpha})^{\mathrm{T}}(\mathbf{y}-\beta G\boldsymbol{\alpha})\right)^{\frac{N+1}{2}+\epsilon_1}}$ respectively. To update $\sigma_\epsilon^2$, first simulate $U \sim$ Uniform$(0,1)$ distribution. **if** $U < \frac{\widetilde{w}_1}{\widetilde{w}_1+\widetilde{w}_2}$, **then** simulate from IG $\left(\frac{N}{2}+\epsilon_1,\ \epsilon_2+\frac{1}{2}(\mathbf{y}-\beta G\boldsymbol{\alpha})^{\mathrm{T}}(\mathbf{y}-\beta G\boldsymbol{\alpha})\right)$ distribution; **else** generate a random sample from IG $\left(\frac{N+1}{2}+\epsilon_1,\ \epsilon_2+\frac{\beta^2}{2\sigma_d^2}+\frac{1}{2}(\mathbf{y}-\beta G\boldsymbol{\alpha})^{\mathrm{T}}(\mathbf{y}-\beta G\boldsymbol{\alpha})\right)$ distribution.

11: Repeat the steps (4) − (11) until all MCMC iterations are finished.

**Algorithm S2** Metropolis-within Gibbs sampler for multi-ancestry TWAS

1: Start

2: Choose initial values for the parameters $\boldsymbol{\lambda}, \tau, \beta, w, \sigma_d^2, \sigma_\epsilon^2, \sigma_{e_1}^2$ and $\sigma_{e_2}^2$. Set $E(q) = \frac{b_1}{b_1+b_2}$.

3: loop:

4: Draw $\boldsymbol{\alpha}_k$ from $\mathrm{N}_m\left(\Sigma_k^{-1}\left(\frac{1}{\sigma_{e_k}^2} G_{k,\mathrm{ref}}^{\mathrm{T}} \mathbf{x}_{k,\mathrm{ref}} + \frac{\beta w_k}{\sigma_\epsilon^2} G_k^{\mathrm{T}} \mathbf{y}_k\right), \Sigma_k^{-1}\right)$ for each $k = 1, 2$, where $\Sigma_k = \frac{1}{\sigma_{e_k}^2} G_{k,\mathrm{ref}}^{\mathrm{T}} G_{k,\mathrm{ref}} + \frac{\beta^2 w_k^2}{\sigma_\epsilon^2} G_k^{\mathrm{T}} G_k + \frac{1}{\sigma_{e_k}^2 \tau^2} D_\lambda^{-1}$. $w_k$ equals to $w$ for $k = 1$ and $(1-w)$ for $k = 2$.

5: For each $j = 1(1)m$, the full conditional distribution of $\lambda_j$ is given as follows: $f(\lambda_j|\cdot) \propto \frac{1}{\lambda_j^2(1+\lambda_j^2)} \exp\left[-\frac{1}{2\tau^2\lambda_j^2} \sum_{k=1}^{2} \frac{\alpha_{kj}^2}{\sigma_{e_k}^2}\right]$. We update each $\lambda_j$ by the Metropolis-Hastings algorithm as discussed in section 5.1.

6: The conditional posterior distribution of $\tau$ is given by $f(\tau|\cdot) \propto \frac{1}{\tau^{2m}(1+\tau^2)} \exp\left[-\frac{1}{2\tau^2} \sum_{k=1}^{2} \frac{\boldsymbol{\alpha}_k^{\mathrm{T}} D_\lambda^{-1} \boldsymbol{\alpha}_k}{\sigma_{e_k}^2}\right]$. We update $\tau$ by the Metropolis-Hastings algorithm outlined in the section 5.2.

7: For each $k = 1, 2$, we update $\sigma_{e_k}^2$ by simulating a random sample from $\mathrm{IG}\left(e_1 + \frac{n_k}{2} + \frac{m}{2},\ e_2 + \frac{\boldsymbol{\alpha}_k^{\mathrm{T}} D_\lambda^{-1} \boldsymbol{\alpha}_k}{2\tau^2} + \frac{1}{2}(\mathbf{x}_{k,\mathrm{ref}} - G_{k,\mathrm{ref}}\boldsymbol{\alpha}_k)^{\mathrm{T}}(\mathbf{x}_{k,\mathrm{ref}} - G_{k,\mathrm{ref}}\boldsymbol{\alpha}_k)\right)$.

8: The full conditional posterior distribution of $w$ is given as follows: $f(w|\cdot) \propto \exp\left[-\frac{1}{2\sigma_\epsilon^2}\left\{\beta^2\left(w^2 \boldsymbol{\alpha}_1^{\mathrm{T}} G_1^{\mathrm{T}} G_1 \boldsymbol{\alpha}_1 + (1-w)^2 \boldsymbol{\alpha}_2^{\mathrm{T}} G_2^{\mathrm{T}} G_2 \boldsymbol{\alpha}_2\right) - 2\beta\left(w \boldsymbol{\alpha}_1^{\mathrm{T}} G_1^{\mathrm{T}} \mathbf{y}_1 + (1-w) \boldsymbol{\alpha}_2^{\mathrm{T}} G_2^{\mathrm{T}} \mathbf{y}_2\right)\right\}\right]$. We update $w$ by random walk Metropolis-Hastings algorithm outlined in section 5.3.

9: The posterior probability that $\beta$ takes value 0 in the full conditional distribution is $\widetilde{w} = \frac{1-E(q)}{1-E(q)+\frac{E(q)\widetilde{\sigma}}{\sigma_d \sigma_\epsilon} \exp\left(\frac{\widetilde{\mu}^2}{2\widetilde{\sigma}^2}\right)}$, where $\widetilde{\mu} = \frac{\sum_{k=1}^{2} w_k \boldsymbol{\alpha}_k^{\mathrm{T}} G_k^{\mathrm{T}} \mathbf{y}_k}{\sum_{k=1}^{2} w_k^2 \boldsymbol{\alpha}_k^{\mathrm{T}} G_k^{\mathrm{T}} G_k \boldsymbol{\alpha}_k + \frac{1}{\sigma_d^2}}$ and $\widetilde{\sigma}^2 = \frac{\sigma_\epsilon^2}{\sum_{k=1}^{2} w_k^2 \boldsymbol{\alpha}_k^{\mathrm{T}} G_k^{\mathrm{T}} G_k \boldsymbol{\alpha}_k + \frac{1}{\sigma_d^2}}$. We update $\beta$ as follows: first simulate $U \sim \mathrm{Uniform}(0,1)$ distribution. **if** $U < \widetilde{w}$, **then** assign $\beta = 0$; **else** draw a sample from $\mathrm{N}(\widetilde{\mu}, \widetilde{\sigma}^2)$ distribution.

Continuation of Algorithm S2

10: To update $\sigma_d^2$, first compute two quantities $\widetilde{w}_1 = \frac{(1-E(q))\delta_{\{0\}}(\beta)\Gamma(d_1)}{d_2^{d_1}}$ and $\widetilde{w}_2 = \frac{E(q)\Gamma(d_1+\frac{1}{2})}{\sqrt{2\pi}\sigma_\epsilon\left(d_2+\frac{\beta^2}{2\sigma_\epsilon^2}\right)^{d_1+\frac{1}{2}}}$, where $\Gamma(\cdot)$ denotes the gamma function. Generate $U \sim$ Uniform$(0,1)$ distribution. **if** $U < \frac{\widetilde{w}_1}{\widetilde{w}_1+\widetilde{w}_2}$, **then** draw a sample from IG$(d_1, d_2)$ distribution; **else** draw from IG $\left(d_1+\frac{1}{2},\ d_2+\frac{\beta^2}{2\sigma_\epsilon^2}\right)$ distribution.

11: The full conditional distribution of $\sigma_\epsilon^2$ can be expressed as a mixture of two inverse gamma distributions, and the weights corresponding to two components are $\widetilde{w}_1 = \frac{(1-E(q))\delta_{\{0\}}(\beta)\Gamma(\frac{N_c}{2}+\epsilon_1)}{\left(\epsilon_2+\frac{1}{2}\sum_{k=1}^2(\mathbf{y}_k-\beta w_k G_k\boldsymbol{\alpha}_k)^{\mathrm{T}}(\mathbf{y}_k-\beta w_k G_k\boldsymbol{\alpha}_k)\right)^{\frac{N_c}{2}+\epsilon_1}}$ and $\widetilde{w}_2 = \frac{E(q)\ \Gamma(\frac{N_c+1}{2}+\epsilon_1)}{\sqrt{2\pi}\sigma_d\left(\epsilon_2+\frac{\beta^2}{2\sigma_d^2}+\frac{1}{2}\sum_{k=1}^2(\mathbf{y}_k-\beta w_k G_k\boldsymbol{\alpha}_k)^{\mathrm{T}}(\mathbf{y}_k-\beta w_k G_k\boldsymbol{\alpha}_k)\right)^{\frac{N_c+1}{2}+\epsilon_1}}$, respectively, where $N_c = N_1 + N_2$. Generate U $\sim$ Uniform$(0,1)$. We update $\sigma_\epsilon^2$ as follows. **if** $U < \frac{\widetilde{w}_1}{\widetilde{w}_1+\widetilde{w}_2}$, **then** draw from IG $\left(\frac{N_c}{2}+\epsilon_1,\ \epsilon_2+\frac{1}{2}\sum_{k=1}^2(\mathbf{y}_k-\beta w_k G_k\boldsymbol{\alpha}_k)^{\mathrm{T}}(\mathbf{y}_k-\beta w_k G_k\boldsymbol{\alpha}_k)\right)$; **else** generate a sample from IG $\left(\frac{N_c+1}{2}+\epsilon_1,\ \epsilon_2+\frac{\beta^2}{2\sigma_d^2}+\frac{1}{2}\sum_{k=1}^2(\mathbf{y}_k-\beta w_k G_k\boldsymbol{\alpha}_k)^{\mathrm{T}}(\mathbf{y}_k-\beta w_k G_k\boldsymbol{\alpha}_k)\right)$.

12: Repeat the steps (4) – (11) until all MCMC iterations are finished.

### Supplemental Figures

Figure S1: Comparison of TWAS effect size estimation accuracy by glossTWAS, PMR-EGGER, TIGAR, and PrediXcan for single-ancestry TWAS. The GReX effect sizes are negative. Two measures are considered: (**A**) relative bias and (**B**) RMSE. The transcriptome and GWAS data sample sizes are $n = 500$ and $N = 10000$, respectively. The local heritability of gene expression is fixed at $h_e^2 = 10\%$.

**(A)**

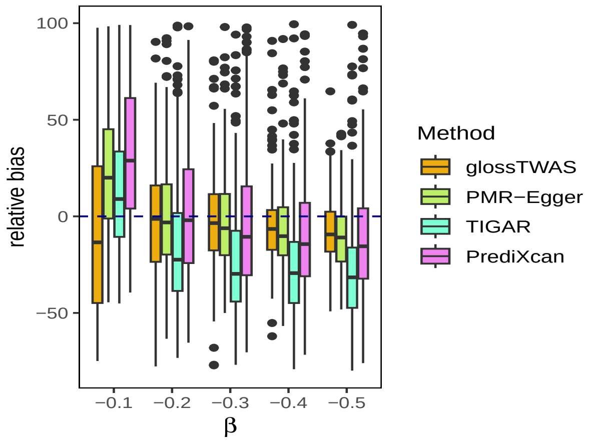

**(B)**

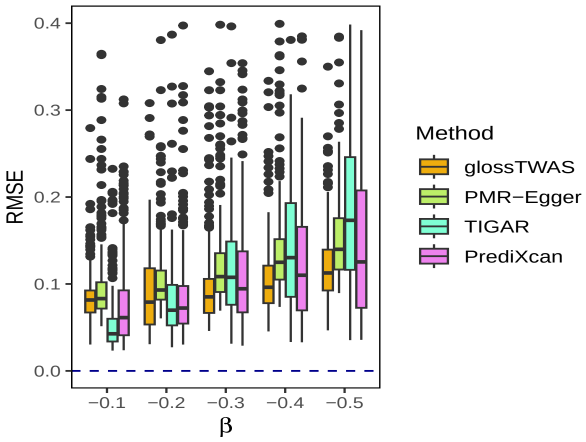

Figure S2: Comparison of estimation accuracy by glossTWAS, PMR-Egger, TIGAR, and PrediXcan in single-ancestry TWAS when the GReX has no effect on the trait, i.e., $\beta = 0$. The transcriptome data sample size ($n$) is varied from 500 to 1500. Two measures are considered: (**A**) bias and (**B**) RMSE. The GWAS sample size is $N = 10000$. The local heritability of gene expression is fixed at $h_e^2 = 10\%$.

**(A)**

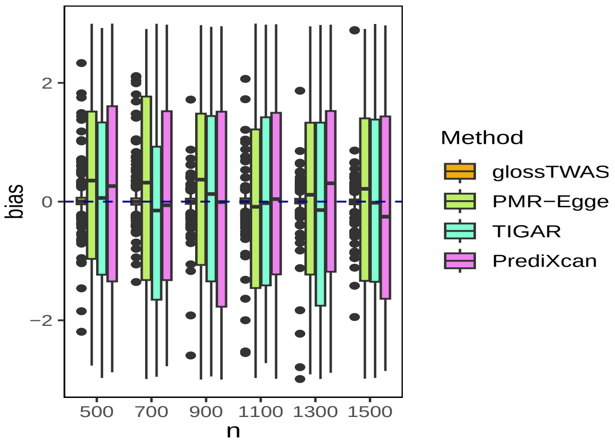

**(B)**

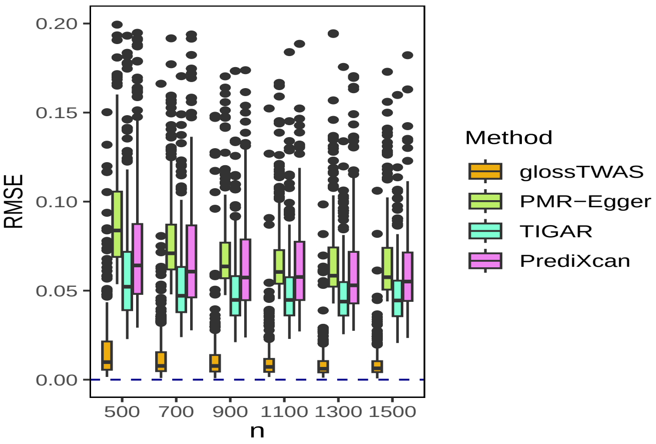

Figure S3: Comparison of estimation accuracy by glossTWAS and METRO in multi-ancestry TWAS when the GReX effect sizes corresponding to EAS ancestry ($\beta_1$) and EUR ancestry ($\beta_2$) are equal and negative. Here, half of the eQTLs are shared between the two ancestries, i.e, $f_0 = 50\%$. Two measures are considered: relative bias and RMSE for the EAS ancestry (Figure **A-B**) and the EUR ancestry (Figure **C-D**). The transcriptome data sample sizes are $n_1 = n_2 = 500$, and GWAS sample sizes are $N_1 = N_2 = 10000$ for the two ancestries. The local heritability of gene expression is fixed at $h_e^2 = 10\%$ in each ancestry.

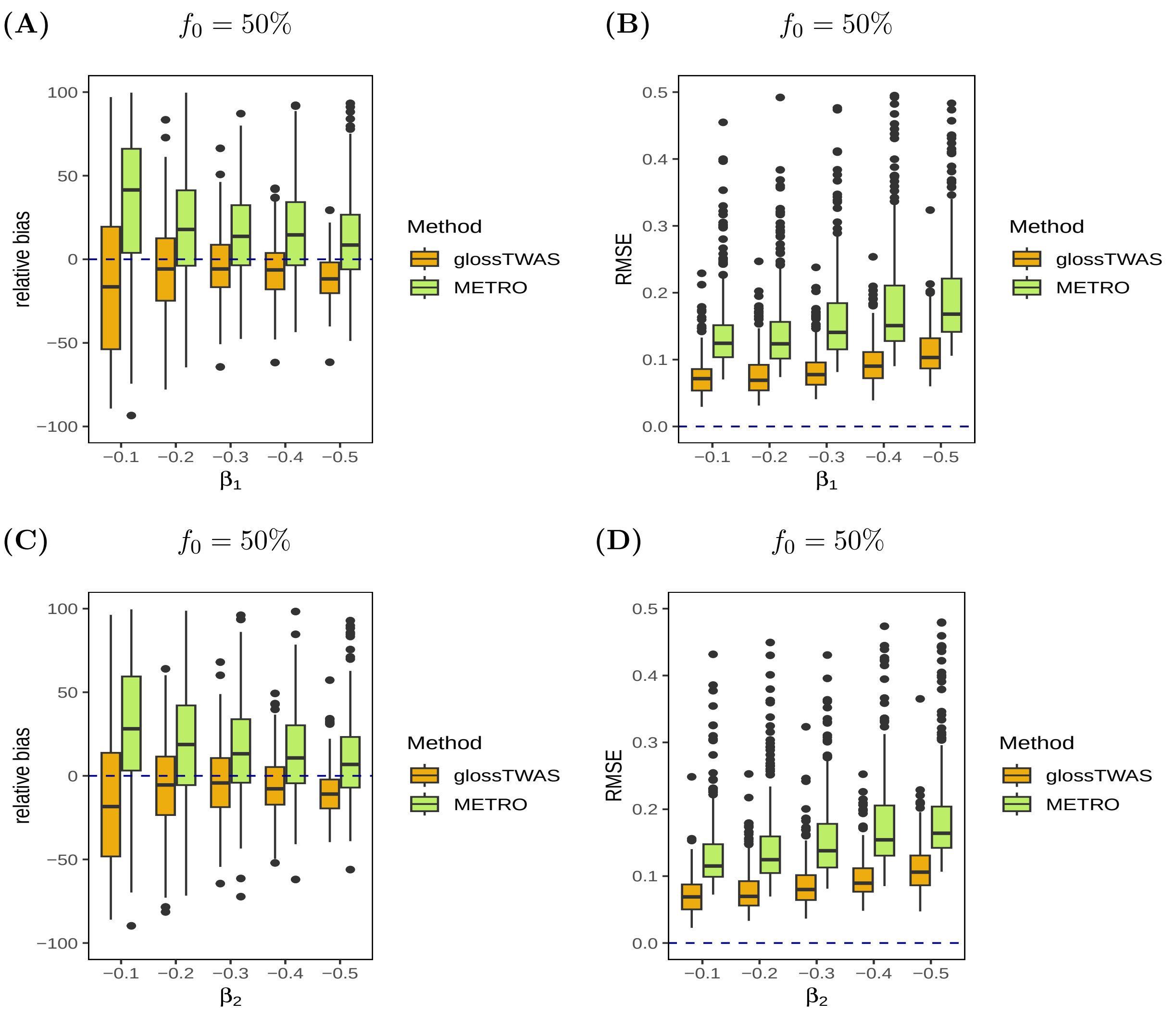

Figure S4: Comparison of estimation accuracy by glossTWAS and METRO in multi-ancestry TWAS when the GReX effect sizes corresponding to EAS ancestry ($\beta_1$) and EUR ancestry ($\beta_2$) are unequal and negative. Here, half of the eQTLs are shared between the two ancestries, i.e, $f_0 = 50\%$. Two measures, relative bias and RMSE, are considered for the following settings: $\beta_1 = -0.1, \beta_2 = -0.5$ (Figure **A-B**) and $\beta_1 = -0.1, \beta_2 = -0.2$ (Figure **C-D**). The transcriptome data sample sizes are $n_1 = n_2 = 500$, and GWAS sample sizes are $N_1 = N_2 = 10000$ for the two ancestries. The local heritability of gene expression is fixed at $h_e^2 = 10\%$ in each ancestry.

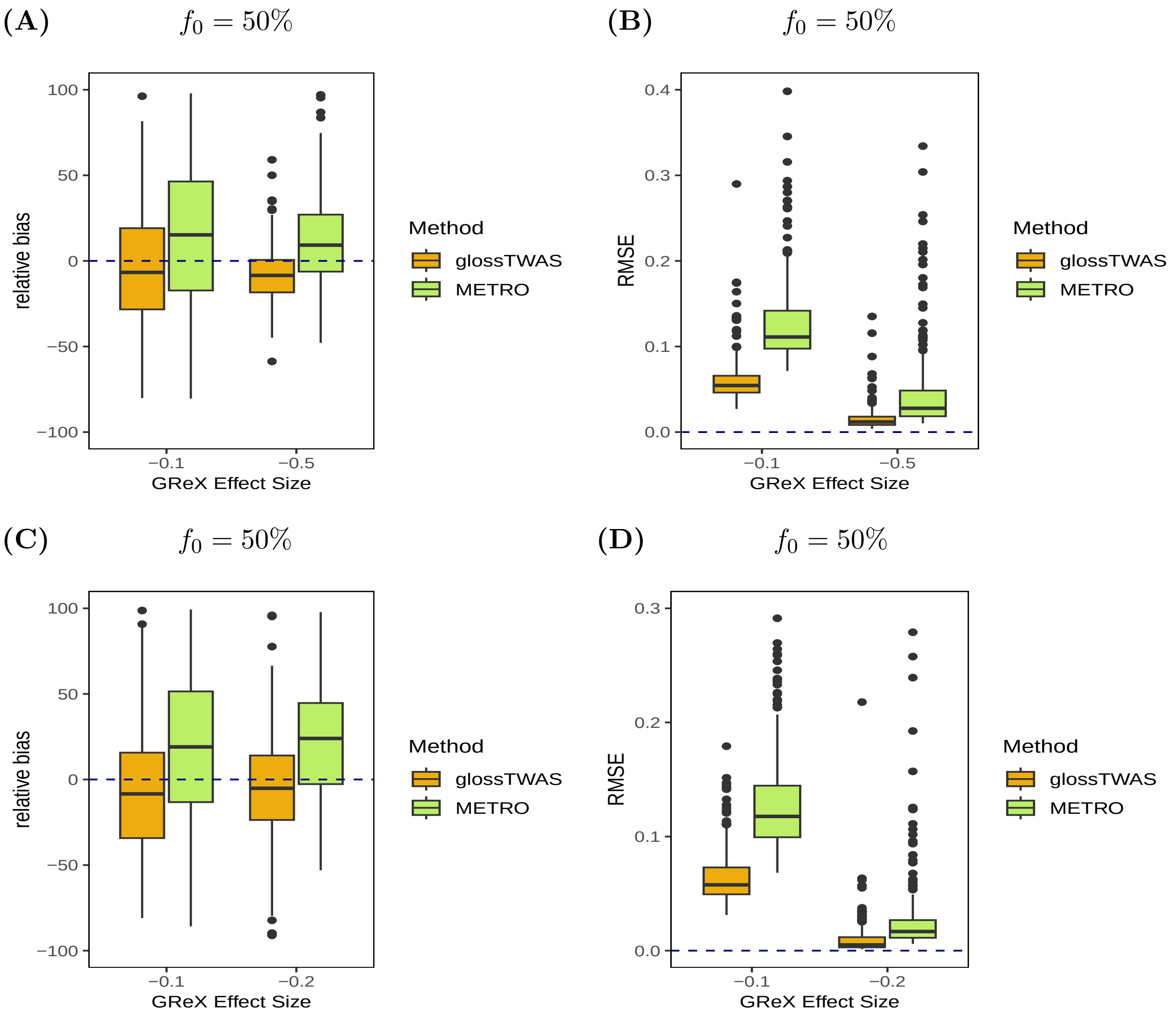

Figure S5: Comparison of estimation accuracy by glossTWAS and METRO in multi-ancestry TWAS when the GReX has no effect on the trait in both ancestries, i.e., $\beta_1 = 0$ and $\beta_2 = 0$. Here, half of the eQTLs are shared between the two ancestries, i.e, $f_0 = 50\%$. The transcriptome data sample size for EAS ancestry ($n_1$) is varied from 500 to 1500, keeping the sample size for the EUR ancestry ($n_2$) fixed at 500. Two measures are considered: bias and RMSE for the EAS ancestry (Figure **A**-**B**) and the EUR ancestry (Figure **C**-**D**). The GWAS sample sizes are $N_1 = N_2 = 10000$ for the two ancestries. The local heritability of gene expression is fixed at $h_e^2 = 10\%$ in each ancestry.

**(A)** $f_0 = 50\%$

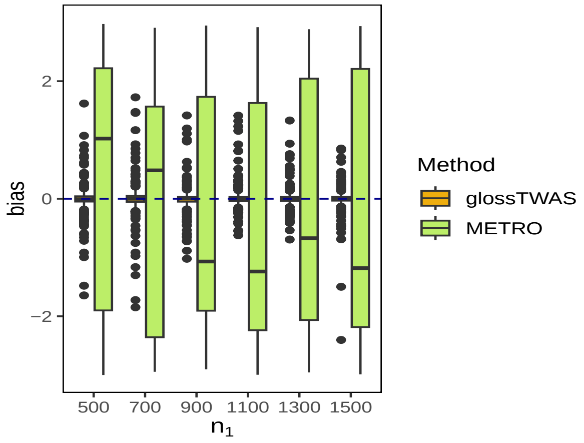

**(B)** $f_0 = 50\%$

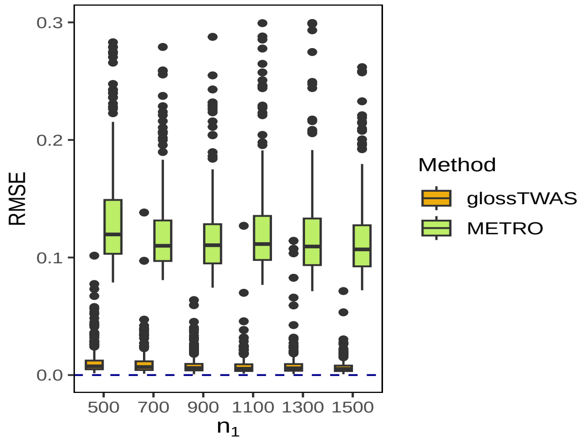

**(C)** $f_0 = 50\%$

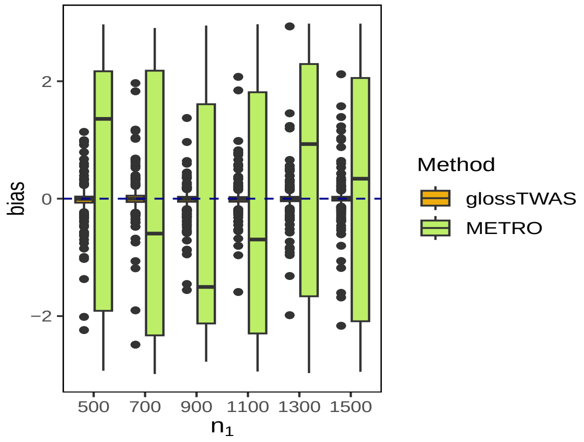

**(D)** $f_0 = 50\%$

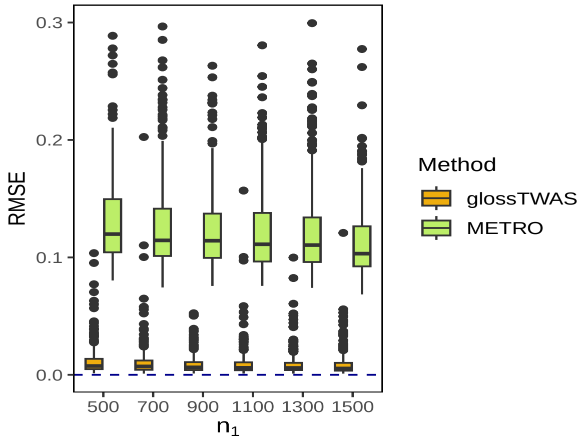

Figure S6: Comparison of estimation accuracy by glossTWAS and METRO in multi-ancestry TWAS when the GReX has no effect on the trait in both ancestries, i.e., $\beta_1 = 0$ and $\beta_2 = 0$. Here, eQTLs are not shared between the two ancestries, i.e, $f_0 = 0\%$. The transcriptome data sample size for EAS ancestry ($n_1$) is varied from 500 to 1500, keeping the sample size for the EUR ancestry ($n_2$) fixed at 500. Two measures are considered: bias and RMSE for the EAS ancestry (Figure **A-B**) and the EUR ancestry (Figure **C-D**). The GWAS sample sizes are $N_1 = N_2 = 10000$ for the two ancestries. The local heritability of gene expression is fixed at $h_e^2 = 10\%$ in each ancestry.

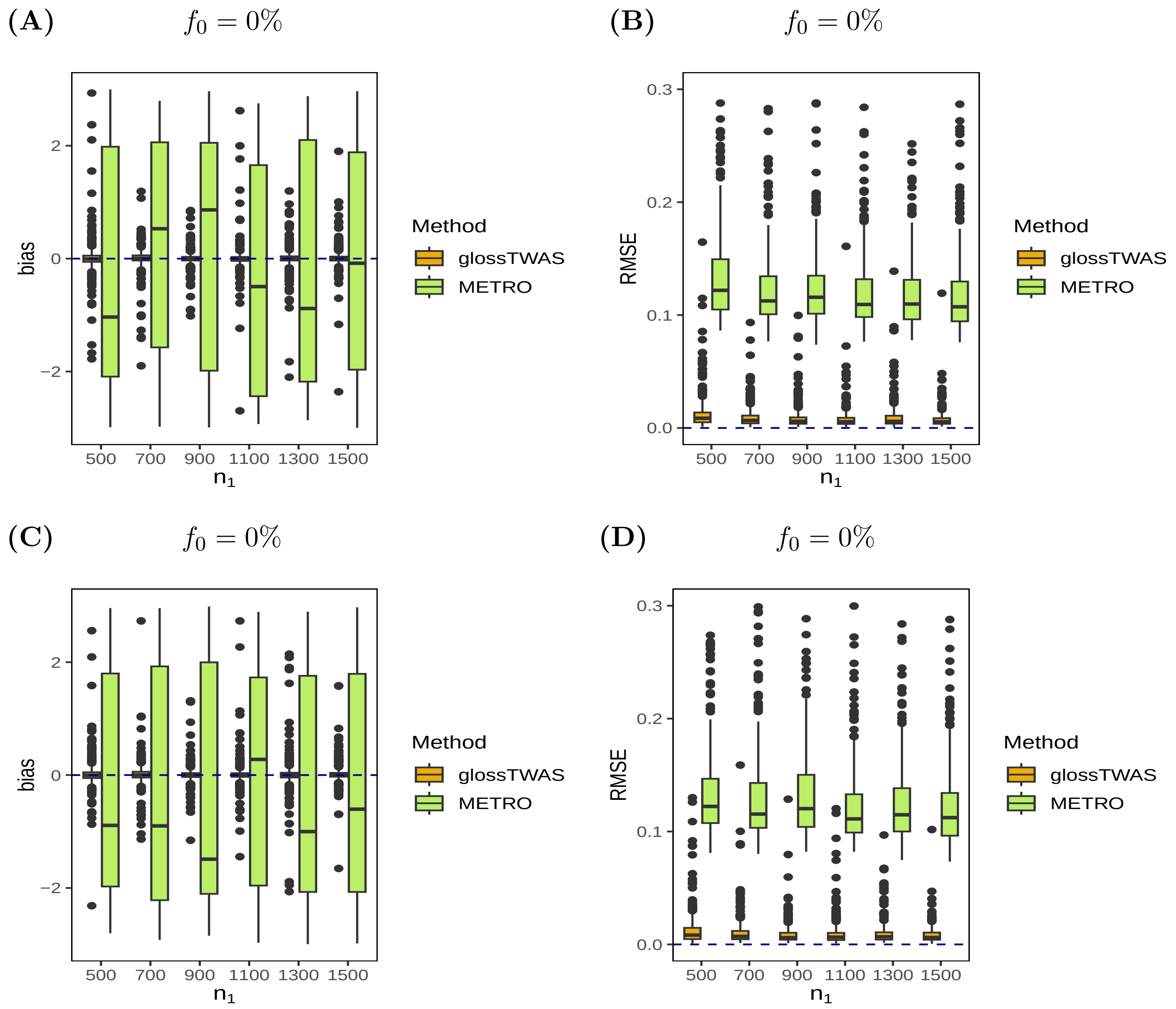

Figure S7: Comparison of estimation accuracy by glossTWAS and METRO in multi-ancestry TWAS when the GReX effect sizes corresponding to EAS ancestry ($\beta_1$) and EUR ancestry ($\beta_2$) are equal. Here, the eQTLs are not shared between the two ancestries, i.e, $f_0 = 0\%$. Two measures are considered: relative bias and RMSE for the EAS ancestry (Figure **A**-**B**) and the EUR ancestry (Figure **C**-**D**). The transcriptome data sample sizes are $n_1 = n_2 = 500$, and GWAS sample sizes are $N_1 = N_2 = 10000$ for the two ancestries. The local heritability of gene expression is fixed at $h_e^2 = 10\%$ in each ancestry.

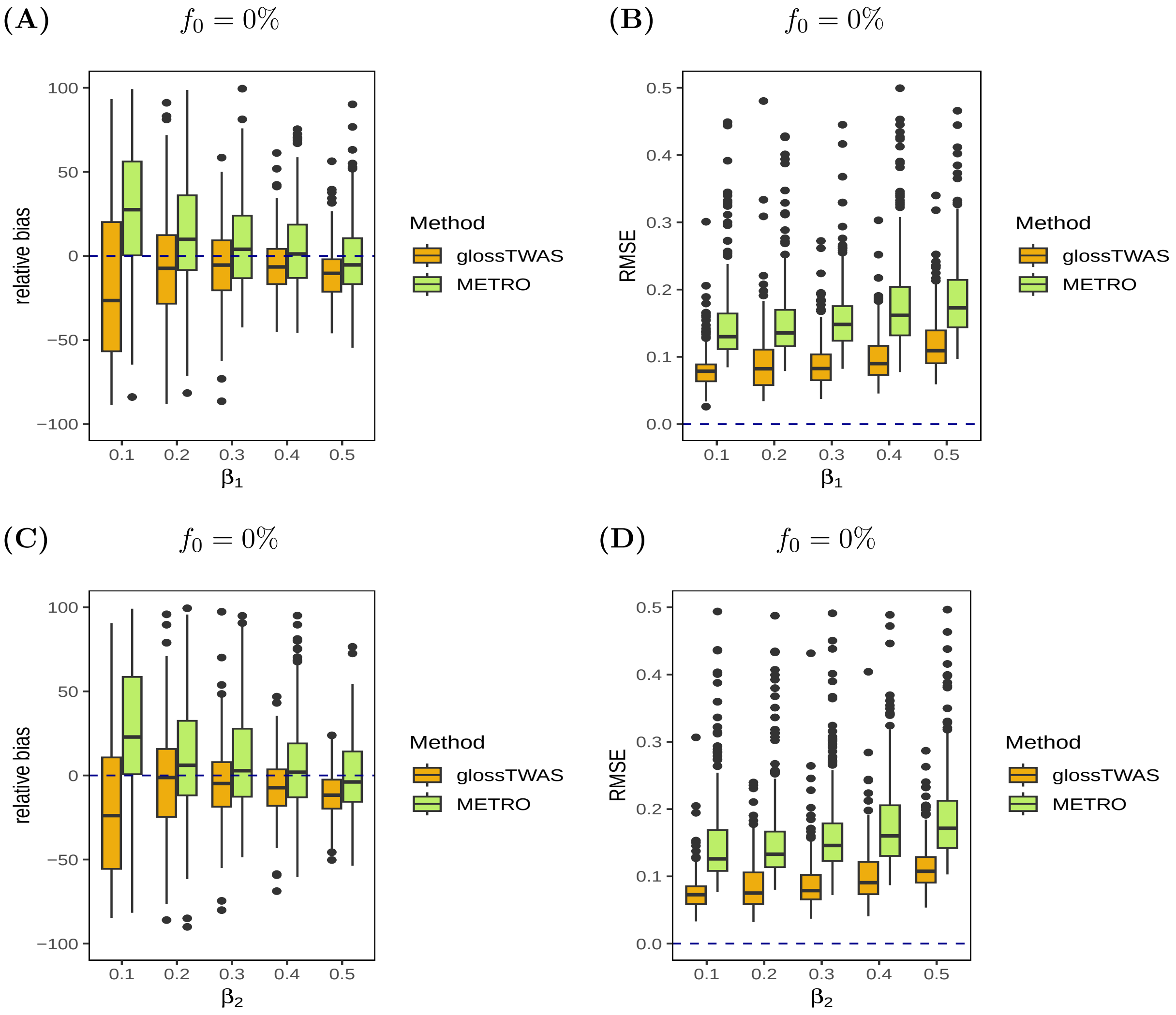

Figure S8: Comparison of estimation accuracy by glossTWAS and METRO in multi-ancestry TWAS when the GReX effect sizes corresponding to EAS ancestry ($\beta_1$) and EUR ancestry ($\beta_2$) are equal and negative. Here, the eQTLs are not shared between the two ancestries, i.e, $f_0 = 0\%$. Two measures are considered: relative bias and RMSE for the EAS ancestry (Figure **A-B**) and the EUR ancestry (Figure **C-D**). The transcriptome data sample sizes are $n_1 = n_2 = 500$, and GWAS sample sizes are $N_1 = N_2 = 10000$ for the two ancestries. The local heritability of gene expression is fixed at $h_e^2 = 10\%$ in each ancestry.

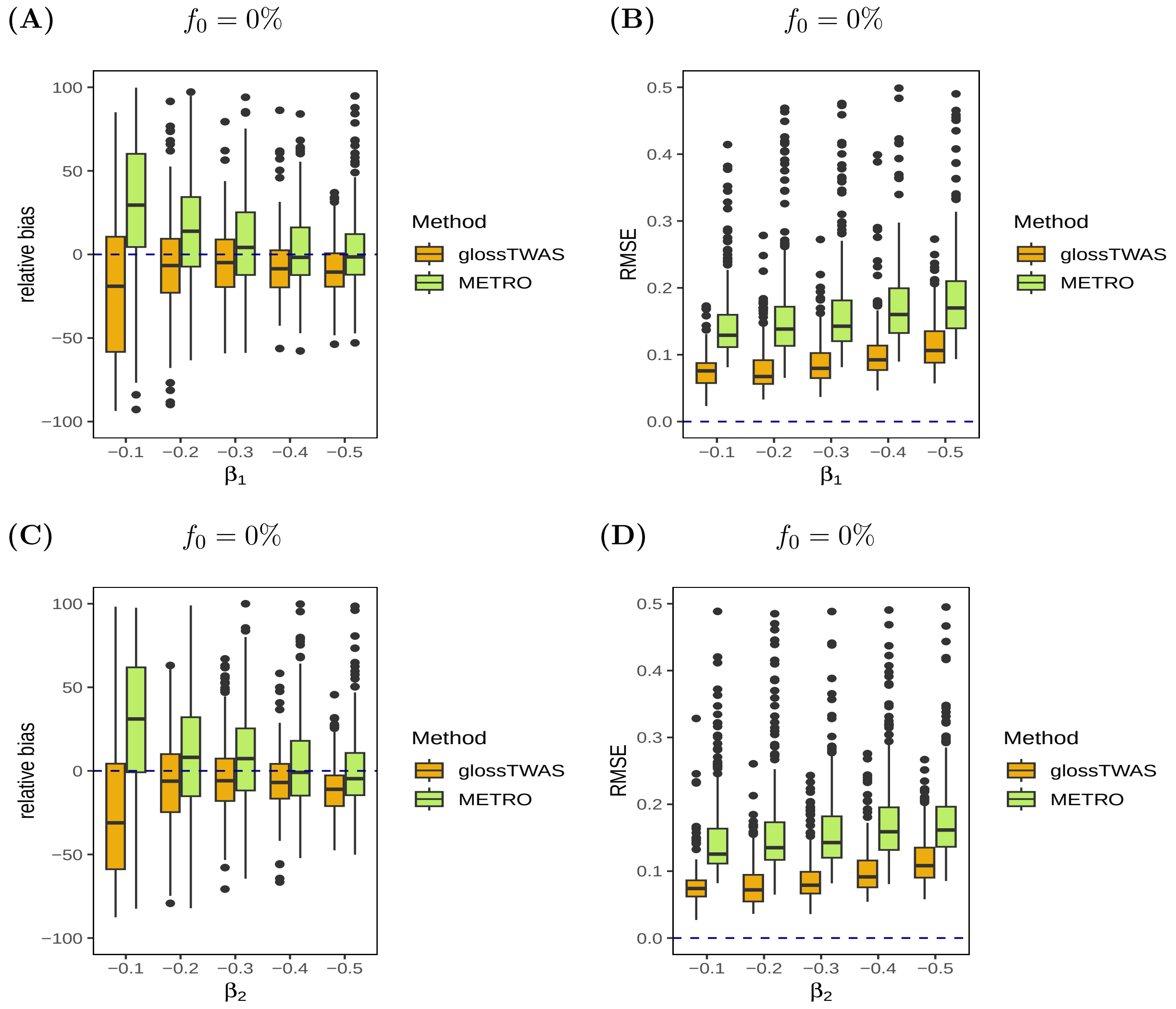

Figure S9: Comparison of estimation accuracy by glossTWAS and METRO in multi-ancestry TWAS when the GReX effect sizes corresponding to EAS ancestry ($\beta_1$) and EUR ancestry ($\beta_2$) are unequal. Here, the eQTLs are not shared between the two ancestries, i.e, $f_0 = 0\%$. Two measures, relative bias and RMSE, are considered for the following settings: $\beta_1 = 0.1, \beta_2 = 0.5$ (Figure **A-B**) and $\beta_1 = 0.1, \beta_2 = 0.2$ (Figure **C-D**). The transcriptome data sample sizes are $n_1 = n_2 = 500$, and GWAS sample sizes are $N_1 = N_2 = 10000$ for the two ancestries. The local heritability of gene expression is fixed at $h_e^2 = 10\%$ in each ancestry.

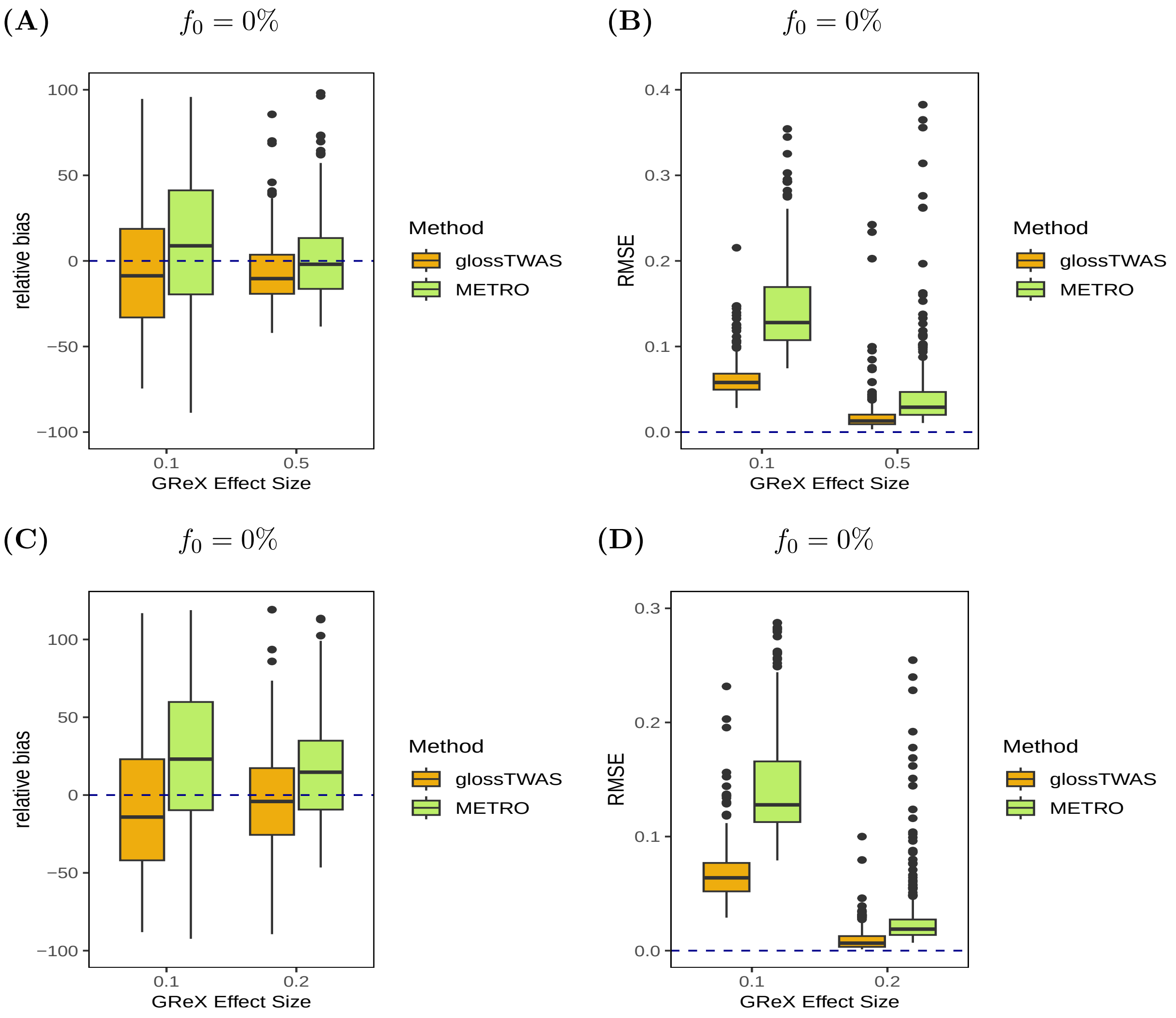

Figure S10: Comparison of estimation accuracy by glossTWAS and METRO in multi-ancestry TWAS when the GReX effect sizes corresponding to EAS ancestry ($\beta_1$) and EUR ancestry ($\beta_2$) are unequal and negative. Here, the eQTLs are not shared between the two ancestries, i.e, $f_0 = 0\%$. Two measures, relative bias and RMSE, are considered for the following settings: $\beta_1 = -0.1, \beta_2 = -0.5$ (Figure **A-B**) and $\beta_1 = -0.1, \beta_2 = -0.2$ (Figure **C-D**). The transcriptome data sample sizes are $n_1 = n_2 = 500$, and GWAS sample sizes are $N_1 = N_2 = 10000$ for the two ancestries. The local heritability of gene expression is fixed at $h_e^2 = 10\%$ in each ancestry.

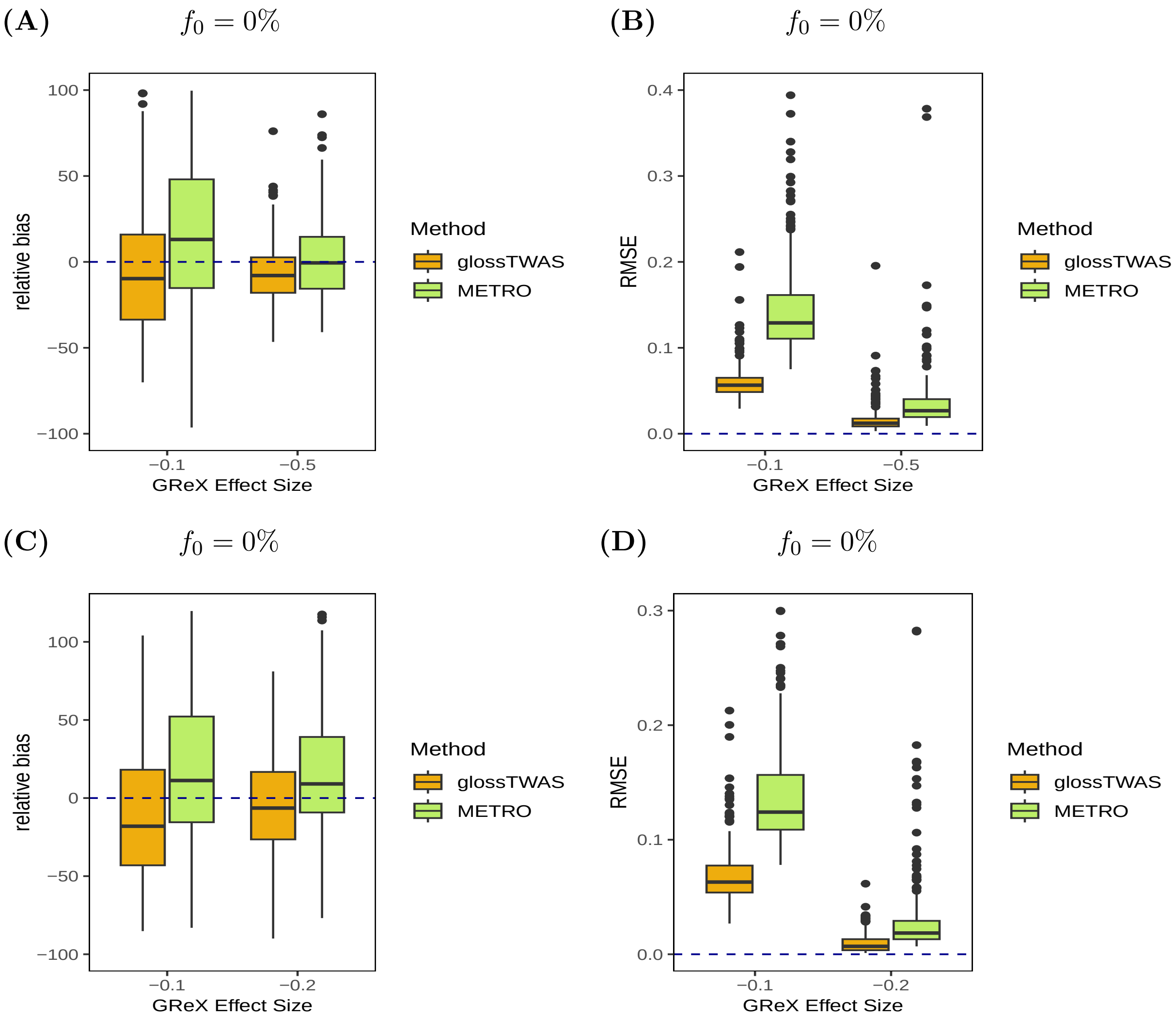

### Supplemental Tables

Table S1: Summaries related to realized FDR (rFDR) and true positive rate (TPR) for different levels of Bayesian FDR (BFDR) for glossTWAS in single-ancestry TWAS.

| BFDR | $n$ | $N$ | $h_e^2$ | $h_g^2$ | rFDR | TPR |
|---|---|---|---|---|---|---|
| 0.01 | 500 | 10000 | 10% | 10% | 0 | 0.1 |
| | | | 10% | 20% | 0 | 0.22 |
| | | | 20% | 10% | 0 | 0.26 |
| | | | 20% | 20% | 0 | 0.45 |
| | | | 30% | 30% | 0 | 0.61 |
| | | | 40% | 40% | 0 | 0.74 |
| 0.05 | 500 | 10000 | 10% | 10% | 0.01 | 0.14 |
| | | | 10% | 20% | 0.02 | 0.27 |
| | | | 20% | 10% | 0.01 | 0.32 |
| | | | 20% | 20% | 0.01 | 0.49 |
| | | | 30% | 30% | 0.01 | 0.65 |
| | | | 40% | 40% | 0.02 | 0.76 |
| 0.01 | 1000 | 10000 | 10% | 10% | 0 | 0.14 |
| | | | 10% | 20% | 0 | 0.28 |
| | | | 20% | 10% | 0 | 0.26 |
| | | | 20% | 20% | 0.01 | 0.41 |
| | | | 30% | 30% | 0 | 0.64 |
| | | | 40% | 40% | 0.01 | 0.75 |
| 0.05 | 1000 | 10000 | 10% | 10% | 0.03 | 0.18 |
| | | | 10% | 20% | 0.02 | 0.33 |
| | | | 20% | 10% | 0.02 | 0.32 |

Continuation of Table S1

| BFDR | $n$ | $N$ | $h_e^2$ | $h_g^2$ | rFDR | TPR |
|---|---|---|---|---|---|---|
| | | | 20% | 20% | 0.02 | 0.47 |
| | | | 30% | 30% | 0.01 | 0.68 |
| | | | 40% | 40% | 0.01 | 0.79 |
| 0.01 | 500 | 20000 | 10% | 10% | 0 | 0.28 |
| | | | 10% | 20% | 0 | 0.34 |
| | | | 20% | 10% | 0 | 0.39 |
| | | | 20% | 20% | 0 | 0.57 |
| | | | 30% | 30% | 0 | 0.68 |
| | | | 40% | 40% | 0 | 0.8 |
| 0.05 | 500 | 20000 | 10% | 10% | 0.01 | 0.34 |
| | | | 10% | 20% | 0 | 0.39 |
| | | | 20% | 10% | 0 | 0.45 |
| | | | 20% | 20% | 0.01 | 0.62 |
| | | | 30% | 30% | 0 | 0.72 |
| | | | 40% | 40% | 0 | 0.83 |

Note: The notations BFDR, FPR, TPR denote the Bayesian false discovery rate, false positive rate and true positive rate, respectively. rFDR is the realized value of the BFDR calculated as the number of false positives divided by the sum of false and true positives. The transcriptome data sample size ($n$), GWAS sample size ($N$), gene expression heritability ($h_e^2$) and trait heritability ($h_g^2$) are varied. The results are obtained based on 50 replications in a simulation scenario.

Table S2: Summaries related to false discoveries for PMR-EGGER, TIGAR and PrediXcan in single-ancestry TWAS.

| | | | | | PMR-Egger | | | TIGAR | | | PrediXcan | | |
|---|---|---|---|---|---|---|---|---|---|---|---|---|---|
| FDR | $n$ | $N$ | $h_e^2$ | $h_g^2$ | rFDR | FPR | TPR | rFDR | FPR | TPR | rFDR | FPR | TPR |
| 0.01 | 500 | 10000 | 10% | 10% | 0 | 0 | 0.1 | 0 | 0 | 0.09 | 0.01 | $1.1 \times 10^{-4}$ | 0.1 |
| | | | 10% | 20% | 0 | $1.1 \times 10^{-4}$ | 0.24 | 0.01 | $1.1 \times 10^{-4}$ | 0.21 | 0.01 | $1.1 \times 10^{-4}$ | 0.2 |
| | | | 20% | 10% | 0.02 | $3.2 \times 10^{-4}$ | 0.23 | 0.01 | $2.1 \times 10^{-4}$ | 0.26 | 0.02 | $4.2 \times 10^{-4}$ | 0.29 |
| | | | 20% | 20% | 0.01 | $3.2 \times 10^{-4}$ | 0.42 | 0.01 | $3.2 \times 10^{-4}$ | 0.42 | 0.01 | $3.2 \times 10^{-4}$ | 0.45 |
| 0.05 | 500 | 10000 | 10% | 10% | 0.05 | $5.3 \times 10^{-4}$ | 0.13 | 0.05 | $1.1 \times 10^{-3}$ | 0.15 | 0.04 | $7.4 \times 10^{-4}$ | 0.15 |
| | | | 10% | 20% | 0.05 | $1.4 \times 10^{-3}$ | 0.3 | 0.07 | $1.1 \times 10^{-3}$ | 0.25 | 0.05 | $1.2 \times 10^{-3}$ | 0.26 |
| | | | 20% | 10% | 0.04 | $1.1 \times 10^{-3}$ | 0.28 | 0.08 | $1.8 \times 10^{-3}$ | 0.31 | 0.08 | $2.2 \times 10^{-3}$ | 0.37 |
| | | | 20% | 20% | 0.06 | $2.2 \times 10^{-3}$ | 0.5 | 0.04 | $1.3 \times 10^{-3}$ | 0.48 | 0.06 | $2.2 \times 10^{-3}$ | 0.51 |
| 0.1 | 500 | 10000 | 10% | 10% | 0.05 | $9.5 \times 10^{-4}$ | 0.15 | 0.07 | $1.9 \times 10^{-3}$ | 0.17 | 0.1 | $1.5 \times 10^{-3}$ | 0.18 |
| | | | 10% | 20% | 0.09 | $2.8 \times 10^{-3}$ | 0.33 | 0.14 | $2.6 \times 10^{-3}$ | 0.28 | 0.12 | $3.5 \times 10^{-3}$ | 0.3 |
| | | | 20% | 10% | 0.07 | $2.2 \times 10^{-3}$ | 0.31 | 0.12 | $3.7 \times 10^{-3}$ | 0.34 | 0.12 | $3.8 \times 10^{-3}$ | 0.4 |
| | | | 20% | 20% | 0.1 | $3.7 \times 10^{-3}$ | 0.53 | 0.1 | $3.9 \times 10^{-3}$ | 0.5 | 0.12 | $4.6 \times 10^{-3}$ | 0.54 |
| 0.2 | 500 | 10000 | 10% | 10% | 0.11 | $2.4 \times 10^{-3}$ | 0.2 | 0.17 | $4.7 \times 10^{-3}$ | 0.21 | 0.17 | $4.9 \times 10^{-3}$ | 0.24 |
| | | | 10% | 20% | 0.16 | $6.2 \times 10^{-3}$ | 0.38 | 0.22 | $5.7 \times 10^{-3}$ | 0.31 | 0.22 | $7.1 \times 10^{-3}$ | 0.33 |
| | | | 20% | 10% | 0.13 | $4.5 \times 10^{-3}$ | 0.34 | 0.26 | $8.9 \times 10^{-3}$ | 0.38 | 0.23 | $8.7 \times 10^{-3}$ | 0.43 |
| | | | 20% | 20% | 0.17 | $7.7 \times 10^{-3}$ | 0.56 | 0.21 | $9.9 \times 10^{-3}$ | 0.54 | 0.21 | 0.01 | 0.57 |
| 0.01 | 1000 | 10000 | 10% | 10% | 0 | 0 | 0.1 | 0.02 | $2.1 \times 10^{-4}$ | 0.11 | 0.01 | $2.1 \times 10^{-4}$ | 0.14 |
| | | | 10% | 20% | 0.01 | $2.1 \times 10^{-4}$ | 0.26 | 0 | 0 | 0.23 | 0 | $1.1 \times 10^{-4}$ | 0.31 |
| | | | 20% | 10% | 0.01 | $1.1 \times 10^{-4}$ | 0.27 | 0.02 | $3.2 \times 10^{-4}$ | 0.27 | 0.02 | $6.3 \times 10^{-4}$ | 0.29 |
| | | | 20% | 20% | 0.01 | $1.1 \times 10^{-4}$ | 0.44 | 0.02 | $3.2 \times 10^{-4}$ | 0.4 | 0.03 | $2.1 \times 10^{-4}$ | 0.46 |
| 0.05 | 1000 | 10000 | 10% | 10% | 0.02 | $3.2 \times 10^{-4}$ | 0.14 | 0.06 | 0.01 | 0.17 | 0.04 | $7.4 \times 10^{-4}$ | 0.19 |

| | | | | | PMR-Egger | | | TIGAR | | | PrediXcan | | |
|---|---|---|---|---|---|---|---|---|---|---|---|---|---|
| FDR | $n$ | $N$ | $h_e^2$ | $h_g^2$ | rFDR | FPR | TPR | rFDR | FPR | TPR | rFDR | FPR | TPR |
| | | | 10% | 20% | 0.03 | $9.5 \times 10^{-4}$ | 0.33 | 0.04 | $8.4 \times 10^{-4}$ | 0.28 | 0.04 | $1.1 \times 10^{-3}$ | 0.35 |
| | | | 20% | 10% | 0.05 | $1.3 \times 10^{-3}$ | 0.35 | 0.08 | $2 \times 10^{-3}$ | 0.34 | 0.06 | $1.8 \times 10^{-3}$ | 0.37 |
| | | | 20% | 20% | 0.05 | $1.9 \times 10^{-3}$ | 0.5 | 0.06 | $1.7 \times 10^{-3}$ | 0.47 | 0.05 | $1.3 \times 10^{-3}$ | 0.51 |
| 0.1 | 1000 | 10000 | 10% | 10% | 0.06 | $9.5 \times 10^{-4}$ | 0.18 | 0.08 | 0.02 | 0.19 | 0.11 | $2.4 \times 10^{-3}$ | 0.22 |
| | | | 10% | 20% | 0.09 | $2.3 \times 10^{-3}$ | 0.36 | 0.09 | 0.01 | 0.33 | 0.1 | $3.2 \times 10^{-3}$ | 0.4 |
| | | | 20% | 10% | 0.1 | $2.9 \times 10^{-3}$ | 0.38 | 0.11 | $3.3 \times 10^{-3}$ | 0.38 | 0.12 | $4 \times 10^{-3}$ | 0.41 |
| | | | 20% | 20% | 0.1 | $4 \times 10^{-3}$ | 0.52 | 0.07 | $2.6 \times 10^{-3}$ | 0.49 | 0.09 | $2.8 \times 10^{-3}$ | 0.54 |
| 0.2 | 1000 | 10000 | 10% | 10% | 0.11 | $2.6 \times 10^{-3}$ | 0.22 | 0.19 | 0.03 | 0.22 | 0.23 | $6 \times 10^{-3}$ | 0.26 |
| | | | 10% | 20% | 0.16 | $4.9 \times 10^{-3}$ | 0.4 | 0.19 | 0.03 | 0.37 | 0.2 | $7 \times 10^{-3}$ | 0.43 |
| | | | 20% | 10% | 0.17 | $6 \times 10^{-3}$ | 0.41 | 0.2 | $7.5 \times 10^{-3}$ | 0.41 | 0.2 | $8.1 \times 10^{-3}$ | 0.45 |
| | | | 20% | 20% | 0.2 | $9.3 \times 10^{-3}$ | 0.57 | 0.16 | $7.1 \times 10^{-3}$ | 0.54 | 0.14 | $6.1 \times 10^{-3}$ | 0.58 |
| 0.01 | 500 | 20000 | 10% | 10% | 0 | $1.1 \times 10^{-4}$ | 0.23 | 0.01 | $1.1 \times 10^{-4}$ | 0.26 | 0.01 | $2.1 \times 10^{-4}$ | 0.27 |
| | | | 10% | 20% | 0 | $1.1 \times 10^{-4}$ | 0.4 | 0 | $1.1 \times 10^{-4}$ | 0.3 | 0.02 | $6.3 \times 10^{-4}$ | 0.31 |
| | | | 20% | 10% | 0.01 | $2.1 \times 10^{-4}$ | 0.41 | 0 | 0 | 0.33 | 0.01 | $2.1 \times 10^{-4}$ | 0.41 |
| | | | 20% | 20% | 0.01 | $3.2 \times 10^{-4}$ | 0.52 | 0.01 | $2.1 \times 10^{-4}$ | 0.53 | 0 | $1.1 \times 10^{-4}$ | 0.58 |
| 0.05 | 500 | 20000 | 10% | 10% | 0.03 | $6.3 \times 10^{-4}$ | 0.29 | 0.06 | $1.6 \times 10^{-3}$ | 0.3 | 0.06 | $1.7 \times 10^{-3}$ | 0.32 |
| | | | 10% | 20% | 0.04 | $1.1 \times 10^{-3}$ | 0.47 | 0.04 | $1.1 \times 10^{-3}$ | 0.35 | 0.06 | $1.7 \times 10^{-3}$ | 0.37 |
| | | | 20% | 10% | 0.03 | $1.2 \times 10^{-3}$ | 0.5 | 0.06 | $1.6 \times 10^{-3}$ | 0.41 | 0.06 | $1.8 \times 10^{-3}$ | 0.47 |
| | | | 20% | 20% | 0.04 | $1.4 \times 10^{-3}$ | 0.57 | 0.04 | $1.5 \times 10^{-3}$ | 0.59 | 0.06 | $2.4 \times 10^{-3}$ | 0.62 |
| 0.1 | 500 | 20000 | 10% | 10% | 0.05 | $1.4 \times 10^{-3}$ | 0.31 | 0.12 | $3.4 \times 10^{-3}$ | 0.33 | 0.08 | $2.3 \times 10^{-3}$ | 0.35 |
| | | | 10% | 20% | 0.06 | $1.9 \times 10^{-3}$ | 0.5 | 0.11 | $3.1 \times 10^{-3}$ | 0.38 | 0.12 | $3.7 \times 10^{-3}$ | 0.41 |
| | | | 20% | 10% | 0.09 | $2.9 \times 10^{-3}$ | 0.5 | 0.1 | $2.9 \times 10^{-3}$ | 0.46 | 0.11 | $3.7 \times 10^{-3}$ | 0.5 |

Continuation of Table S2

| Continuation of Table S2 | | | | | | | | | | | | | |
|---|---|---|---|---|---|---|---|---|---|---|---|---|---|
| | | | | | PMR-Egger | | | TIGAR | | | PrediXcan | | |
| FDR | $n$ | $N$ | $h_e^2$ | $h_g^2$ | rFDR | FPR | TPR | rFDR | FPR | TPR | rFDR | FPR | TPR |
| | | | 20% | 20% | 0.08 | $3.1 \times 10^{-3}$ | 0.61 | 0.09 | $3.7 \times 10^{-3}$ | 0.62 | 0.12 | $5.6 \times 10^{-3}$ | 0.64 |
| 0.2 | 500 | 20000 | 10% | 10% | 0.15 | $4.4 \times 10^{-3}$ | 0.36 | 0.21 | $7.8 \times 10^{-3}$ | 0.39 | 0.22 | $7.5 \times 10^{-3}$ | 0.39 |
| | | | 10% | 20% | 0.13 | $4.9 \times 10^{-3}$ | 0.53 | 0.22 | $7.9 \times 10^{-3}$ | 0.42 | 0.2 | $7.9 \times 10^{-3}$ | 0.44 |
| | | | 20% | 10% | 0.2 | $8.8 \times 10^{-3}$ | 0.56 | 0.18 | $6.4 \times 10^{-3}$ | 0.5 | 0.18 | $7.3 \times 10^{-3}$ | 0.55 |
| | | | 20% | 20% | 0.16 | $6.8 \times 10^{-3}$ | 0.63 | 0.19 | $9.5 \times 10^{-3}$ | 0.64 | 0.19 | 0.01 | 0.66 |

Note: The notations FDR, FPR, and TPR denote the false discovery rate, false positive rate, and true positive rate, respectively. rFDR is the realized value of the FDR calculated as the number of false positives divided by the sum of false and true positives. The transcriptome data sample size ($n$), GWAS sample size ($N$), gene expression heritability ($h_e^2$) and trait heritability ($h_g^2$) are varied. The results are obtained based on 50 replications in a simulation scenario.

Table S3: Comparison between glossTWAS and other TWAS methods based on the summary of different GReX effect sizes in single-ancestry TWAS.

| | glossTWAS | | | PMR-Egger | | | TIGAR | | | PrediXcan | | |
|---|---|---|---|---|---|---|---|---|---|---|---|---|
| $\beta$ | $\hat{\beta}$ | SD | RMSE | $\hat{\beta}$ | SD | RMSE | $\hat{\beta}$ | SD | RMSE | $\hat{\beta}$ | SD | RMSE |
| 0.5 | 0.45 | 0.08 | 0.12 | 0.45 | 0.12 | 0.15 | 0.34 | 0.04 | 0.19 | 0.45 | 0.05 | 0.15 |
| 0.4 | 0.38 | 0.08 | 0.11 | 0.38 | 0.11 | 0.14 | 0.28 | 0.04 | 0.15 | 0.38 | 0.06 | 0.13 |
| 0.3 | 0.29 | 0.07 | 0.09 | 0.29 | 0.09 | 0.12 | 0.23 | 0.04 | 0.11 | 0.29 | 0.05 | 0.11 |
| 0.2 | 0.20 | 0.07 | 0.08 | 0.20 | 0.09 | 0.11 | 0.17 | 0.04 | 0.08 | 0.21 | 0.05 | 0.08 |
| 0.1 | 0.10 | 0.07 | 0.08 | 0.13 | 0.08 | 0.1 | 0.12 | 0.04 | 0.05 | 0.15 | 0.05 | 0.08 |
| -0.1 | -0.10 | 0.07 | 0.08 | -0.13 | 0.08 | 0.1 | -0.12 | 0.04 | 0.05 | -0.15 | 0.05 | 0.07 |
| -0.2 | -0.19 | 0.07 | 0.09 | -0.20 | 0.09 | 0.11 | -0.17 | 0.04 | 0.08 | -0.21 | 0.05 | 0.08 |
| -0.3 | -0.29 | 0.07 | 0.09 | -0.29 | 0.10 | 0.12 | -0.23 | 0.04 | 0.11 | -0.29 | 0.05 | 0.12 |
| -0.4 | -0.38 | 0.08 | 0.1 | -0.37 | 0.11 | 0.14 | -0.30 | 0.04 | 0.14 | -0.37 | 0.06 | 0.13 |
| -0.5 | -0.46 | 0.09 | 0.12 | -0.45 | 0.12 | 0.15 | -0.36 | 0.04 | 0.19 | -0.46 | 0.05 | 0.16 |

Note: The notations $\beta$, SD, and RMSE denote the true GReX effect size, standard deviation, and root mean square error, respectively. $\hat{\beta}$ is the estimate of $\beta$. The transcriptome data and GWAS sample sizes are $n = 500$ and $N = 10000$. The local heritability of gene expression is fixed at $h_e^2 = 10\%$. The results are obtained based on 300 effective replications in each simulation scenario.

Table S4: Comparison between glossTWAS and other TWAS methods based on relative bias of different GReX effect sizes in single-ancestry TWAS.

| | glossTWAS | | PMR-Egger | | TIGAR | | PrediXcan | |
|---|---|---|---|---|---|---|---|---|
| $\beta$ | mean | SD | mean | SD | mean | SD | mean | SD |
| 0.5 | -10.0 | 14 | -9.5 | 15 | -31.4 | 24 | -10.6 | 30 |
| 0.4 | -5.3 | 17 | -6.0 | 18 | -29.0 | 25 | -5.3 | 34 |
| 0.3 | -3.0 | 19 | -4.1 | 22 | -24.7 | 27 | -4.1 | 33 |
| 0.2 | 0.5 | 29 | 2.2 | 27 | -15.2 | 31 | 6.2 | 37 |
| 0.1 | -0.8 | 45 | 33.4 | 40 | 18.8 | 34 | 52.5 | 56 |
| -0.1 | -4.7 | 43 | 29.8 | 39 | 18.4 | 36 | 49.8 | 51 |
| -0.2 | -3.3 | 29 | 1.3 | 28 | -15.5 | 30 | 5.1 | 36 |
| -0.3 | -2.0 | 20 | -3.3 | 22 | -23.0 | 28 | -2.3 | 39 |
| -0.4 | -6.0 | 16 | -7.2 | 18 | -25.7 | 25 | -6.7 | 32 |
| -0.5 | -7.9 | 14 | -10.4 | 15 | -28.2 | 28 | -8.9 | 35 |

Note: The notations $\beta$ and SD denote the true GReX effect size and standard deviation, respectively. The mean and SD columns for each method denote the average relative bias and the corresponding standard deviation. All the numerical values are represented in percentages. The transcriptome data and GWAS sample sizes are $n = 500$ and $N = 10000$. The local heritability of gene expression is fixed at $h_e^2 = 10\%$. The results are obtained based on 300 effective replications in each simulation scenario.

Table S5: Summary of GReX effect sizes obtained by glossTWAS in multi-ancestry TWAS. Here, eQTLs are not shared between the two ancestries, i.e., $f_0 = 0\%$.

| | | EAS ancestry | | | | EUR ancestry | | | | | |
|---|---|---|---|---|---|---|---|---|---|---|---|
| $\beta_1$ | $\beta_2$ | mean | SD | median | 95% CPI | mean | SD | median | 95% CPI | PPNA | $\log_{10}$(BF) |
| 0.5 | 0.5 | 0.45 | 0.08 | 0.44 | (0.31, 0.60) | 0.45 | 0.08 | 0.44 | (0.31, 0.60) | $2.7 \times 10^{-41}$ | 73.9 |
| 0.4 | 0.4 | 0.37 | 0.07 | 0.37 | (0.25, 0.51) | 0.37 | 0.07 | 0.36 | (0.24, 0.51) | $1.9 \times 10^{-21}$ | 47.4 |
| 0.3 | 0.3 | 0.28 | 0.07 | 0.28 | (0.17, 0.41) | 0.28 | 0.07 | 0.28 | (0.17, 0.41) | $6.9 \times 10^{-11}$ | 24.2 |
| 0.2 | 0.2 | 0.19 | 0.07 | 0.18 | (0.08, 0.31) | 0.19 | 0.06 | 0.19 | (0.08, 0.31) | $9.3 \times 10^{-3}$ | 8.6 |
| 0.1 | 0.1 | 0.09 | 0.06 | 0.08 | (0.01, 0.20) | 0.08 | 0.06 | 0.07 | (0.01, 0.18) | 0.25 | 1.9 |
| 0 | 0 | 0.001 | 0.02 | 0.001 | (-0.004, 0.009) | 0.001 | 0.02 | 0.001 | (-0.003, 0.009) | 0.95 | -0.54 |
| -0.1 | -0.1 | -0.08 | 0.06 | -0.07 | (-0.18, -0.01) | -0.08 | 0.06 | -0.07 | (-0.18, -0.01) | 0.26 | 1.9 |
| -0.2 | -0.2 | -0.19 | 0.06 | -0.18 | (-0.30, -0.09) | -0.19 | 0.06 | -0.18 | (-0.30, -0.08) | $5.5 \times 10^{-3}$ | 9.4 |
| -0.3 | -0.3 | -0.29 | 0.07 | -0.28 | (-0.41, -0.17) | -0.29 | 0.07 | -0.28 | (-0.41, -0.17) | $5.5 \times 10^{-10}$ | 25.1 |
| -0.4 | -0.4 | -0.37 | 0.07 | -0.36 | (-0.51, -0.24) | -0.37 | 0.08 | -0.37 | (-0.52, -0.25) | $1.2 \times 10^{-21}$ | 46.6 |
| -0.5 | -0.5 | -0.45 | 0.08 | -0.44 | (-0.61, -0.31) | -0.45 | 0.08 | -0.44 | (-0.60, -0.31) | $7.5 \times 10^{-40}$ | 75.5 |

Note: $\beta_1$ and $\beta_2$ denote the true GReX effect sizes for the EAS and EUR ancestries, respectively. The abbreviations SD, CPI, PPNA, and BF represent the standard deviation, central posterior interval, posterior probability of null association, and Bayes factor, respectively. The transcriptome data sample sizes are $n_1 = n_2 = 500$, and GWAS data sample sizes are $N_1 = N_2 = 10000$ for the two ancestries. The local heritability of gene expression is fixed at $h_e^2 = 10\%$ in each ancestry. The summaries are obtained based on 300 effective replications in a simulation scenario.

Table S6: Summary of GReX effect sizes obtained by glossTWAS in multi-ancestry TWAS. The effect sizes are unequal and eQTLs are not shared between the two ancestries, i.e., $f_0 = 0\%$.

| | | EAS ancestry | | | | EUR ancestry | | | | | |
|---|---|---|---|---|---|---|---|---|---|---|---|
| $\beta_1$ | $\beta_2$ | mean | SD | median | 95% CPI | mean | SD | median | 95% CPI | PPNA | $\log_{10}$(BF) |
| 0.1 | 0.5 | 0.10 | 0.05 | 0.09 | (0.01, 0.19) | 0.46 | 0.09 | 0.45 | (0.31, 0.63) | $6 \times 10^{-16}$ | 36.1 |
| 0.1 | 0.2 | 0.09 | 0.05 | 0.09 | (0.01, 0.19) | 0.19 | 0.07 | 0.18 | (0.08, 0.31) | 0.06 | 4.9 |
| -0.1 | -0.5 | -0.09 | 0.05 | -0.09 | (-0.18, -0.01) | -0.46 | 0.09 | -0.46 | (-0.63, -0.32) | $1 \times 10^{-13}$ | 38.4 |
| -0.1 | -0.2 | -0.09 | 0.05 | -0.08 | (-0.19, -0.01) | -0.19 | 0.07 | -0.18 | (-0.31, -0.07) | 0.07 | 5 |

Note: $\beta_1$ and $\beta_2$ denote the true GReX effect sizes for the EAS and EUR ancestries, respectively. The abbreviations SD, CPI, PPNA, and BF represent the standard deviation, central posterior interval, posterior probability of null association, and Bayes factor, respectively. The transcriptome data sample sizes are $n_1 = n_2 = 500$, and GWAS data sample sizes are $N_1 = N_2 = 10000$ for the two ancestries. The local heritability of gene expression is fixed at $h_e^2 = 10\%$ in each ancestry. The summaries are obtained based on 300 effective replications in a simulation scenario.

Table S7: Summaries related to realized FDR (rFDR) and true positive rate (TPR) for different levels of Bayesian FDR (BFDR) for glossTWAS in multi-ancestry TWAS.

| $f_0$ | BFDR | $h_e^2$ | $h_g^2$ | rFDR | TPR |
|---|---|---|---|---|---|
| 0% | 0.01 | 10% | 10% | 0 | 0.21 |
| | | 10% | 20% | 0.01 | 0.39 |
| | | 20% | 10% | 0 | 0.37 |
| | | 20% | 20% | 0.01 | 0.53 |
| | | 30% | 30% | 0 | 0.69 |
| | | 40% | 40% | 0 | 0.8 |
| 0% | 0.05 | 10% | 10% | 0.01 | 0.24 |
| | | 10% | 20% | 0.03 | 0.42 |
| | | 20% | 10% | 0.02 | 0.43 |
| | | 20% | 20% | 0.01 | 0.58 |
| | | 30% | 30% | 0.03 | 0.7 |
| | | 40% | 40% | 0.01 | 0.83 |
| 50% | 0.01 | 10% | 10% | 0 | 0.21 |
| | | 10% | 20% | 0 | 0.37 |
| | | 20% | 10% | 0 | 0.37 |
| | | 20% | 20% | 0 | 0.58 |
| | | 30% | 30% | 0 | 0.73 |
| | | 40% | 40% | 0.01 | 0.78 |
| 50% | 0.05 | 10% | 10% | 0.01 | 0.25 |
| | | 10% | 20% | 0.02 | 0.43 |
| | | 20% | 10% | 0.02 | 0.42 |
| | | 20% | 20% | 0.01 | 0.62 |

Continuation of Table S7

| $f_0$ | BFDR | $h_e^2$ | $h_g^2$ | rFDR | TPR |
|---|---|---|---|---|---|
| | | 30% | 30% | 0.02 | 0.75 |
| | | 40% | 40% | 0.02 | 0.8 |

Note: The abbreviations BFDR, FPR, and TPR denote the Bayesian false discovery rate, false positive rate, and true positive rate, respectively. rFDR is the realized value of the BFDR calculated as the number of false positives divided by the sum of false and true positives. The transcriptome data sample sizes are $n_1 = n_2 = 500$ and GWAS sample sizes are $N_1 = N_2 = 10000$. The choices of gene expression heritability ($h_e^2$) and trait heritability ($h_g^2$) are varied. The results are obtained based on 50 replications in a simulation scenario.

Table S8: Summaries related to false discoveries for METRO in multi-ancestry TWAS.

| | | | | METRO | | |
|---|---|---|---|---|---|---|
| $f_0$ | FDR | $h_e^2$ | $h_g^2$ | rFDR | FPR | TPR |
| 0% | 0.01 | 10% | 10% | 0.02 | $1.1 \times 10^{-4}$ | 0.22 |
| | | 10% | 20% | 0 | 0 | 0.37 |
| | | 20% | 10% | 0.01 | $2.1 \times 10^{-4}$ | 0.4 |
| | | 20% | 20% | 0 | 0 | 0.53 |
| 0% | 0.05 | 10% | 10% | 0.03 | $4.2 \times 10^{-4}$ | 0.26 |
| | | 10% | 20% | 0.02 | $5.3 \times 10^{-4}$ | 0.42 |
| | | 20% | 10% | 0.02 | $8.4 \times 10^{-4}$ | 0.43 |
| | | 20% | 20% | 0.03 | $1.1 \times 10^{-3}$ | 0.57 |
| 0% | 0.1 | 10% | 10% | 0.03 | $6.3 \times 10^{-4}$ | 0.28 |
| | | 10% | 20% | 0.06 | $1.6 \times 10^{-3}$ | 0.44 |
| | | 20% | 10% | 0.08 | $2.9 \times 10^{-3}$ | 0.45 |
| | | 20% | 20% | 0.06 | $2.3 \times 10^{-3}$ | 0.59 |
| 0% | 0.2 | 10% | 10% | 0.1 | $2.2 \times 10^{-3}$ | 0.32 |
| | | 10% | 20% | 0.11 | $3.7 \times 10^{-3}$ | 0.49 |
| | | 20% | 10% | 0.12 | $4.7 \times 10^{-3}$ | 0.48 |
| | | 20% | 20% | 0.11 | $5.4 \times 10^{-3}$ | 0.63 |
| 50% | 0.01 | 10% | 10% | 0 | 0 | 0.21 |
| | | 10% | 20% | 0 | 0 | 0.38 |
| | | 20% | 10% | 0 | 0 | 0.43 |
| | | 20% | 20% | 0 | 0 | 0.55 |
| 50% | 0.05 | 10% | 10% | 0.02 | $4.2 \times 10^{-4}$ | 0.27 |
| | | 10% | 20% | 0.01 | $3.2 \times 10^{-4}$ | 0.42 |

Continuation of Table S8

| $f_0$ | FDR | $h_e^2$ | $h_g^2$ | METRO rFDR | METRO FPR | METRO TPR |
|---|---|---|---|---|---|---|
| | | 20% | 10% | 0.04 | $1.1 \times 10^{-3}$ | 0.47 |
| | | 20% | 20% | 0.02 | $7.4 \times 10^{-4}$ | 0.59 |
| 50% | 0.1 | 10% | 10% | 0.04 | $1.2 \times 10^{-3}$ | 0.3 |
| | | 10% | 20% | 0.03 | $9.5 \times 10^{-4}$ | 0.45 |
| | | 20% | 10% | 0.05 | $1.9 \times 10^{-3}$ | 0.5 |
| | | 20% | 20% | 0.04 | $1.6 \times 10^{-3}$ | 0.61 |
| 50% | 0.2 | 10% | 10% | 0.07 | $2.1 \times 10^{-3}$ | 0.32 |
| | | 10% | 20% | 0.1 | $3.1 \times 10^{-3}$ | 0.49 |
| | | 20% | 10% | 0.1 | $3.8 \times 10^{-3}$ | 0.53 |
| | | 20% | 20% | 0.07 | $2.8 \times 10^{-3}$ | 0.63 |

Note: The abbreviations FDR, FPR, and TPR denote the false discovery rate, false positive rate, and true positive rate, respectively. rFDR is the realized value of the FDR calculated as the number of false positives divided by the sum of false and true positives. The transcriptome data sample sizes are $n_1 = n_2 = 500$ and GWAS sample sizes are $N_1 = N_2 = 10000$. The choices of gene expression heritability ($h_e^2$) and trait heritability ($h_g^2$) are varied. The results are obtained based on 50 replications in a simulation scenario.

Table S9: Comparison between glossTWAS and METRO based on the summary of GReX effect sizes in multi-ancestry TWAS. Here, effect sizes are equal and half of the eQTLs are shared between the two ancestries, i.e., $f_0 = 50\%$.

| | | glossTWAS | | | | | | METRO | | | | | |
|---|---|---|---|---|---|---|---|---|---|---|---|---|---|
| | | EAS ancestry | | | EUR ancestry | | | EAS ancestry | | | EUR ancestry | | |
| $\beta_1$ | $\beta_2$ | $\hat{\beta}_1$ | SD | RMSE | $\hat{\beta}_2$ | SD | RMSE | $\hat{\beta}_1$ | SD | RMSE | $\hat{\beta}_2$ | SD | RMSE |
| 0.5 | 0.5 | 0.45 | 0.08 | 0.11 | 0.45 | 0.08 | 0.11 | 0.56 | 0.15 | 0.19 | 0.56 | 0.15 | 0.2 |
| 0.4 | 0.4 | 0.37 | 0.07 | 0.09 | 0.37 | 0.07 | 0.1 | 0.45 | 0.14 | 0.17 | 0.45 | 0.13 | 0.18 |
| 0.3 | 0.3 | 0.29 | 0.06 | 0.08 | 0.29 | 0.06 | 0.08 | 0.36 | 0.13 | 0.17 | 0.35 | 0.12 | 0.16 |
| 0.2 | 0.2 | 0.19 | 0.06 | 0.08 | 0.19 | 0.06 | 0.08 | 0.25 | 0.11 | 0.15 | 0.25 | 0.11 | 0.15 |
| 0.1 | 0.1 | 0.08 | 0.05 | 0.07 | 0.09 | 0.05 | 0.07 | 0.16 | 0.10 | 0.14 | 0.14 | 0.11 | 0.14 |
| -0.1 | -0.1 | -0.09 | 0.05 | 0.07 | -0.09 | 0.05 | 0.07 | -0.15 | 0.10 | 0.14 | -0.14 | 0.10 | 0.13 |
| -0.2 | -0.2 | -0.19 | 0.06 | 0.08 | -0.19 | 0.06 | 0.08 | -0.25 | 0.11 | 0.14 | -0.24 | 0.11 | 0.14 |
| -0.3 | -0.3 | -0.29 | 0.06 | 0.09 | -0.29 | 0.06 | 0.09 | -0.35 | 0.12 | 0.16 | -0.35 | 0.12 | 0.16 |
| -0.4 | -0.4 | -0.38 | 0.07 | 0.09 | -0.37 | 0.07 | 0.09 | -0.47 | 0.13 | 0.19 | -0.46 | 0.14 | 0.18 |
| -0.5 | -0.5 | -0.45 | 0.08 | 0.11 | -0.45 | 0.08 | 0.11 | -0.55 | 0.15 | 0.19 | -0.55 | 0.14 | 0.19 |

Note: $\beta_1$ and $\beta_2$ are true GReX effect sizes for the EAS and EUR ancestries, while $\hat{\beta}_1$ and $\hat{\beta}_2$ are corresponding estimates. The abbreviations SD and RMSE represent the standard deviation and root mean square error, respectively. The transcriptome data sample sizes are $n_1 = n_2 = 500$, and GWAS data sample sizes are $N_1 = N_2 = 10000$ for the two ancestries. The local heritability of gene expression is fixed at $h_e^2 = 10\%$ in each ancestry. The summaries are obtained based on 300 effective replications in a simulation scenario.

Table S10: Comparison between glossTWAS and METRO based on relative bias of GReX effect sizes in multi-ancestry TWAS. Here, effect sizes are equal and half of the eQTLs are shared between the two ancestries, i.e., $f_0 = 50\%$.

| | | glossTWAS | | | | METRO | | | |
|---|---|---|---|---|---|---|---|---|---|
| | | EAS ancestry | | EUR ancestry | | EAS ancestry | | EUR ancestry | |
| $\beta_1$ | $\beta_2$ | mean | SD | mean | SD | mean | SD | mean | SD |
| 0.5 | 0.5 | -10.2 | 13 | -10.3 | 13 | 11.2 | 23 | 12.5 | 23 |
| 0.4 | 0.4 | -7.4 | 14 | -6.9 | 16 | 12.4 | 22 | 13.3 | 24 |
| 0.3 | 0.3 | -2.9 | 19 | -4.0 | 17 | 19.0 | 29 | 17.9 | 27 |
| 0.2 | 0.2 | -6.0 | 29 | -3.9 | 29 | 26.2 | 40 | 24.1 | 39 |
| 0.1 | 0.1 | -18.8 | 39 | -12.8 | 45 | 55.6 | 67 | 41.7 | 76 |
| -0.1 | -0.1 | -12.2 | 44 | -14.8 | 41 | 50.7 | 73 | 37.9 | 71 |
| -0.2 | -0.2 | -5.5 | 25 | -6.5 | 25 | 22.9 | 37 | 19.8 | 36 |
| -0.3 | -0.3 | -4.6 | 18 | -3.6 | 19 | 16.7 | 27 | 16.6 | 28 |
| -0.4 | -0.4 | -5.9 | 15 | -6.3 | 15 | 17.7 | 27 | 14.1 | 25 |
| -0.5 | -0.5 | -10.9 | 12 | -10.3 | 12 | 10.7 | 22 | 9.2 | 23 |

Note: $\beta_1$ and $\beta_2$ are true GReX effect sizes for the EAS and EUR ancestries. The notation SD denotes the standard deviation. The mean and SD columns for each method denote the average relative bias and the corresponding standard deviation. All the numerical values are represented in percentages. The transcriptome data sample sizes are $n_1 = n_2 = 500$, and GWAS data sample sizes are $N_1 = N_2 = 10000$ for the two ancestries. The local heritability of gene expression is fixed at $h_e^2 = 10\%$ in each ancestry. The summaries are obtained based on 300 effective replications in a simulation scenario.

Table S11: Comparison between glossTWAS and METRO based on the summary of GReX effect sizes in multi-ancestry TWAS. Here, effect sizes are unequal and half of the eQTLs are shared between the two ancestries, i.e., $f_0 = 50\%$.

| | | glossTWAS | | | | | | METRO | | | | | |
|---|---|---|---|---|---|---|---|---|---|---|---|---|---|
| | | EAS ancestry | | | EUR ancestry | | | EAS ancestry | | | EUR ancestry | | |
| $\beta_1$ | $\beta_2$ | $\hat{\beta}_1$ | SD | RMSE | $\hat{\beta}_2$ | SD | RMSE | $\hat{\beta}_1$ | SD | RMSE | $\hat{\beta}_2$ | SD | RMSE |
| 0.1 | 0.5 | 0.10 | 0.05 | 0.06 | 0.46 | 0.08 | 0.12 | 0.13 | 0.10 | 0.13 | 0.56 | 0.14 | 0.19 |
| 0.1 | 0.2 | 0.10 | 0.05 | 0.06 | 0.19 | 0.06 | 0.08 | 0.14 | 0.11 | 0.14 | 0.25 | 0.11 | 0.14 |
| -0.1 | -0.5 | -0.10 | 0.05 | 0.06 | -0.46 | 0.08 | 0.12 | -0.13 | 0.10 | 0.13 | -0.56 | 0.14 | 0.19 |
| -0.1 | -0.2 | -0.09 | 0.05 | 0.06 | -0.19 | 0.06 | 0.08 | -0.13 | 0.10 | 0.13 | -0.25 | 0.11 | 0.15 |

Note: $\beta_1$ and $\beta_2$ are true GReX effect sizes for the EAS and EUR ancestries, while $\hat{\beta}_1$ and $\hat{\beta}_2$ are corresponding estimates. The abbreviations SD and RMSE represent the standard deviation and root mean square error, respectively. The transcriptome data sample sizes are $n_1 = n_2 = 500$, and GWAS data sample sizes are $N_1 = N_2 = 10000$ for the two ancestries. The local heritability of gene expression is fixed at $h_e^2 = 10\%$ in each ancestry. The summaries are obtained based on 300 effective replications in a simulation scenario.

Table S12: Comparison between glossTWAS and METRO based on relative bias of GReX effect sizes in multi-ancestry TWAS. Here, effect sizes are unequal and half of the eQTLs are shared between the two ancestries, i.e., $f_0 = 50\%$.

| | | glossTWAS | | | | METRO | | | |
|---|---|---|---|---|---|---|---|---|---|
| | | EAS ancestry | | EUR ancestry | | EAS ancestry | | EUR ancestry | |
| $\beta_1$ | $\beta_2$ | mean | SD | mean | SD | mean | SD | mean | SD |
| 0.1 | 0.5 | 0.7 | 38 | -7.7 | 14 | 26.3 | 75 | 11.8 | 24 |
| 0.1 | 0.2 | -4.4 | 38 | -6.7 | 28 | 36.1 | 69 | 23.6 | 36 |
| -0.1 | -0.5 | -2.2 | 37 | -8.2 | 14 | 26.5 | 71 | 11.2 | 22 |
| -0.1 | -0.2 | -5.3 | 37 | -5.5 | 30 | 33.7 | 73 | 25.4 | 34 |

Note: $\beta_1$ and $\beta_2$ are true GReX effect sizes for the EAS and EUR ancestries. The notation SD denotes the standard deviation. The mean and SD columns for each method denote the average relative bias and the corresponding standard deviation. All the numerical values are represented in percentages. The transcriptome data sample sizes are $n_1 = n_2 = 500$, and GWAS data sample sizes are $N_1 = N_2 = 10000$ for the two ancestries. The local heritability of gene expression is fixed at $h_e^2 = 10\%$ in each ancestry. The summaries are obtained based on 300 effective replications in a simulation scenario.

Table S13: Comparison between glossTWAS and METRO based on the summary of GReX effect sizes in multi-ancestry TWAS. Here, effect sizes are equal and eQTLs are not shared between the two ancestries, i.e., $f_0 = 0\%$.

| | | glossTWAS | | | | | | METRO | | | | | |
|---|---|---|---|---|---|---|---|---|---|---|---|---|---|
| | | EAS ancestry | | | EUR ancestry | | | EAS ancestry | | | EUR ancestry | | |
| $\beta_1$ | $\beta_2$ | $\hat{\beta}_1$ | SD | RMSE | $\hat{\beta}_2$ | SD | RMSE | $\hat{\beta}_1$ | SD | RMSE | $\hat{\beta}_2$ | SD | RMSE |
| 0.5 | 0.5 | 0.45 | 0.08 | 0.12 | 0.45 | 0.08 | 0.11 | 0.49 | 0.16 | 0.21 | 0.50 | 0.16 | 0.19 |
| 0.4 | 0.4 | 0.37 | 0.07 | 0.1 | 0.37 | 0.07 | 0.1 | 0.41 | 0.15 | 0.19 | 0.42 | 0.14 | 0.18 |
| 0.3 | 0.3 | 0.28 | 0.06 | 0.09 | 0.28 | 0.06 | 0.09 | 0.32 | 0.13 | 0.16 | 0.33 | 0.13 | 0.17 |
| 0.2 | 0.2 | 0.19 | 0.06 | 0.09 | 0.19 | 0.06 | 0.08 | 0.23 | 0.13 | 0.15 | 0.22 | 0.12 | 0.15 |
| 0.1 | 0.1 | 0.09 | 0.06 | 0.08 | 0.08 | 0.06 | 0.07 | 0.14 | 0.11 | 0.14 | 0.13 | 0.11 | 0.14 |
| -0.1 | -0.1 | -0.08 | 0.05 | 0.07 | -0.08 | 0.05 | 0.07 | -0.13 | 0.12 | 0.14 | -0.13 | 0.11 | 0.15 |
| -0.2 | -0.2 | -0.19 | 0.06 | 0.08 | -0.19 | 0.06 | 0.08 | -0.23 | 0.13 | 0.16 | -0.22 | 0.13 | 0.16 |
| -0.3 | -0.3 | -0.29 | 0.06 | 0.09 | -0.29 | 0.07 | 0.09 | -0.32 | 0.14 | 0.17 | -0.33 | 0.13 | 0.16 |
| -0.4 | -0.4 | -0.37 | 0.07 | 0.1 | -0.37 | 0.07 | 0.1 | -0.41 | 0.15 | 0.18 | -0.41 | 0.14 | 0.18 |
| -0.5 | -0.5 | -0.45 | 0.08 | 0.11 | -0.45 | 0.08 | 0.11 | -0.51 | 0.16 | 0.21 | -0.49 | 0.15 | 0.19 |

Note: $\beta_1$ and $\beta_2$ are true GReX effect sizes for the EAS and EUR ancestries, while $\hat{\beta}_1$ and $\hat{\beta}_2$ are corresponding estimates. The abbreviations SD and RMSE represent the standard deviation and root mean square error, respectively. The transcriptome data sample sizes are $n_1 = n_2 = 500$, and GWAS data sample sizes are $N_1 = N_2 = 10000$ for the two ancestries. The local heritability of gene expression is fixed at $h_e^2 = 10\%$ in each ancestry. The summaries are obtained based on 300 effective replications in a simulation scenario.

Table S14: Comparison between glossTWAS and METRO based on relative bias of GReX effect sizes in multi-ancestry TWAS. Here, effect sizes are equal and eQTLs are not shared between the two ancestries, i.e., $f_0 = 0\%$.

| | | glossTWAS | | | | METRO | | | |
|---|---|---|---|---|---|---|---|---|---|
| | | EAS ancestry | | EUR ancestry | | EAS ancestry | | EUR ancestry | |
| $\beta_1$ | $\beta_2$ | mean | SD | mean | SD | mean | SD | mean | SD |
| 0.5 | 0.5 | -10.7 | 15 | -11.0 | 13 | -2.5 | 20 | -0.03 | 21 |
| 0.4 | 0.4 | -6.7 | 15 | -7.3 | 16 | 3.6 | 24 | 5.1 | 26 |
| 0.3 | 0.3 | -5.2 | 19 | -5.1 | 19 | 7.1 | 26 | 8.3 | 31 |
| 0.2 | 0.2 | -6.0 | 31 | -4.0 | 29 | 14.7 | 34 | 12.2 | 38 |
| 0.1 | 0.1 | -11.5 | 49 | -19.2 | 40 | 41.8 | 76 | 26.9 | 90 |
| -0.1 | -0.1 | -16.5 | 45 | -22.0 | 42 | 33.6 | 75 | 29.3 | 86 |
| -0.2 | -0.2 | -6.0 | 25 | -7.1 | 26 | 14.0 | 40 | 11.3 | 44 |
| -0.3 | -0.3 | -4.4 | 19 | -4.8 | 19 | 5.7 | 26 | 8.8 | 26 |
| -0.4 | -0.4 | -7.9 | 15 | -6.4 | 15 | 2.4 | 22 | 2.8 | 24 |
| -0.5 | -0.5 | -9.3 | 14 | -10.6 | 13 | 2.3 | 22 | -1.1 | 19 |

Note: $\beta_1$ and $\beta_2$ are true GReX effect sizes for the EAS and EUR ancestries. The notation SD denotes the standard deviation. The mean and SD columns for each method denote the average relative bias and the corresponding standard deviation. All the numerical values are represented in percentages. The transcriptome data sample sizes are $n_1 = n_2 = 500$, and GWAS data sample sizes are $N_1 = N_2 = 10000$ for the two ancestries. The local heritability of gene expression is fixed at $h_e^2 = 10\%$ in each ancestry. The summaries are obtained based on 300 effective replications in a simulation scenario.

Table S15: Comparison between glossTWAS and METRO based on the summary of GReX effect sizes in multi-ancestry TWAS. Here, effect sizes are unequal and eQTLs are not shared between the two ancestries, i.e., $f_0 = 0\%$.

| | | glossTWAS | | | | | | METRO | | | | | |
|---|---|---|---|---|---|---|---|---|---|---|---|---|---|
| | | EAS ancestry | | | EUR ancestry | | | EAS ancestry | | | EUR ancestry | | |
| $\beta_1$ | $\beta_2$ | $\hat{\beta}_1$ | SD | RMSE | $\hat{\beta}_2$ | SD | RMSE | $\hat{\beta}_1$ | SD | RMSE | $\hat{\beta}_2$ | SD | RMSE |
| 0.1 | 0.5 | 0.10 | 0.05 | 0.06 | 0.46 | 0.09 | 0.12 | 0.10 | 0.11 | 0.15 | 0.50 | 0.16 | 0.2 |
| 0.1 | 0.2 | 0.09 | 0.05 | 0.07 | 0.19 | 0.06 | 0.09 | 0.12 | 0.12 | 0.15 | 0.23 | 0.13 | 0.16 |
| -0.1 | -0.5 | -0.09 | 0.05 | 0.06 | -0.46 | 0.08 | 0.12 | -0.10 | 0.12 | 0.15 | -0.51 | 0.14 | 0.18 |
| -0.1 | -0.2 | -0.09 | 0.05 | 0.07 | -0.19 | 0.07 | 0.09 | -0.10 | 0.11 | 0.14 | -0.23 | 0.12 | 0.15 |

Note: $\beta_1$ and $\beta_2$ are true GReX effect sizes for the EAS and EUR ancestries, while $\hat{\beta}_1$ and $\hat{\beta}_2$ are corresponding estimates. The abbreviations SD and RMSE represent the standard deviation and root mean square error, respectively. The transcriptome data sample sizes are $n_1 = n_2 = 500$, and GWAS data sample sizes are $N_1 = N_2 = 10000$ for the two ancestries. The local heritability of gene expression is fixed at $h_e^2 = 10\%$ in each ancestry. The summaries are obtained based on 300 effective replications in a simulation scenario.

Table S16: Comparison between glossTWAS and METRO based on relative bias of GReX effect sizes in multi-ancestry TWAS. Here, effect sizes are unequal and eQTLs are not shared between the two ancestries, i.e., $f_0 = 0\%$.

| | | glossTWAS | | | | METRO | | | |
|---|---|---|---|---|---|---|---|---|---|
| | | EAS ancestry | | EUR ancestry | | EAS ancestry | | EUR ancestry | |
| $\beta_1$ | $\beta_2$ | mean | SD | mean | SD | mean | SD | mean | SD |
| 0.1 | 0.5 | -4.2 | 36 | -8.4 | 16 | 4.4 | 96 | 0.7 | 22 |
| 0.1 | 0.2 | -7.3 | 41 | -6.7 | 32 | 17.2 | 86 | 15.2 | 34 |
| -0.1 | -0.5 | -8.0 | 34 | -7.0 | 15 | 1.6 | 90 | 1.2 | 20 |
| -0.1 | -0.2 | -9.4 | 41 | -5.8 | 31 | 3.9 | 84 | 15.8 | 37 |

Note: $\beta_1$ and $\beta_2$ are true GReX effect sizes for the EAS and EUR ancestries. The notation SD denotes the standard deviation. The mean and SD columns for each method denote the average relative bias and the corresponding standard deviation. All the numerical values are represented in percentages. The transcriptome data sample sizes are $n_1 = n_2 = 500$, and GWAS data sample sizes are $N_1 = N_2 = 10000$ for the two ancestries. The local heritability of gene expression is fixed at $h_e^2 = 10\%$ in each ancestry. The summaries are obtained based on 300 effective replications in a simulation scenario.

Table S17: Results for top 10 associated genes for IOP identified by glossTWAS in single-ancestry TWAS for Europeans at BFDR = 0.01. The prior probability that each null hypothesis is true is fixed at $\pi_0 = 0.95$.

| Chr | Gene | Effect size | SD | PPNA | $\log_{10}$(BF) |
|---|---|---|---|---|---|
| 17 | USP43 | 0.75 | 0.36 | $1.9 \times 10^{-42}$ | 43 |
| 3 | PLD1 | -0.71 | 0.37 | $6 \times 10^{-40}$ | 40.4 |
| 9 | RXRA | -0.92 | 0.5 | $1.1 \times 10^{-29}$ | 30.2 |
| 6 | FOXC1 | -0.76 | 0.32 | $1 \times 10^{-28}$ | 29.3 |
| 3 | TNFSF10 | -0.57 | 0.23 | $7.3 \times 10^{-28}$ | 28.4 |
| 7 | CAV1 | -0.73 | 0.42 | $3.7 \times 10^{-23}$ | 23.7 |
| 9 | WDR5 | -0.6 | 0.25 | $6.2 \times 10^{-23}$ | 23.5 |
| 9 | ZBTB34 | 0.8 | 0.52 | $1.1 \times 10^{-22}$ | 23.2 |
| 9 | ZBTB43 | 0.75 | 0.47 | $3 \times 10^{-22}$ | 22.8 |
| 7 | TES | -0.88 | 0.45 | $6.7 \times 10^{-22}$ | 22.5 |

Note: glossTWAS identifies 339 significant genes at BFDR = 0.01 with $\pi_0 = 0.95$. For brevity, we present the top 10 associated genes. The abbreviations SD, PPNA, and BF represent the standard deviation, posterior probability of null association, and Bayes factor, respectively. The results are obtained based on 7000 MCMC iterations with a burn-in sample size of 2000 and a thinning period of 5.

Table S18: Results for top 10 associated genes for IOP identified by PMR-EGGER in single-ancestry TWAS for Europeans at FDR = 0.01.

| Chr | Gene | Effect size | p-value |
|---|---|---|---|
| 3 | PLD1 | -0.74 | $5 \times 10^{-29}$ |
| 6 | FOXC1 | -0.65 | $1.2 \times 10^{-20}$ |
| 9 | RXRA | -2.20 | $7 \times 10^{-19}$ |
| 7 | TES | 0.90 | $3 \times 10^{-18}$ |
| 3 | TNFSF10 | -1.06 | $2.8 \times 10^{-17}$ |
| 9 | ANGPTL2 | 0.71 | $1.6 \times 10^{-16}$ |
| 7 | ST7 | 1.55 | $1.7 \times 10^{-16}$ |
| 9 | RALGPS1 | -3.20 | $7.4 \times 10^{-16}$ |
| 7 | CAV1 | -0.54 | $1.4 \times 10^{-15}$ |
| 11 | LRP4 | 0.21 | $3.8 \times 10^{-15}$ |

Note: PMR-EGGER identifies 189 significant genes at FDR = 0.01. For brevity, we present the top 10 associated genes.

Table S19: Results for top 10 associated genes for IOP identified by PrediXcan in single-ancestry TWAS for Europeans at FDR = 0.01.

| Chr | Gene | Effect size | SD | p-value |
|---|---|---|---|---|
| 11 | KBTBD4 | 1.53 | 0.19 | $1.4 \times 10^{-15}$ |
| 11 | MADD | 0.19 | 0.03 | $1.1 \times 10^{-13}$ |
| 11 | CEND1 | 0.22 | 0.03 | $6.4 \times 10^{-12}$ |
| 11 | ACP2 | 0.15 | 0.02 | $6.9 \times 10^{-12}$ |
| 11 | MTCH2 | 0.34 | 0.05 | $1.7 \times 10^{-11}$ |
| 1 | ALDH9A1 | 0.19 | 0.03 | $2.2 \times 10^{-10}$ |
| 11 | NR1H3 | 0.12 | 0.02 | $3 \times 10^{-8}$ |
| 1 | LYPLAL1 | 0.29 | 0.06 | $2.3 \times 10^{-6}$ |
| 16 | ZNF276 | 0.15 | 0.03 | $2.3 \times 10^{-6}$ |
| 5 | THBS4 | -0.06 | 0.01 | $2.6 \times 10^{-6}$ |

Note: PrediXcan identifies 16 significant genes at FDR = 0.01. We present the top 10 associated genes.

Table S20: Results for top 10 associated genes for IOP identified by glossTWAS in multi-ancestry TWAS for Europeans and Africans at BFDR = 0.01. The prior probability that each null hypothesis is true is fixed at $\pi_0 = 0.95$.

| | | EUR ancestry | | AFR ancestry | | | |
|---|---|---|---|---|---|---|---|
| Chr | Gene | Effect size | SD | Effect size | SD | PPNA | $\log_{10}$(BF) |
| 17 | USP43 | 0.84 | 0.34 | 0.43 | 0.51 | $1.7 \times 10^{-41}$ | 42.1 |
| 3 | PLD1 | -0.51 | 0.31 | -0.56 | 0.50 | $6.2 \times 10^{-36}$ | 36.5 |
| 6 | FOXC1 | -0.87 | 0.44 | -0.46 | 0.35 | $5.8 \times 10^{-29}$ | 29.5 |
| 9 | RXRA | -0.70 | 0.40 | -0.30 | 0.26 | $5.3 \times 10^{-28}$ | 28.6 |
| 7 | TES | 0.80 | 0.56 | 0.54 | 0.73 | $1.8 \times 10^{-24}$ | 25 |
| 9 | WDR5 | -0.59 | 0.34 | -0.45 | 0.34 | $4.3 \times 10^{-23}$ | 23.6 |
| 7 | CAV1 | -0.61 | 0.54 | -0.31 | 0.33 | $1.2 \times 10^{-22}$ | 23.2 |
| 9 | ZBTB34 | 0.58 | 0.27 | 0.17 | 0.17 | $2.3 \times 10^{-22}$ | 22.9 |
| 3 | NCEH1 | -0.59 | 0.29 | -0.66 | 0.60 | $4 \times 10^{-22}$ | 22.7 |
| 3 | TNFSF10 | -0.45 | 0.18 | -0.52 | 0.31 | $5 \times 10^{-41}$ | 22.6 |

Note: glossTWAS identifies 299 significant genes at BFDR = 0.01 with $\pi_0 = 0.95$. For brevity, we present the top 10 associated genes. The abbreviations SD, PPNA, and BF represent the standard deviation, posterior probability of null association, and Bayes factor, respectively. The results are obtained based on 7000 MCMC iterations with a burn-in sample size of 2000 and a thinning period of 5.

Table S21: Results for top 10 associated genes for IOP identified by METRO in multi-ancestry TWAS for Europeans and Africans at FDR = 0.01.

| Chr | Gene | EUR Effect size | AFR Effect size | p-value |
|---|---|---|---|---|
| 17 | USP43 | -0.29 | 0 | $2.5 \times 10^{-25}$ |
| 3 | PLD1 | -0.40 | -0.47 | $2.2 \times 10^{-24}$ |
| 6 | FOXC1 | 0.62 | 0.64 | $5 \times 10^{-21}$ |
| 7 | TES | -0.48 | -0.12 | $4.8 \times 10^{-20}$ |
| 7 | CAV1 | 0.16 | -0.04 | $1.4 \times 10^{-19}$ |
| 9 | RXRA | 6.12 | -0.9 | $5.6 \times 10^{-19}$ |
| 7 | TFEC | 0.69 | 0.39 | $7.4 \times 10^{-19}$ |
| 9 | ZBTB43 | 0.9 | 0.3 | $8.2 \times 10^{-19}$ |
| 11 | DDB2 | -0.58 | 0.16 | $1.1 \times 10^{-17}$ |
| 11 | KBTBD4 | 0.61 | -0.61 | $1.2 \times 10^{-17}$ |

Note: METRO identifies 304 significant genes at FDR = 0.01. For brevity, we present the top 10 associated genes.
